# Targeted genetic manipulation and yeast-like evolutionary genomics in the green alga *Auxenochlorella*

**DOI:** 10.1101/2025.02.07.637104

**Authors:** Rory J. Craig, Marco A. Dueñas, Dimitrios J. Camacho, Sean D. Gallaher, Maria Clara Avendaño-Monsalve, Yang-Tsung Lin, Crysten E. Blaby-Haas, Jeffrey L. Moseley, Sabeeha S. Merchant

**Affiliations:** California Institute for Quantitative Biosciences (QB3), University of California, Berkeley, CA 94720, USA; School of BioSciences, University of Melbourne, Parkville, VIC 3010, Australia; Department of Plant and Microbial Biology, University of California, Berkeley, CA 94720, USA; Department of Molecular and Cell Biology, University of California, Berkeley, CA 94720, USA; Molecular Foundry, Lawrence Berkeley National Laboratory, Berkeley, CA 94720, USA; US Department of Energy Joint Genome Institute, Lawrence Berkeley National Laboratory, Berkeley, CA 94720, USA; Environmental Genomics and Systems Biology, Lawrence Berkeley National Laboratory, Berkeley, CA 94720, USA

**Keywords:** permuted tRNA, lncRNA, chlorophyll aerobic cyclase, CHL27, reverse genetics, gene targeting, oleaginous, triacylglyceride, GreenCut, epigenetics, synthetic biology

## Abstract

*Auxenochlorella* spp. are diploid oleaginous green algae whose streamlined genomes can be readily manipulated by homologous recombination, making them highly amenable to discovery research and bioengineering. Vegetatively diploid organisms experience specific evolutionary phenomena, including allodiploid hybridization, mitotic recombination, loss-of-heterozygosity and aneuploidy; however, studies of these forces have largely focused on yeasts. Here, we present a telomere-to-telomere phased diploid genome assembly of *Auxenochlorella* UTEX 250-A (haploid length 22 Mb) and introduce a genetic toolkit for site-specific manipulation of the nuclear genome in multiple strains, featuring several selectable markers, inducible promoters, and fluorescent reporters for protein localization. UTEX 250-A is an allodiploid hybrid of *Auxenochlorella protothecoides* and *Auxenochlorella symbiontica*, two species differentiated by extensive chromosomal rearrangements. UTEX 250-A haplotypes are a mosaic of each parental species following mitotic recombination, and two chromosomes are trisomic. Loss-of-heterozygosity events are pervasive across *Auxenochlorella* and can evolve rapidly in the laboratory. High-quality structural annotation yielded ∼7,500 genes per haplotype. *Auxenochlorella* have experienced gene family loss and reduction, including core photosynthesis genes, and exhibit periodic adenine and cytosine methylation at promoters and gene bodies, respectively. Approximately 10% of genes, especially those involved in DNA repair and sex, overlap antisense long noncoding RNAs, which may participate in a regulatory mechanism. We demonstrate the utility of *Auxenochlorella* for fundamental research by knockout of a chlorophyll biosynthesis enzyme, and confirm one trisomy by allele-specific transformation. These results demonstrate the generality of several evolutionary forces associated with vegetative diploidy and provide a foundation for use of *Auxenochlorella* as a reference organism.

**One-sentence summary:** *Auxenochlorella*, green algae shaped by evolutionary forces acting on vegetative diploids, are amenable to discovery research and bioengineering via efficient site-specific homologous recombination

## INTRODUCTION

Green algae present excellent opportunities for discovery in plant biology, especially concerning fundamental molecular and cellular processes. The primary algal model *Chlamydomonas reinhardtii* has been central to research in photosynthesis and chloroplast biogenesis, in addition to functions and processes that are broadly relevant to eukaryotic cells, such as cilia / ciliogenesis and the cell cycle (Sasso et al. 2018; Salome and Merchant 2019). As photosynthetic organisms that can utilize atmospheric carbon dioxide in the production of high value bioproducts, including specialty chemicals and nutraceuticals, green algae also hold great potential in biotechnology (Goold et al. 2024). The ability to edit, add or remove genes is essential to both of these endeavors. Despite the high growth rates and experimental tractability of several algae, the tools to genetically manipulate algal genomes have typically lagged behind those available for many bacteria and yeasts (Sproles et al. 2021). In *C. reinhardtii*, transgenes are typically integrated at untargeted locations via non-homologous end-joining, although recent advances in ribonucleoprotein-mediated approaches have enabled site-specific manipulation (Nievergelt 2025). Non-targeted transgene integration and CRISPR-Cas9 gene editing are also possible in species from industrially relevant trebouxiophyte genera such as *Chlorella* and *Picochlorum* (Yang et al. 2016; Lin and Ng 2020; Krishnan et al. 2025). However, efficient site-specific transformation by homologous recombination, as is commonplace in the budding yeast *Saccharomyces cerevisiae* (Orr-Weaver et al. 1981), is incredibly rare in green algae, having been reported sparingly in species like the ecologically relevant prasinophyte *Ostreococcus tauri* (Lozano et al. 2014).

*Auxenochlorella* is a genus of trebouxiophyte green algae in which homologous recombination is efficient. Site-specific transformation of the nuclear genome in *Auxenochlorella protothecoides*, as well its close relatives from the genus *Prototheca*, has been reported in patent filings (Franklin et al. 2013; Moseley et al. 2024). While *Prototheca* are obligate heterotrophs that have lost the ability to photosynthesize, *Auxenochlorella* can grow robustly as a phototroph, mixotroph or heterotroph, undergoing a metabolic switch that degrades the photosynthetic apparatus when provided with an organic carbon source (Matsuka et al. 1966). Therefore, it is facile to study photosynthetic gene function through reverse genetics, making *Auxenochlorella* an attractive model in plant biology. Both *Auxenochlorella* and *Prototheca* are oleaginous, and *Auxenochlorella* oil and biomass have generally recognized as safe (GRAS) status, favoring their use for engineering of specific lipids for bioproducts and food applications, in addition to biofuels (Franklin et al. 2013; Brooks et al. 2024; Goold et al. 2024). We recently demonstrated the utility of *Auxenochlorella* for discovery research in a study requiring introduction and quantitation of gene expression from several synthetic gene constructs to deduce the mechanism underlying the translation of bicistronic genes (Dueñas et al. 2025a), which are widespread in green algae (Gallaher et al. 2021).

Like budding yeast, *Auxenochlorella* are vegetatively diploid. Microbial organisms that undergo periods of asexual reproduction as diploids are associated with several molecular and evolutionary phenomena. These processes are best studied in *S. cerevisiae* and its close relatives, which undergo a facultatively sexual life cycle featuring long periods of asexual division as a diploid (Tsai et al. 2008; Fischer et al. 2021). Although inter-tetrad outcrossing is relatively rare in yeast, one interesting facet of this life cycle is occasional alloploid hybridization. Mating of haploid gametes from divergent lineages produces an allodiploid hybrid that inherits one set of chromosomes from each parent, whereas allotetraploid hybrids can arise via conjugation of divergent diploid cells or whole-genome duplication of an allodiploid individual (Sipiczki 2008; Gabaldón 2020). Although allodiploid hybrids may be incapable of meiosis due to incompatibilities between the parental chromosomes, they can persist via asexual division (Gabaldón 2020). Indeed, hybrids may exhibit increased fitness or unique phenotypes relative to their parents (i.e., heterosis), and hybrid *Saccharomyces* strains have successfully adapted to anthropomorphic environments such as alcoholic ferments (Sipiczki 2008; Peris et al. 2018; Gallone et al. 2019; Langdon et al. 2019) and olive brine (Pontes et al. 2019). Hybridization has also been linked to the evolution of pathogenicity in taxa including *Candida* yeasts (Pryszcz et al. 2015; Schroder et al. 2016) and *Aspergillus* filamentous fungi (Steenwyk et al. 2020).

Importantly, allodiploid hybridization is not necessarily an evolutionary dead end, since the correct pairing of homologous chromosomes and sexual compatibility can be restored via whole-genome duplication (Charron et al. 2019). Exemplifying this process, an ancient hybridization event is thought to have preceded the whole-genome duplication in the lineage leading to *Saccharomyces* (Marcet-Houben and Gabaldón 2015). Allopolyploidy (hybridization and genome duplication) has also long been recognized as a major force in plant evolution and speciation (Soltis and Soltis 2009).

Although recombination is primarily associated with meiotic cell division, various mitotic recombination processes can occur during asexual growth. In *S. cerevisiae*, mitotic recombination occurs randomly due to double-strand breaks at a rate that is orders of magnitude lower than meiotic recombination (Mancera et al. 2008; Sui et al. 2020). Nonetheless, over lengthy periods of asexuality, mitotic recombination can drastically homogenize genomes via loss-of-heterozygosity (LOH) events, where an allele of one chromosome is replaced by the allele carried by its homologous chromosome (Dutta and Schacherer 2025). LOH can occur at short regions along chromosomes of generally less than 10 kb via gene conversion (i.e., interstitial LOH), or at larger regions that typically extend from an internal location to the telomere (terminal LOH), which are the products of mitotic crossovers or the break-induced replication DNA repair pathway (Sui et al. 2020; Dutta et al. 2021). While LOH can negatively impact fitness (e.g., by unmasking recessive deleterious mutations), it is also an important mechanism by which allelic incompatibilities are purged in hybrids (Smukowski Heil et al. 2017; Lancaster et al. 2019). Furthermore, reciprocal mitotic crossovers result in the shuffling of haplotypes between the parental sub-genomes of hybrids, producing chromosomes that are a mosaic of the ancestral parents (Sipiczki 2008). Mitotic recombination has also been characterized in diploid oomycetes (Dale et al. 2019) and diatoms (Bulankova et al. 2021), suggesting that it is an important evolutionary process across the tree of life.

Another phenomenon associated with asexual reproduction is the emergence of aneuploidy via the loss or gain of copies of specific chromosomes. Unbalanced chromosome numbers have been observed in many eukaryotes and are prevalent in several fungi (Vande Zande et al. 2023). As with hybridization, aneuploidy and its subsequent effects (e.g., gene expression changes) are associated with adaptation to new environments and pathogenicity, although negative fitness effects are also common (Santaguida and Amon 2015). For example, whole or partial aneuploidies are common among clinical isolates of *Candida albicans* (Hirakawa et al. 2015), and an aneuploidy affecting a specific chromosome arm causes antifungal drug resistance (Selmecki et al. 2006). Under laboratory evolution of *C. albicans*, chromosomal duplications have also been observed as transient adaptations to abrupt heat stress (Yona et al. 2012). Transient aneuploidy can result in LOH at the scale of entire chromosomes via duplication of one chromosome copy and loss of the other (Yuen et al. 2007; Andersen et al. 2008).

These drivers of evolution have received scarce attention in chlorophyte algae. Although facultatively sexual, *C. reinhardtii* (class Chlorophyceae) grows asexually as a haploid, with the diploid stage usually limited to a dormant zygospore that obligately divides by meiosis (Harris 2001). Similarly, asexual reproduction in *Chlorella* (Trebouxiophyceae) is haploid, with a possible cryptic uncharacterized sexual cycle (Blanc et al. 2010; Fučiková et al. 2015). Ancient mating type loci have also been identified in haploid *Ostreococcus* species (Mamelliophyceae), demonstrating the presence of haplontic life cycles at the deepest evolutionary branch of the Chlorophyta (Benites et al. 2021). Despite this, several vegetatively diploid species have recently been reported (Figure 1A). Diploid genome assemblies have been assembled for trebouxiophyte algae from the related genera *Nannochloris* (Sanders et al. 2022) and *Picochlorum* (Foflonker et al. 2018; Becker et al. 2020; Barten et al. 2022; da Roza et al. 2024), and for the lichen-forming *Trebouxia lynniae* (Gazquez et al. 2024). *Chloropicon primus* (Chloropicophyceae) is trisomic (three copies) for a single chromosome and is otherwise diploid (Lemieux et al. 2019). In the Chlorophyceae, diploids from the genera *Haematococcus* (Volvocales) and *Tetradesmus* (Sphaeropleales) have been reported (Calhoun et al. 2021; Marcolungo et al. 2024). Most notably, Biondi et al. (2024) described a diploid *Tetradesmus obliquus* strain with extensive heterozygosity but one entirely homozygous chromosome, patterns potentially consistent with hybridization followed by chromosome-scale LOH. These results suggest that the presence of diplontic life cycles may be underestimated in Chlorophyta, although the prevalence of hybridization, LOH and aneuploidy is presently unknown.

**Figure 1.**
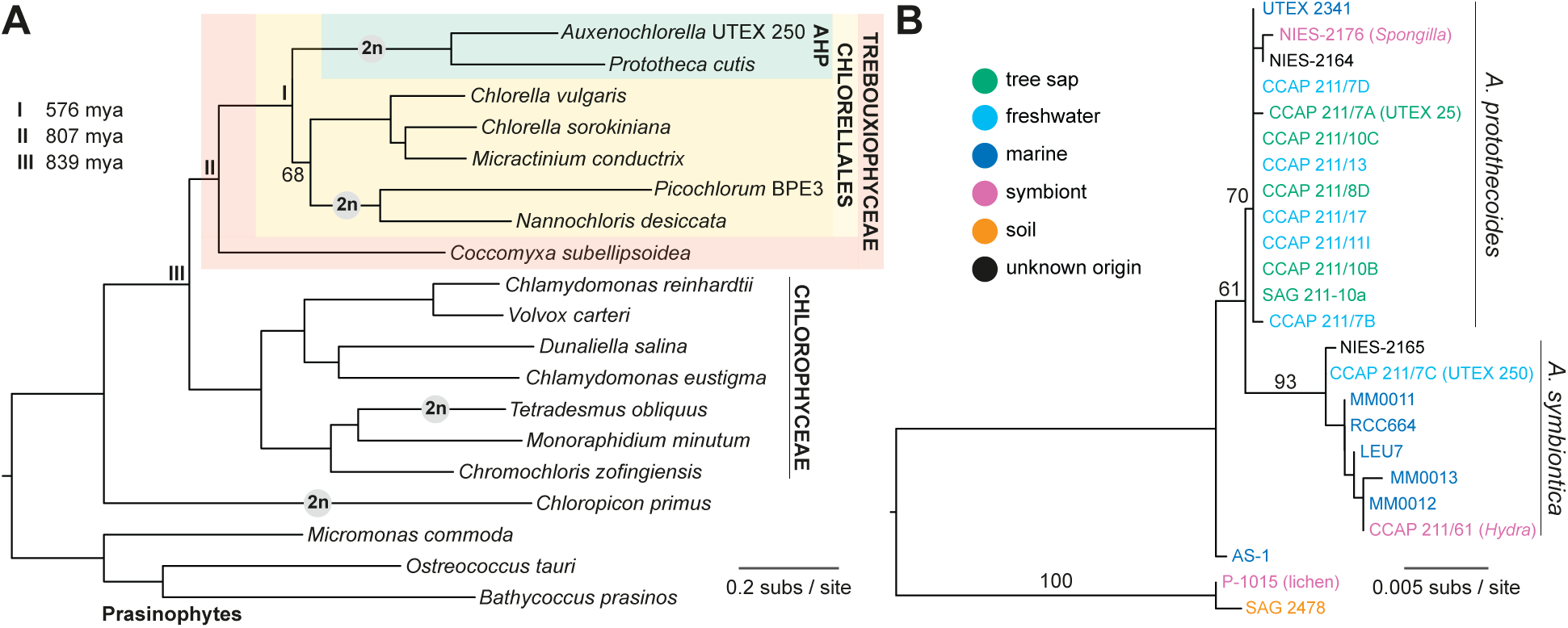
Phylogenetic relationships of *Auxenochlorella*. A) Maximum likelihood phylogeny based on concatenated protein alignment of 669 single-copy orthologs, using the LG+F+R5 substitution model. All nodes received ultrafast bootstrap values >95%, except the node connecting *Picochlorum* and *Nannochloris* to *Chlorella* and *Micractinium* (68% ultrafast bootstrap support). Putative transitions to at least partial vegetative diploidy are shown by the “2n” symbol. Divergence estimates were taken from Del Cortona et al. (2020). B) Maximum likelihood phylogeny based on 18S ribosomal DNA alignment, using the TNe+I substitution model. See Dataset S1 for strain metadata.

Here, we present a fully phased, telomere-to-telomere diploid genome assembly for the *Auxenochlorella* strain UTEX 250-A. We find an unusual genome architecture where six of the twelve chromosome pairs are highly rearranged and non-homologous. We demonstrate that UTEX 250-A is an allodiploid hybrid of *Auxenochlorella protothecoides* and *Auxenochlorella symbiontica*, which each differ in their respective karyotypes. Despite the high heterozygosity introduced by hybridization, several local genomic regions and three entire chromosomes are homozygous following extensive LOH, while two other chromosomes are trisomic. We introduce a highly curated structural annotation of the genome, reporting a streamlined nuclear genome architecture and gene content. Finally, we present a toolkit for genetic manipulation, featuring multiple selectable markers and fluorescent protein expression for intracellular localization studies. We demonstrate application of these tools for reverse genetics via targeted disruption of a gene encoding a chlorophyll biosynthesis enzyme in UTEX 250-A and strains of both parental species, and experimentally verify one UTEX 250-A trisomy by allele-specific targeting of a reporter construct. These resources demonstrate that a green algal genome can be shaped by forces experienced by diplontic taxa like yeasts, and provides a platform for the use of *Auxenochlorella* as a facile reference organism in plant biology and bioengineering.

## RESULTS

### The genus *Auxenochlorella* and existing genomic data

First isolated in the 1890s as “*Chlorella*” *protothecoides* (Krüger 1894b; Krüger 1894a), *Auxenochlorella* was later established as an independent monotypic genus based on physiological, morphological and genetic data (Kalina and Punčochárová 1987; Huss et al. 1999). Indeed, *Auxenochlorella* and true *Chlorella* species likely diverged more than 550 million years ago (Del Cortona et al. 2020) and form distinct lineages within the taxonomically diverse order Chlorellales (Figure 1A). The closest relatives of *Auxenochlorella* have all lost photosynthesis; *Prototheca* is a polyphyletic assemblage of heterotrophic free-living and opportunistically parasitic algae, and *Helicosporidium* are obligate parasites of arthropods (Jagielski et al. 2019). Collectively, these algae form the “AHP lineage” (Ueno et al. 2005), and phylogenetic analyses support multiple independent transitions to heterotrophy, with *Auxenochlorella* enduring as the sole photosynthetic lineage (Figueroa-Martinez et al. 2015; Suzuki et al. 2018; Guo et al. 2022).

Several isolates classified as *A. protothecoides* are maintained in culture centers, including the original strain isolated by W. Krüger from the sap of a poplar tree (UTEX 25 at the University of Texas Culture Collection of Algae). *Auxenochlorella* have also been isolated from fresh and salt water, soil, and as symbionts of freshwater hydrozoans and sponges (Figure 1B). Based on ribosomal DNA (rDNA) analysis, Darienko and Pröschold (2015) described a second species, *A. symbiontica*, with the type strain CCAP 211/61 isolated from *Hydra viridis*. We produced an 18S rDNA phylogeny including some additional isolates (Dataset S1), which supports an association between *A. protothecoides* and both tree sap and freshwater habitats, whereas most *A. symbiontica* strains were isolated from marine environments (Figure 1B). However, the marine strain UTEX 2341 groups with *A. protothecoides* and freshwater symbionts have been isolated for both species, suggesting that both *A. protothecoides* and *A. symbiontica* can live in multiple and overlapping environments.

Gao et al. (2014) produced the first draft assembly of *A. protothecoides* “0710” (=UTEX 25, see below), while additional draft assemblies of UTEX 25 (Vogler et al. 2018) and UTEX 2341 are available at NCBI (Dataset S2). Several draft assemblies of *Prototheca* and *Helicosporidium* have been produced (Pombert et al. 2014; Severgnini et al. 2018; Suzuki et al. 2018; Bakuła et al. 2021; Guo et al. 2022). Although these assemblies are all collapsed to haploids, Guo et al. (2022) reported heterozygosity in two *Prototheca wickerhamii* genomes. Every available AHP lineage genome is highly streamlined relative to the ∼40-60 Mb genomes of *Chlorella* species; for example, the *A. protothecoides* 0710 assembly has a haploid length of 22.9 Mb, with ∼7,000 predicted genes and low repeat content (Gao et al. 2014) (Table 1). It is noteworthy that *Picochlorum* and *Nannochloris* species also possess diploid streamlined genomes (Table 1), and phylogenetic placement of these species as a sister lineage of *Chlorella* (including *Micractinium*) received low bootstrap support in our analysis (Figure 1A). Whether the transition to a diplontic lifecycle occurred independently in the AHP lineage and *Picochlorum*/*Nannochloris*, or whether these lineages may in fact form sister groups, is an outstanding question.

**Table 1.**
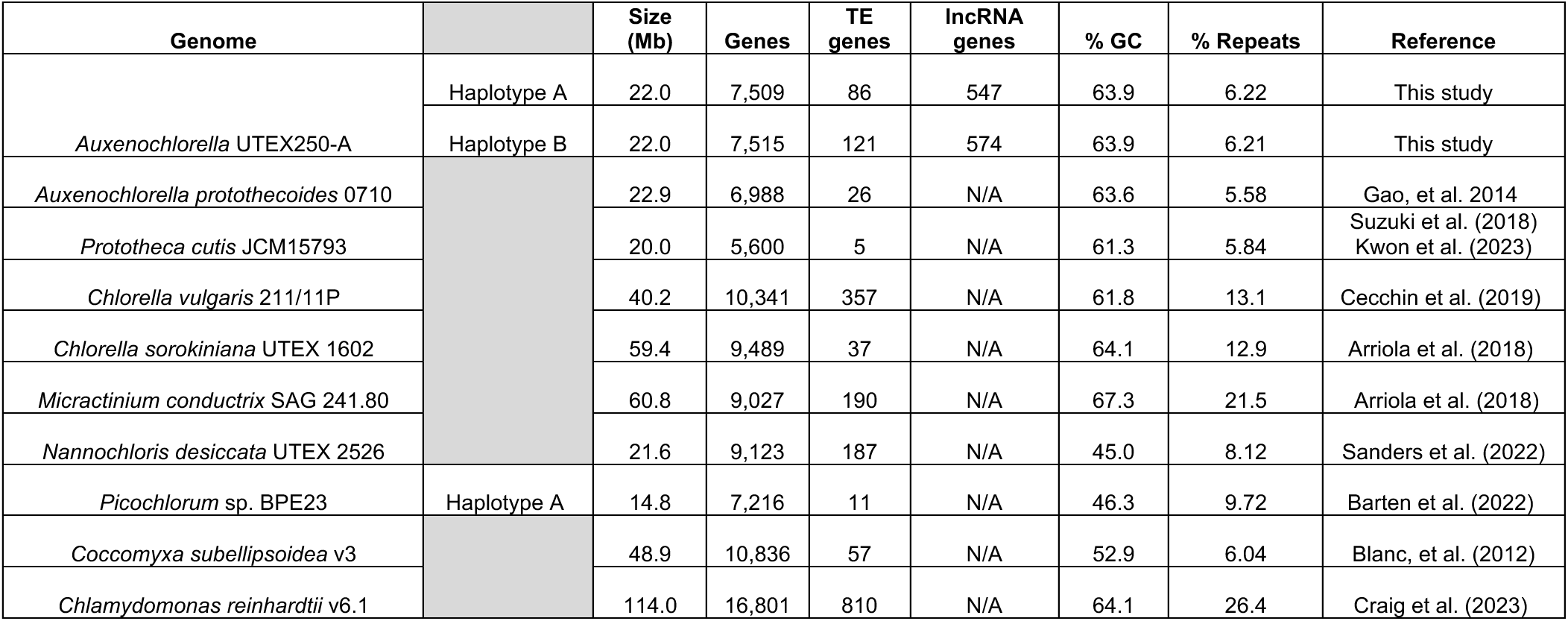
Summary statistics of representative trebouxiophyte genomes with *C. reinhardtii* as an outgroup. See methods for determination of transposable element (TE) genes included in annotations; note that these numbers are not representative of the true number of TE genes in each genome due to differences in annotation methodology. lncRNA = long noncoding RNA. N/A = not applicable.

### Telomere-to-telomere, phased diploid assembly of *Auxenochlorella* UTEX 250-A

To serve as a foundation for discovery research and bioengineering, we sought to produce a reference-quality genome assembly of *Auxenochlorella*. We selected the strain UTEX 250 based on its robust performance in high cell density fermentations (Brooks et al. 2024). Metadata on UTEX 250 are sparse; the strain was isolated before 1950 in the Netherlands by A. J. Kluyver, possibly from freshwater. Although described as *A. protothecoides*, UTEX 250 groups with *A. symbiontica* based on 18S rDNA (Darienko and Pröschold 2015) (Figure 1B). Since our line of the strain has been independently maintained for several years after acquisition from UTEX, we performed single colony purification prior to genome sequencing, and we designate this sub-clone “UTEX 250-A”.

We produced high-coverage Pacific Biosciences (PacBio) HiFi and linked-read OmniC datasets for UTEX 250-A, from which we produced a genome assembly via automated and manual assembly (see Methods). This effort yielded a telomere-to-telomere diploid genome assembly comprising two haplotypes, each featuring 12 gapless nuclear chromosomes spanning ∼22.0 Mb (i.e., collectively 24 nuclear chromosomes spanning 44.0 Mb) (Table 1). Heterozygosity between the two haplotypes is 2.71% (i.e., homologous chromosomal regions differ by ∼27 single nucleotide polymorphisms per 1 kb; heterozygosity calculations exclude entirely homozygous regions, see below). This extensive inter-haplotype variation made it possible to fully phase the assembled chromosomes using the accurate PacBio reads, so that each haplotype corresponds to a physical chromosome in UTEX 250-A. We arbitrarily label these haplotypes “A” and “B”. As presented below, only six of the twelve chromosome pairs exhibit one-to-one homology, and the remaining six are highly rearranged between the two haplotypes.

All chromosomes terminate in the telomeric repeat (TTAGGG)n, except for one chromosome arm per haplotype that terminates in a truncated rDNA array. This telomeric motif differs from the (TTTAGGG)_n_ repeat found in the trebouxiophyte genera *Chlorella* and *Coccomyxa*, which is likely ancestral to green algae, suggesting an independent transition to (TTAGGG)_n_ repeats as observed in some other green algae (Fulnečková et al. 2012). The 5S rDNA genes, which generally form tandem arrays independent of the major rDNA arrays, are present as 16 (haplotype A) or 17 (haplotype B) standalone genes dispersed throughout the genome. Similar non-tandem 5S rDNA genes have been observed in species including the yeasts *Schizosaccharomyces pombe* and *Yarrowia lipolytica* (Torres-Machorro et al. 2010). An initial search for tRNA genes using tRNAscan-SE (Chan et al. 2021) yielded an incomplete set, and we subsequently detected permuted tRNA genes in which the 5’ and 3’ halves of the tRNA gene are organized in reverse (Figure S1). This unusual structure was previously reported in the genomes of the red alga *Cyanidioschyzon merolae* (Soma et al. 2007) and prasinophyte green algae (Maruyama et al. 2010).

As in *A. protothecoides* 0710, repeat content is low, with interspersed and tandem repeats spanning 3.4% and 2.8% of the genome, respectively (Table 1). Among the interspersed repeats, we identified potentially active families of *Metaviridae* long terminal repeat (LTR) retrotransposons, and *hAT* and *EnSpm-Plavaka* cut-and-paste DNA transposons, with many insertions segregating as polymorphisms between the two haplotypes (Figure S2, Dataset S3).

The most abundant repeat is an uncharacterized element that encodes a Fanzor protein, which are RNA-guided endonucleases carried by diverse transposons in eukaryotes and their giant viruses (Saito et al. 2023). Phylogenetically diverse Fanzors are sporadically found in green algal genomes including *C. reinhardtii* (Bao and Jurka 2013) and *Chloropicon primus* (Yoon et al. 2023), and presumably arise via lateral gene transfer.

Green algal centromeres identified thus far are transposon-enriched regions of tens to hundreds of kilobases in species of *Chlorella*, *Coccomyxa* and *Chlamydomonas* (Blanc et al. 2012; Craig et al. 2021; Wang et al. 2024). In line with the paucity of repeats, we found no obvious repeat-rich centromere candidates in UTEX 250-A. We searched for short regional centromeres, nonrepetitive AT-rich regions of 1-5 kb as found in *C. merolae* (Kanesaki et al. 2015) and the diatom *Phaeodactylum tricornutum* (Diner et al. 2017). On 9 of the 12 chromosomes, we identified single AT-rich (mean 48.0% GC, relative to 63.9% GC genome-wide) intergenic regions spanning 2.1 – 7.6 kb (Figure S3, Dataset S4). Although ChIP-sequencing of the centromeric histone variant will be required to confirm centromere locations, considered alongside the low repeat content and non-tandem 5S rDNA genes, these candidates support a highly streamlined architecture of the UTEX 250-A genome.

We produced complete circular genome assemblies for the plastome (84.6 kb, 30.8% GC; Figure S4) and mitogenome (54.0 kb, 29.0% GC; Figure S5). The UTEX 250-A plastome is broadly consistent with existing assemblies from *A. protothecoides* UTEX 25 (Yan et al. 2015; Park et al. 2022). Briefly, the plastome features 78 protein-coding genes and lacks internal repeats, introns, and trans-spliced genes, resulting in a more compact architecture relative to most trebouxiophyte plastomes. The mitogenome features 37 protein-coding genes, including two LAGLIDADG homing endonucleases encoded by introns in *cox1*, as also observed in the *P. wickerhamii* mitogenome (Wolff et al. 1993).

### *Auxenochlorella* UTEX 250 is a putative allodiploid hybrid

Considering the high heterozygosity and rearranged nature of half of the UTEX 250-A diploid chromosomes, we speculated that UTEX 250 may be an allodiploid hybrid between two distinct parental species that possess different karyotypes. Given the close evolutionary relationship between *A. protothecoides* and *A. symbiontica* (Figure 1B), we explored this hypothesis via genome sequencing of strains of both species. We assembled gapless chromosome-level assemblies for *A. protothecoides* strains UTEX 25 and UTEX 2341, and *A. symbiontica* CCAP 211/61, from PacBio HiFi data (supplemented by Oxford Nanopore Technologies, ONT, sequencing for CCAP 211/61; see Methods). The three assemblies were broadly consistent with our UTEX 250-A assembly; each nuclear genome comprises 12 diploid chromosomes, with haploid genome sizes ranging from 21.9 Mb to 23.0 Mb (Dataset S5). However, unlike UTEX 250-A, we observed no rearrangements between the homologous chromosome pairs in each genome, implying a typical diploid karyotype in these strains.

We first considered the six homologous UTEX 250-A chromosomes. Each of these six chromosomes corresponds to a pair of homologous chromosomes in UTEX 25, UTEX 2341, and CCAP 211/61, suggesting that they are ancestral to *Auxenochlorella*. In UTEX 250-A, three of these chromosomes (1, 2 and 6) are mostly heterozygous, whereas the remaining chromosomes (4, 9 and 10) are entirely homozygous (Figure 2A). We presume that the homozygous chromosomes were originally heterozygous, with homozygosity arising following chromosome-scale LOH events (see below). As introduced, heterozygosity in UTEX 250-A is ∼2.7%, although many local genomic regions are homozygous (e.g., the ∼73 kb homozygous region at the left arm terminus of chromosome 1, Figure 2A). In *A. protothecoides*, heterozygosity is substantially lower at 0.59% (UTEX 25) and 0.62% (UTEX 2341) in heterozygous regions, with both genomes also exhibiting extensive regional homozygosity. Average heterozygosity in CCAP 211/61 is 1.35% in heterozygous regions, implying that *A. symbiontica* harbors greater genetic diversity than *A. protothecoides*.

**Figure 2.**
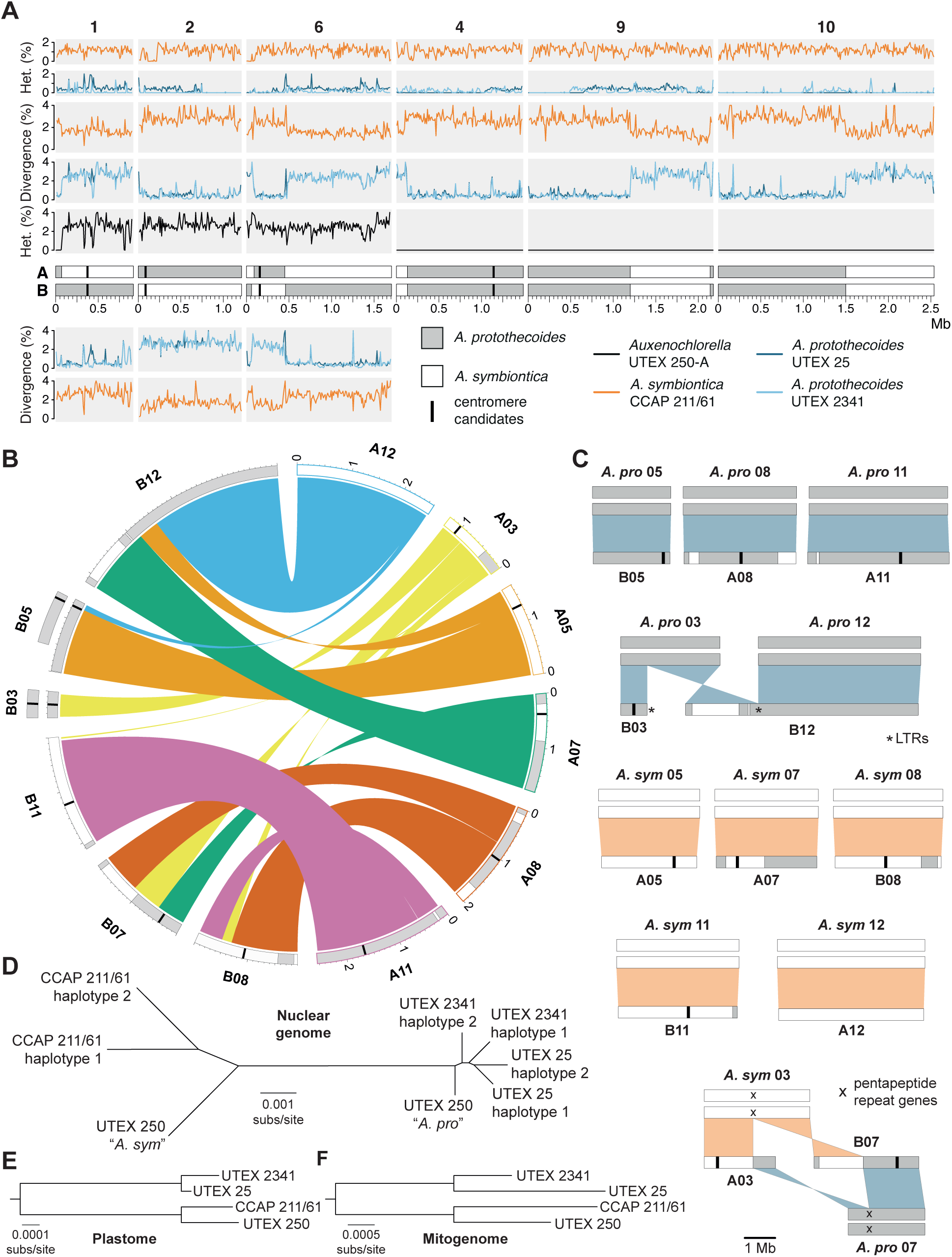
Allodiploid hybrid origin of UTEX 250. A) Heterozygosity and divergence for four *Auxenochlorella* strains in 10 kb windows across the six UTEX 250-A chromosomes with one-to-one homology. For divergence metrics the upper panels refer to haplotype A, lower panels to haplotype B. B) Genomic rearrangements among the remaining six diploid UTEX 250-A chromosomes. C) Comparison between rearranged UTEX 250-A chromosomes and diploid *A. protothecoides* and *A. symbiontica* chromosomes. D) Neighbor joining phylogeny of all *Auxenochlorella* haplotypes (all bootstrap values 100%). E) Neighbor joining phylogeny of plastome sequences (all bootstrap values 100%). F) Neighbor joining phylogeny of mitogenome sequences (all bootstrap values 100%).

When comparing the A and B haplotypes of the UTEX 250-A chromosomes to the other *Auxenochlorella* strains, we observed substantial inter-haplotype variation in genetic divergence (Figure 2A). For example, on chromosome 2, haplotype A differs on average by 0.57% and 0.49% relative to the homologous chromosomes of UTEX 25 and UTEX 2341, respectively.

Meanwhile, haplotype B differs by 2.69% (UTEX 25) and 2.66% (UTEX 2341). Therefore, for chromosome 2, the divergence between haplotype A and the *A. protothecoides* genomes is comparable to heterozygosity between chromosome pairs in *A. protothecoides*, whereas the divergence between haplotype B and *A. protothecoides* is comparable to heterozygosity in UTEX 250-A. The reciprocal pattern is observed via comparison to *A. symbiontica*; divergence between haplotype B and CCAP 211/61 (1.64%) is comparable to heterozygosity in CCAP 211/61, whereas divergence between haplotype A and CCAP 211/61 (2.84%) is comparable to UTEX 250-A heterozygosity. These results are consistent with haplotype A of chromosome 2 originating from an *A. protothecoides*-like ancestor, and haplotype B originating from an *A. symbiontica*-like ancestor. Extending this logic, it is possible to use genetic divergence to assign a putative parent of origin to all genomic regions in UTEX 250-A (Figure 2A, B).

In contrast to chromosome 2, we observed intra-haplotype switches in assigned parentage on the remaining five homologous chromosome pairs (Figure 2A). On chromosome 1, a single switch corresponds to the 73 kb homozygous region at the left arm terminus, and can be attributed to a LOH event in which the ancestral haplotype A copy of this region was replaced by haplotype B, resulting in both resembling *A. protothecoides*. Conversely, the switches on chromosome 6 are reciprocal, resulting in a ∼360 kb region of *A. protothecoides*-like sequence on an otherwise *A. symbiontica*-like chromosome for haplotype A (and vice versa on haplotype B). This mosaicism could be explained by mitotic crossover following hybridization, i.e., each haplotype originally corresponded to one parent (as for chromosome 2), but has subsequently undergone reciprocal exchange. The homozygous chromosomes 4, 9 and 10 also exhibit switches in assigned parentage that are consistent with either LOH or crossover events occurring in a heterozygous state that preceded complete LOH. Note that the A and B labels are arbitrary, and the genomic mosaicism makes it impossible to correspond physical haplotypes to parents of origin in UTEX 250-A.

We next tested whether the UTEX 250-A rearranged chromosomes correspond to the ancestral karyotypes of *A. protothecoides* and *A. symbiontica*. The extent of the rearrangements among these chromosomes can only be explained by multiple translocations, one fusion, and one fission (Figure 2B). Remarkably, eight of the twelve chromosome copies correspond exactly to a homologous chromosome pair in either *A. protothecoides* (both UTEX 25 and UTEX 2341, which have an identical karyotype) or *A. symbiontica* (CCAP 211/61). Specifically, UTEX 250-A chromosomes B05 (i.e., chromosome 5, haplotype B), A08 and A11 correspond to homologous chromosome pairs in *A. protothecoides*, whereas A05, A07, B08, B11 and A12 correspond to homologous chromosome pairs in *A. symbiontica* (Figure 2C). Therefore, most of the rearrangements in UTEX 250-A can be explained by direct inheritance from parental species with different karyotypes. The remaining chromosomal rearrangements can be explained by events that occurred following hybridization. The UTEX 250-A chromosomes B03 and B12 correspond to a pair of *A. protothecoides* chromosomes, suggesting fission of the ancestral *A. protothecoides* chromosome 3 and subsequent fusion of one of the resulting chromosomal fragments (which is predicted to have lacked a centromere) to the ancestral *A. protothecoides* chromosome 12 (Figure 2C). Interestingly, LTR elements are present at both the *de novo* terminus of B03 and the fusion site of B12, suggesting that retrotransposons may have invaded (and potentially stabilized) the unprotected chromosomal ends following fission. Finally, the UTEX 250-A chromosomes A03 and B07 are consistent with a reciprocal translocation between the ancestral *A. protothecoides* chromosome 7 and *A. symbiontica* chromosome 3 (Figure 2C). These ancestral chromosomes feature a common locus of ∼6 kb that corresponds to two genes that are part of the largest *Auxenochlorella*-specific gene family (see below), suggesting that the translocation occurred via ectopic recombination between these nonhomologous repeats. It is also noteworthy that switches in the assigned parent of origin are frequently observed on the rearranged chromosomes (Figure 2B, C), implying that mitotic recombination has occurred between homologous regions despite the rearrangements.

Finally, we constructed a phylogeny based on genomic regions that i) are present on UTEX 250-A chromosomes that have not obviously experienced mitotic crossovers, and ii) are not affected by LOH in any of the four strains. As expected, this analysis clearly divided the UTEX 250-A genome into regions that clustered with either UTEX 25 and UTEX 2341 (*A. protothecoides-* like) or CCAP 211/61 (*A. symbiontica*-like), with the longer branch lengths for *A. symbiontica* reflecting the higher genetic diversity in the species (Figure 2D). We also constructed trees for the plastome (Figure 2E) and mitogenome (Figure 2F), which demonstrated putative inheritance from *A. symbiontica* in both cases.

Overall, *A. protothecoides* and *A. symbiontica* represent authentic species that are ∼2.7% divergent at the sequence-level and are distinguished at the chromosome-level by multiple rearrangements. Assuming that the sequenced strains are representative, genetic diversity may be more than two-fold higher in *A. symbiontica* than *A. protothecoides*. UTEX 250 is a putative allodiploid hybrid that presumably originated via the fusion of haploid cells of *A. symbiontica* and *A. protothecoides*. The ancestrally inherited haplotypes have subsequently been mixed via nonreciprocal and reciprocal mitotic exchange, and four ancestrally inherited chromosomes have been involved in post-hybridization rearrangements. Given the efficiency of homologous recombination in *Auxenochlorella* (see below), it is possible that the rapid emergence of chromosomal differences between the closely related *A. protothecoides* and *A. symbiontica* may have been driven by ectopic recombination events. Indeed, assuming a sexual life cycle for *Auxenochlorella*, UTEX 250-A is presumably incapable of meiosis due to the non-homologous nature of its chromosomes (Garagna et al. 2014), and the karyotypic variation between the two species may present a barrier to gene flow.

### Pervasive loss-of-heterozygosity and evolution in the laboratory

LOH events are associated with several mechanisms and can affect local genomic regions (interstitial LOH), large regions that extend to a chromosome terminus (terminal LOH), or entire chromosomes. We observed all three LOH categories in UTEX 250-A, resulting in at least 35.5% of the genome being homozygous (conservatively assuming a minimum interstitial LOH length of 1 kb). The three entirely homozygous chromosomes, 4, 9 and 10, collectively span 28.2% of the genome. Terminal LOH events affect seven chromosomal termini (e.g., the left arm terminus of chromosome A01, Figure 2A), ranging from ∼12 kb to ∼271 kb (mean 111 kb) and collectively spanning 3.5% of the genome. We observed 278 putative interstitial LOH events, 36 of which exceed 5 kb in length, reaching a maximum of 37.1 kb. Notably, LOH is pervasive in all of the sequenced *Auxenochlorella* genomes; 23.3% of the *A. symbiontica* CCAP 211/61 genome is homozygous, whereas levels of homozygosity reach 67.2% (UTEX 25) and 80.3% (UTEX 2341) in *A. protothecoides*.

Since all the studied *Auxenochlorella* strains have been in culture for decades, we questioned whether LOH could have accumulated in the laboratory. UTEX 25 is one of the oldest microbes in culture, having been isolated in the late 19^th^ century (Krüger 1894b; Krüger 1894a). CCAP 211/61 was isolated in 1982, and UTEX 2341 prior to 1984 (Seto et al. 1984). Conveniently, many of these strains are maintained at different culture centers, including UTEX and CCAP (Culture Collection of Algae and Protozoa). Although records are uncertain, a sister lineage of UTEX 250 has likely been independently maintained for more than 70 years as CCAP 211/7C (Figure 3A). To test whether LOH has occurred during culture, we obtained CCAP 211/7C and performed high-coverage ONT sequencing.

**Figure 3.**
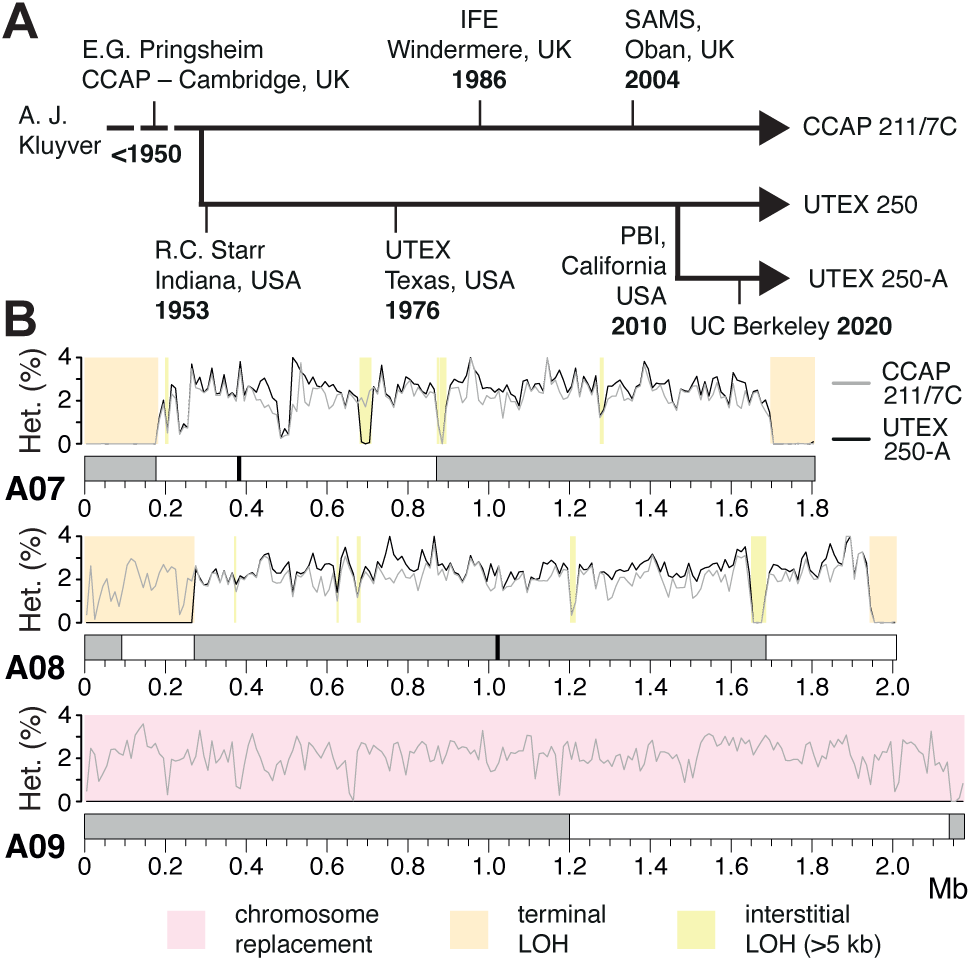
Loss-of-heterozygosity and evolution in the laboratory. A) Putative strain history of UTEX 250. IFE = Institute of Freshwater Ecology, SAMS = Scottish Association for Marine Science, PBI = Phycoil Biotechnology International, Inc. B) Heterozygosity in 10 kb windows across three example chromosomes. Chromosomes are shaded according to parental species of origin and candidate centromeres are marked where available. LOH = loss-of-heterozygosity. Different LOH categories in UTEX 250-A are shown by colored boxes.

Remarkably, LOH events affect only 19.7% of the CCAP 211/7C genome. Although chromosome 10 is entirely homozygous in both UTEX 250-A and CCAP 211/7C, chromosomes 4 and 9 are heterozygous in CCAP 211/7C (Figure 3B, S6). Thus, two of the chromosome-scale LOH events must have occurred during culture at UTEX or in our laboratory. Similarly, of the seven UTEX 250-A terminal LOH events, only five are present in CCAP 211/7C (Figure S6).

For example, both terminal LOH events on chromosome A07 and the right arm terminal LOH event on A08 are shared, whereas the event on the left arm terminus of A08 is unique to UTEX 250-A (Figure 3B). The same pattern is observed for interstitial LOH, for example, the largest interstitial event in the genome (37.1 kb at ∼1.66 Mb on chromosome A08) is shared, whereas the second largest (29.2 kb, ∼0.69 Mb on A07) is unique to UTEX 250-A (Figure 3B).

Interestingly, almost all of the strain-specific LOH events are unique to UTEX 250-A, suggesting that the frequency of LOH during culture at CCAP is substantially lower.

These results support the rapid accumulation of LOH events under at least some laboratory conditions, as has been observed experimentally in yeast (Sui et al. 2020; Dutta et al. 2021). Contrastingly, the UTEX 250-A chromosomal rearrangements that likely followed hybridization are also present in CCAP 211/7C (Figure S6), suggesting that these events occurred prior to sampling of the original strain (or at least before the UTEX and CCAP lineages were established). Although it is possible that some of the LOH events in UTEX 250-A and CCAP 211/7C were fixed by positive selection to overcome post-hybridization incompatibles, there is no obvious directionality with respect to parental species: approximately equal numbers of bases within regions affected by interstitial and terminal LOH events can be assigned to *A. protothecoides* and *A. symbiontica*. Furthermore, LOH is even more prominent in the non-hybrid *A. protothecoides* strains, suggesting that it is a widespread phenomenon in *Auxenochlorella*. It is possible that the different frequencies of LOH events in CCAP 211/7C and UTEX 250-A could be attributed to differences in culture conditions (e.g., liquid vs agar cultures that would affect clonal population sizes). We recommend cryopreservation and avoiding bottlenecks in culture sizes to avoid the fixation of LOH events by drift.

### Aneuploidy is widespread in *Auxenochlorella*

While assembling the UTEX250-A genome, we noticed that two genomic regions exhibited unexpectedly high read coverage. Specifically, when mapping the PacBio reads against the diploid assembly, the entirety of chromosome B03 and a 932 kb fragment of B05 exhibit median read coverages of 528x and 519x, respectively, approximately twice the 272x coverage of the other individual chromosomes, suggesting that these regions are trisomic when considering both A and B haplotypes (Figure 4A). The putatively duplicated part of B05 features the right arm terminus and associated telomere, and we identified reads featuring telomeric repeats that map internally to B05 precisely at the location where coverage doubles (Figure S7). Conceivably, B05 may have duplicated and then undergone fission, with the fragment retaining the centromere acquiring a de novo telomere at its left terminus, yielding a partly aneuploid chromosome. We compared read coverage between UTEX 250-A and CCAP 211/7C to explore whether the observed aneuploidies could have evolved during laboratory culture. Surprisingly, B05 exhibits normal coverage in the CCAP 211/7C ONT read dataset, whereas B03 has three times the expected coverage (163x relative to 53x genome-wide, Figure 4B), suggesting tetrasomy for this chromosome (three copies of B03 and one copy of the homologous region on A03; Figure S6).

**Figure 4.**
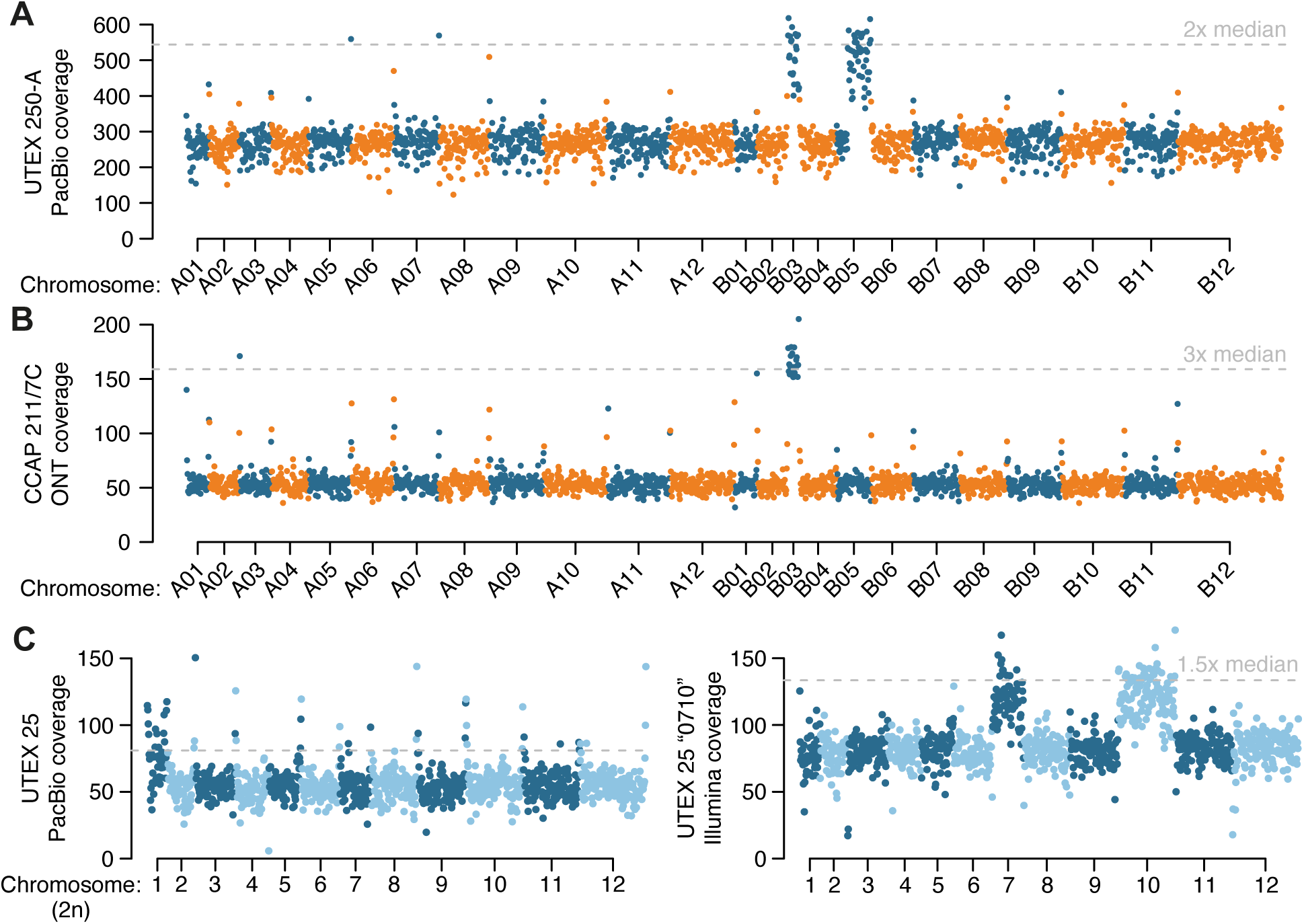
Aneuploidy in *Auxenochlorella.* A) UTEX 250-A PacBio read coverage for 20-kb windows across the phased diploid genome assembly of UTEX 250-A. B) CCAP 211/7C ONT read coverage for 20-kb windows across the phased diploid genome assembly of UTEX 250-A. C) UTEX 25 PacBio read coverage and strain “0710” Illumina read coverage for 20-kb windows across a representative haplotype of the UTEX 25 genome assembly.

Thus, the B05 duplication appears to have occurred during culture leading to UTEX 250-A (or alternatively CCAP 211/7C has reverted to disomy), whereas aneuploidy of B03 may have existed prior to sampling, with subsequent copy number change during culture.

We next checked for possible aneuploidies in *A. protothecoides* UTEX 25 and UTEX 2341, and *A. symbiontica* CCAP 211/61, by mapping PacBio reads for each strain against one representative haplotype from their respective genome assembly. For UTEX 2341 and CCAP 211/61, coverage estimates were consistent across all chromosomes, suggesting euploidy (Figure S8). However, median read coverage for chromosome 1 (76x) of UTEX 25 was approximately 50% higher than median coverage across the rest of the genome (54x), implying trisomy (Figure 4C). We also noted that the existing *A. protothecoides* “0710” genome assembly exhibited almost no variation relative to our UTEX 25 assembly, indicating that they are replicates of the same strain. Surprisingly, when mapping the “0710” Illumina genomic reads to our UTEX 25 assembly we found no increase in coverage for chromosome 1, whereas coverage for chromosomes 7 and 10 (131 and 134x, respectively) were ∼50% higher than the median of 89x across the rest of the genome (Figure 4C). Thus, it appears that the evolution of aneuploidy is dynamic, occurring on short timescales of laboratory culture, at least in the UTEX 250 and UTEX 25 genetic backgrounds. These results are in line with chromosome duplication events observed in several haploid green algal species during mutation accumulation experiments (Krasovec et al. 2023; Lopez-Cortegano et al. 2023). Transient aneuploidy may explain the rapid emergence of whole-chromosome LOH, suggesting that it is a prominent evolutionary force in *Auxenochlorella*. It is unclear whether any of the observed aneuploidies confer a selective advantage; at least one additional copy of B03 has been maintained over decades of culture in UTEX 250 and CCAP 211/7C, potentially indicating that it could be beneficial. There are 149 genes on B03, and analyses focusing on the effects of dosage on gene and protein expression will be required to address this.

### Gene family reduction and loss contribute to the streamlined *Auxenochlorella* genome

To facilitate the development of *Auxenochlorella* as a model system, we aimed to produce a reference quality structural annotation for the UTEX 250-A genome. We sequenced PacBio cDNA (IsoSeq) libraries from UTEX 250-A grown either autotrophically or with glucose, and multiple Illumina RNAseq libraries from different growth conditions. Utilizing these datasets as evidence, we annotated gene models independently for the A and B haplotypes using a combination of gene predictors and extensive manual curation (see Methods). This yielded 7,509 and 7,515 protein-coding genes on the A and B haplotypes, respectively, with an additional 86 and 121 genes carried by transposable elements included in the annotation (Table 1). More than two thirds of the gene models are derived from multiple full-length IsoSeq reads, enabling high confidence identification of transcription start sites (TSS), terminators, and alternative splice variants. Approximately 43% of genes are annotated with at least one alternative transcript, totaling 13,125 and 13,345 transcripts on the A and B haplotypes, respectively. Considering the A haplotype, the majority (56%) of alternative isoforms differ only in their UTR sequences (e.g., due to alternative TSSs or terminators) relative to another isoform of the same gene. For alternative isoforms that alter the predicted protein sequence, 31% feature a retained intron and 18% an exon with an alternate 5’ or 3’ splice junction. The remaining 51% were classified as complex isoforms, many of which are predicted to originate from internal promoters that produce truncated proteins (see below). Alternative transcripts that skip exons are rare (<1%). Finally, 612 gene models were corrected based on manual curation, including 43 bicistronic loci analyzed by Dueñas et al. (2025a), which are frequently misannotated by automated approaches (Gallaher et al. 2021). Speaking to the compactness of the UTEX 250-A genome, 74% of genes encode a protein with a Pfam domain (Dataset S6), a metric that is typically 25-50% in algal genomes (Blaby-Haas and Merchant 2019). Via orthology to *C. reinhardtii* proteins, one-third of genes were also assigned formal gene symbols (Dataset S7), which generally correspond to at least some level of functional curation (Blaby et al. 2014; Craig et al. 2023).

Via synteny analysis, we were able to match ∼98% of genes as allelic pairs between the A and B haplotypes. Genes were given a unique ID featuring the haplotype and chromosome (e.g., A11, B08) and a five-digit identifier that is shared between allelic pairs, e.g., despite being located on rearranged chromosomes, UTEX250_A11.30955 and UTEX250_B08.30955 refer to alleles of the same gene. As expected, heterozygosity between the A and B alleles at four-fold degenerate sites in coding sequence is substantially higher (3.44%) than at zero-fold degenerate sites (0.84%), although this nonetheless corresponds to more than 35,000 allelic variants that result in amino acid differences. Heterozygosity is highest in intergenic (3.51%) and intronic (4.02%) sites (Dataset S8). Only ∼15% of the haplotype-specific genes encode a predicted protein with a Pfam domain (Dataset S6), and many of these genes may be evolutionarily young or misannotated long noncoding RNA (lncRNA) genes that carry a spurious open reading frame (ORF). Some haplotype-specific genes are the result of structural variation between the A and B haplotypes, including a few copy number and presence/absence variants.

Although we identified over 500 additional genes in our UTEX 250-A genome relative to the existing *A. protothecoides* 0710 assembly, the *Auxenochlorella* gene number remains lower than that of most trebouxiophyte species, and substantially lower than that of chlorophyceaen species such as *C. reinhardtii* (Table 1). We performed an orthology analysis of UTEX 250-A proteins against the best quality chlorophyte annotations, arbitrarily using the A haplotype genes supplemented with genes unique to the B haplotype. Figure 1A presents a phylogeny based on 669 single-copy orthologs identified by this analysis, and Figure 5A shows the intersecting sets of orthogroups among Chlorellales species, with *C. subellipsoidea* and *C. reinhardtii* as outgroups. Ignoring species-specific orthogroups, the largest set corresponds to gene families present in all species (2,974 orthogroups), many of which are presumably essential core-

**Figure 5.**
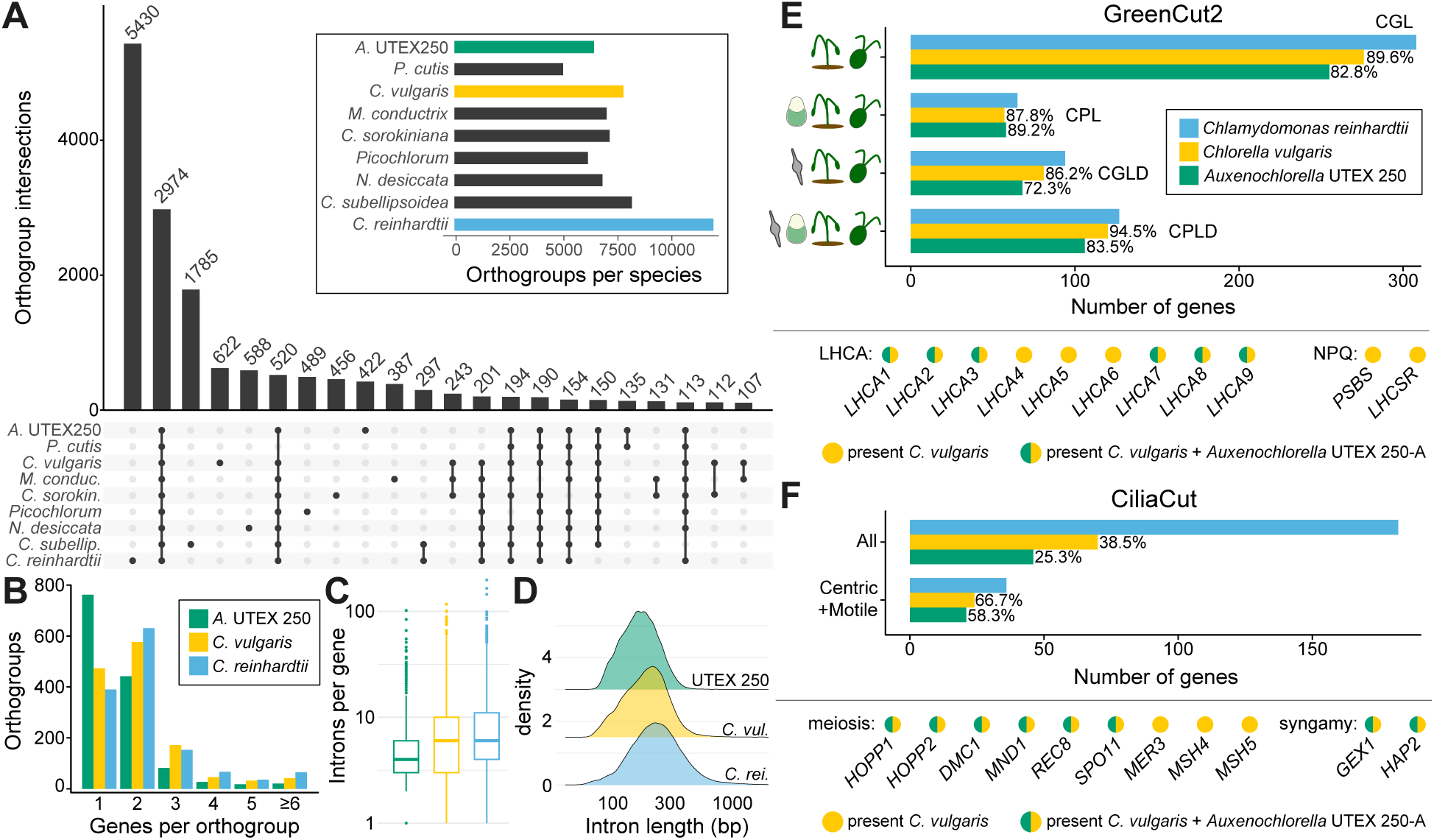
Gene family loss and reduction in *Auxenochlorella* UTEX 250-A. A) Upset plot showing intersection between the presence and absence of orthogroups among Trebouxiophyte algae, with *C. reinhardtii* as an outgroup. Only intersects with more than 100 orthogroups are shown. B) Number of genes per orthogroup for orthogroups that are present in each of *Auxenochlorella* UTEX 250-A, *C. vulgaris* and *C. reinhardtii*, and have more than one gene in at least one of the species. C) Introns per gene for multi-exonic genes. D) Intron length distributions. E) Presence of GreenCut2 genes in UTEX 250-A and *C. vulgaris* relative to *C. reinhardtii*. Light-harvesting complex genes of the photosystem I antenna (*LHCA*) and genes functioning in nonphotochemical quenching (NPQ) are highlighted; *PSBS* and *LHCSR* are present as two and three paralogous genes in *C. reinhardtii* and single orthologs in *C. vulgaris* (see Dataset S13). The CGL, CPL, CGLD and CPLD divisions are defined in the main text. F) Presence of CiliaCut, meiosis and syngamy genes in UTEX 250-A and *C. vulgaris* relative to *C. reinhardtii*.

Chlorophyte gene families (Dataset S9). The second largest set features gene families specifically absent in the non-photosynthetic *P. cutis* (520 orthogroups), most of which are expected to have functions related to photosynthesis (Suzuki et al. 2018). With respect to reduced gene content in *Auxenochlorella*, 297 orthogroups are absent across all Chlorellales species, and 201 orthogroups are specifically absent in *Auxenochlorella* UTEX 250-A and *P. cutis* but present in all other species.

For more detailed analyses, we compared the *Auxenochlorella* UTEX 250-A annotation to the high-quality gene models of *Chlorella vulgaris* (Cecchin et al. 2019) and *C. reinhardtii* (Craig et al. 2023), excluding any genes carried by transposable elements. Considering gene family loss, there are 782 orthogroups that are absent from UTEX 250-A but present in both other species, featuring 1,025 *C. vulgaris* genes and 1,212 *C. reinhardtii* genes. Considering gene family size, 1,341 orthogroups are present in all three species and have more than one gene in at least one species, featuring 2,228 UTEX 250-A genes, 2,915 *C. vulgaris* genes, and 3,272 *C. reinhardtii* genes. Approximately 56% of these orthogroups feature a single UTEX 250-A gene, relative to 35% and 29% for *C. vulgaris* and *C. reinhardtii*, respectively (Figure 5B). UTEX 250-A genes also contain fewer and shorter introns (mean 5.1 introns per gene, mean length 179 bp) relative to *C. vulgaris* (means 7.3 and 206 bp) and *C. reinhardtii* (means 7.8 and 272 bp) (Figure 5C, D). Thus, the streamlined *Auxenochlorella* UTEX 250-A genome can be explained by both complete loss or lower complexity of gene families, alongside lower repeat content (Table 1) and the presence of fewer and shorter introns.

Many of the lost genes can be linked to known metabolic pathways. Two distinguishing features of *Auxenochlorella* are an absolute requirement for thiamine and the inability to grow on nitrate as a N source (Kalina and Punčochárová 1987). As expected, the nitrate reductase gene *NIA1* and *THIC1*, encoding 4-amino-5-hydroxymethyl-2-methylpyrimidine phosphate synthase necessary for thiamine biosynthesis, are among the genes present in *C. vulgaris* and *C. reinhardtii* but absent in UTEX 250-A. All other components of the thiamine biosynthesis pathway are present in UTEX 250-A, enabling thiamine production via salvage of breakdown products. Similarly, as reported by Gao et al. (2014), *Auxenochlorella* cannot utilize urea as an N source and genes responsible for urea assimilation (*DUR1*, *DUR2*) and transport (*DUR3*) have been lost. Several other transporters present in at least one copy in both *C. vulgaris* and *C. reinhardtii* are also absent in UTEX 250-A, including the borate transporter *BOR1* and the *SLT-*family sodium/sulfate co-transporters. Other transporter gene families feature a single copy in UTEX 250-A, including the *CTR*-type copper ion transporters (three genes in each of *C. vulgaris* and *C. reinhardtii*) and the *PTB*-family sodium/phosphate symporters (four genes in *C. vulgaris*, at least nine in *C. reinhardtii*).

The reduction in gene complexity can also be seen by BUSCO (Benchmarking Universal Single Copy Orthologs) (Manni et al. 2021) analysis against the UTEX 250-A predicted proteins. Of the 1,519 chlorophyte BUSCO genes, 1,488 (98%) were identified as complete, with only nine (0.6%) complete and duplicated. One protein was called as fragmented, although it was manually confirmed to be a correct gene model. Of the 30 missing BUSCOs, eight could be manually recovered by cross-checking against the orthogroup analysis. The remaining 22 genes were also undetected when searching BUSCOs against the UTEX 250-A genome, suggesting they are biologically absent and not misannotated (Dataset S10). Most notably, nine of the 22 missing BUSCOs are part of GreenCut2, a set of almost 600 genes conserved among select photosynthetic eukaryotes and absent from non-photosynthetic species (Karpowicz et al. 2011). A specific analysis of genes involved in photosynthesis and sexual reproduction is presented below.

We also analyzed a small number of gene families that have specifically expanded in UTEX 250-A. Import of glucose and galactose in *Chlorella kessleri* is facilitated by three genes encoding H+/Hexose symporters (*HUP1, HUP2* and *HUP3*) (Stadler et al. 1995), and three paralogous genes that are most similar to *C. kessleri HUP2* were identified in the *A. protothecoides* 0710 genome (Gao et al. 2014). We identified five *HUP* genes on both haplotypes of the UTEX 250-A genome, including a tandem array of three genes on chromosome A12/B12 that is disrupted by contig breaks in the 0710 assembly that resulted in misannotation. For orthogroups featuring genes from UTEX 250-A, *C. vulgaris* and *C. reinhardtii* (Figure 5B), we identified 13 cases involving UTEX 250-A specific expansion via multiple gene duplications (Dataset S11). A tandem array of four genes on A07/B12 encode predicted ribonuclease (RNAse) T2 proteins, which have diverse functions including roles in stress response and pathogen defense (MacIntosh 2011), although the function of the single RNAse T2 gene from *C. reinhardtii* is unknown. A tandem array of six (A01) or five (B01) genes encode homologs of *C. reinhardtii* ferrireductase *FRE1*, which reduces Fe(III) to Fe(II) for Fe assimilation (Allen et al. 2007). Four of the expanded orthogroups are associated with glycosylation. Three of these correspond to gene families (*HPAT*, *RRA* and xyloglucanase-113) that are specifically involved in arabinose *O*-glycosylation of the Hyp-rich glycoproteins (HRGPs) that form the vegetative and zygote cell walls in *C. reinhardtii* (Joo et al. 2017). With respect to proteins potentially involved in cell wall formation, it is also notable that UTEX 250-A carries multiple genes encoding polyketide synthases, as reported previously for *A. protothecoides* 0710 (He et al. 2016; Heimerl et al. 2018). These include homologs of *PKS1* from *C. reinhardtii*, encoding a type I polyketide synthase involved in formation of the zygote cell wall (Heimerl et al. 2018), and *LAP5* from *Arabidopsis thaliana*, encoding a type III polyketide synthase involved in the formation of sporopollenin (Kim et al. 2010; He et al. 2016). Further functional characterization of these expanded gene families may provide insights into the unique biology of *Auxenochlorella* species.

Of the 146 and 655 genes that are uniquely present in the AHP lineage or only in UTEX 250-A, respectively, only 10% encode predicted proteins with an identifiable domain. Most notably, 61 haplotype A genes are present in five AHP or *Auxenochlorella-*specific orthogroups that are associated with pentapeptide repeat domains, and a further 39 genes are present in a single similarly associated orthogroup that otherwise features only one gene in each of the Chlorellales species (including *P. cutis*). Genes encoding pentapeptide repeat proteins are abundant in many bacterial genomes, particularly those of some cyanobacteria, although their functions remain elusive (Vetting et al. 2006). Two highly conserved GreenCut2 proteins feature a pentapeptide repeat domain of unknown function, CPLD59 and CPLD44, and are localized to the thylakoid lumen (Kieselbach et al. 1998; Schubert et al. 2002). Some mycobacterial pentapeptide repeat proteins adopt a fold that mimics double-stranded DNA and can compete with DNA in DNA-protein interactions (Feng et al. 2021). Although their functions are unclear, these 100 genes are dispersed throughout the UTEX 250-A genome, and their repetitive nature can apparently mediate genomic rearrangements via ectopic recombination (Figure 2B, C).

### *Auxenochlorella* encodes a reduced set of proteins related to photosynthesis and sexual reproduction

#### The GreenCut

Intrigued by the potential loss of core photosynthesis genes, we next queried the UTEX 250-A and *C. vulgaris* annotated proteins against the entire GreenCut2 dataset, using *C. reinhardtii* proteins as a reference set. GreenCut2 is divided into four categories: genes conserved among green algae and land plants (CGL, i.e., the green lineage), the green lineage and the red alga *C. merolae* (CPL), the green lineage and diatoms (CGLD), and all of these groups (CPLD). While 12.5% of the GreenCut2 genes were not found in *C. vulgaris*, 18.0% were absent in UTEX 250-A, corresponding to the specific loss of 33 photosynthesis-related genes despite retaining photosynthesis (Figure 5E; Dataset S12). For example, UTEX 250-A encodes a reduced repertoire of light-harvesting complex proteins. Three of the *LHCA* genes that encode the antenna system of photosystem I are absent, namely *LHCA4, LHCA5* and *LHCA6*, but are present in all analyzed trebouxiophytes including *C. vulgaris* (Figure 5E; Dataset S13). This suggests that the PSI-LHCI supercomplex of *Auxenochlorella* likely consists of a single LHCI tetramer (with subunits LHCA1, LHCA8, LHCA7 and LHCA3) and one LHCI dimer (LHCA2 and LHCA9), as in *Dunaliella* species (class Chlorophyceae) (Caspy et al. 2020; Liu et al. 2025). The *PSBS* and *LHCSR* genes, encoding light-harvesting complex-like subunits involved in nonphotochemical quenching (Allorent et al. 2016), are also absent in *Auxenochlorella* UTEX 250-A but present in *C. vulgaris* (Cecchin et al. 2019), raising the question of how *Auxenochlorella* dissipates excess excitation energy.

The absence of certain GreenCut2 genes in *Auxenochlorella* demonstrates that they are not essential for photosynthesis despite their deep conservation in photosynthetic organisms. Nevertheless, we note that with comparison restricted to *C. reinhardtii*, some false negatives are expected. Small, divergent proteins with organellar targeting peptides would be especially susceptible to exclusion in orthology analysis. One example is RBCX (Dataset S12), a protein chaperone that is essential for proper assembly of the RuBisCo large subunit into the octameric RbcL_8_ intermediate, and subsequent association with RuBisCo small subunit to form the holoenzyme (Saschenbrecker et al. 2007; Liu et al. 2010). Eukaryotic RBCX proteins are found in two distinct isoforms; RBCX-I is most similar to cyanobacterial RbcX, and RBCX-II is more divergent (Saschenbrecker et al. 2007; Bracher et al. 2015). While *A. thaliana* has both forms, the *C. reinhardtii RBCX2A* and *RBCX2B* genes both encode RBCX-II proteins (Kolesiński et al. 2011; Bracher et al. 2015). We identified an ortholog of the *C. reinhardtii* RBCX genes in *C. vulgaris,* but not in UTEX 250-A, despite RBCX chaperone function likely being indispensable. Manual searches using the RBCX-I proteins from *A. thaliana* and *Synechocystis* sp. PCC 6803 did reveal a candidate protein (UTEX250_A12.36490, Figure S9A) featuring an RbcX-like domain and a 56 amino acid N-terminus plastid transit peptide predicted by TargetP-2.0 (Almagro Armenteros et al. 2019). Interestingly, *C. vulgaris* also encodes a highly similar protein, suggesting that both RBCX-I and RBCX-II isoforms are present in the species (as in *A. thaliana,* Figure S9B), whereas UTEX 250-A has only RBCX-I and *C. reinhardtii* only RBCX-II. Comparisons of monomer AlphaFold structure models of the putative RBCX-I proteins from UTEX 250-A and *C. vulgaris* with the corresponding crystal structures from *Synechocystis* sp. PCC 6803 (Tanaka et al. 2007) and *A. thaliana* (Kolesiński et al. 2013) confirmed structural homology, supporting the contention that they perform the same RbcL chaperone function (Figure S9C).

### The CiliaCut and meiosis genes

The putative hybrid origin of UTEX 250 suggests the presence of a cryptic life cycle stage in *Auxenochlorella* species, which may include meiosis of otherwise vegetatively diploid cells and the formation of gametes (or at least implies an asexual transition between diploid and haploid cells). Direct evidence for sexual reproduction in the Trebouxiophyceae is scarce, although Fučiková et al. (2015) identified a core set of nine meiosis genes that are mostly present in trebouxiophyte genomes. Although *Chlorella* species have never been observed to form ciliated gametes, Blanc et al. (2010) identified orthologs of *C. reinhardtii* “CiliaCut” genes, which are conserved among ciliated organisms but absent from species without cilia (Merchant et al. 2007). Specifically, *Chlorella variabilis* has genes encoding the outer and inner dynein arm proteins, but lacks most or all of the genes encoding components of the intraflagellar transport particle, radial spoke and central pair complex (Blanc et al. 2010). The cilia genes encoded by *C. variabilis* significantly overlap with those present in the centric diatom *Thalassiosira pseudonana* (i.e., the “CentricCut”), which produces male gametes with motile cilia (Moore et al. 2017). Cecchin et al. (2019) confirmed this result, finding an even more substantial intersect between “MotileCut” cilia genes common to *C. vulgaris* and *T. pseudonana*, but absent in *Caenorhabditis elegans* (which produces sensory cilia, but not motile cilia), suggesting that *Chlorella* species may carry the necessary repertoire of genes to produce motile ciliated gametes. Most recently, Gazquez et al. (2024) demonstrated that the ciliated life stage of the lichen-forming *Trebouxia lynniae* corresponds to gametes, providing one of the few characterizations of a sexual lifecycle in a trebouxiophyte alga.

Only 25.3% of CiliaCut genes from *C. reinhardtii* are present in the UTEX 250-A genome, relative to 38.5% in *C. vulgaris* (Figure 5F). However, 58.3% of the genes that form the intersect of the MotileCut and CentricCut are present in UTEX 250-A, relative to 66.7% in *C. vulgaris*, corresponding to only four genes that are uniquely absent in UTEX 250-A (Dataset S14).

Fučiková et al. (2015) identified four of the nine core meiosis genes in the *A. protothecoides* 0710 genome, with *HOPP1*, *REC8*, *MER3*, *MSH4* and *MSH5* absent. We identified *HOPP1* and *REC8* in UTEX 250-A (Figure 5F), suggesting that they were previously missed due to misannotation. *MSH4* and *MSH5*, homologs of the bacterial mismatch repair *MutS* family, are absent in some sexual species such as *Plasmodium vivax*, although *MER3*, encoding a helicase involved in the formation of meiotic crossovers (Altmannova et al. 2023), is universally present in sexual species (Fučiková et al. 2015). We also identified UTEX 250-A orthologs of two critical genes for syngamy, *GEX1*, which functions in nuclear fusion (Ning et al. 2013), and *HAP2*, which functions in gamete fusion (Fedry et al. 2017). Thus, *Auxenochlorella* species may be able to reproduce sexually, possibly via the formation of ciliated gametes, although as with photosynthesis the set of genes functioning in these processes has been substantially reduced.

Notably, our analyses confirm the prior observation that all trebouxiophytes have lost the KNOX/BELL homeodomain transcription factors (Joo et al. 2018), typified by the *C. reinhardtii* genes *GSP1* and *GSM1*, that regulate haploid-to-diploid transitions in Archaeplastida (Lee et al. 2008; Hisanaga et al. 2021; Hirooka et al. 2022). Thus, the molecular pathways underlying putative sexual lifecycles in both haplontic and diplontic trebouxiophytes remain cryptic.

### Periodic cytosine and adenine methylation supports widespread internal antisense transcription

#### DNA methylation

The DNA modification 5-methylcytosine (5mC) is widespread across all domains of life and has several essential functions (Mattei et al. 2022), although relatively little is understood about cytosine methylation in green algae. In *C. variabilis*, gene bodies are extensively methylated at CpG dinucleotides, whereas promoter regions are hypomethylated (Zemach et al. 2010).

Although 5mC is generally restricted to centromeres and subtelomeres in *C. reinhardtii* (Lopez et al. 2015; Chaux-Jukic et al. 2021; Craig et al. 2023), extensive gene body methylation was recently shown for the chlorophyceaen alga *T. obliquus* (Biondi et al. 2024), suggesting that this pattern may be widespread in chlorophytes. In the extremely compact genomes of prasinophytes such as *Ostreococcus lucimarinus* and *Micromonas pusilla*, CpG dinucleotides in gene bodies are also highly methylated, although 5mC occurs in periodic clusters that correspond to the linker DNA between nucleosomes (Huff and Zilberman 2014). Notably, periodic adenine methylation (6mA), mostly at ApT dinucleotides, was also discovered in *C. reinhardtii* (Fu et al. 2015). Periodicity of 6mA is also due to specific methylation of nucleosome linkers, although unlike for 5mC, the adenine methylation is present around TSSs and not in the gene body.

Romero Charria et al. (2024) demonstrated that periodic adenine methylation of linker DNA downstream of TSSs is also present in both prasinophytes and *C. variabilis*, as well as in other distantly related taxa, suggesting that it is an ancient feature of eukaryotic genomes. The function of 6mA in this context is not known; the biophysical properties of this modification may play some role in promoting transcription (Bochtler and Fernandes 2021), and 6mA could also facilitate the coordination of promoter chromatin marks such as H3K4me3 (Romero Charria et al. 2024).

We used the ONT reads from CCAP 211/7C (equivalent to UTEX 250, Figure 3A) to call both 5mC at CpG sites and 6mA at ApT sites across the UTEX 250-A genome. Using only the high-confidence TSSs from IsoSeq-based gene models, periodicity of both modifications was evident (Figure 6A). Cytosine methylation is essentially absent at sites adjacent to the majority of TSSs, and universal at the hypermethylated peaks within gene bodies (on average 5mC is called at ∼95% of CpG sites at the highest point of the peaks). Conversely, two periodic peaks of 6mA are present downstream of the TSS, although far fewer ApT sites are methylated and the signal is noisier between the peaks (Figure S10). Assuming that the peaks correspond to nucleosome linkers, it appears that the first two linkers downstream of the TSS are specifically associated with adenine methylation, whereas cytosine methylation is present at the third linker and reaches a stable maximum at subsequent linkers. These patterns closely resemble those of the prasinophytes, as opposed to *C. variabilis* where 6mA exhibits periodicity but 5mC is relatively constant across the gene body (Romero Charria et al. 2024). Huff and Zilberman (2014) proposed that 5mC periodicity contributes to nucleosome positioning and hypothesized that this could be an adaptation to extremely compact nuclei and small cell sizes. Our results suggest this unusual genomic architecture may have been present in the common ancestor of the core-Chlorophyta (Trebouxiophyceae and Chlorophyceae in Figure 1A), or alternatively that it re-evolved on the lineage leading to *Auxenochlorella*.

**Figure 6.**
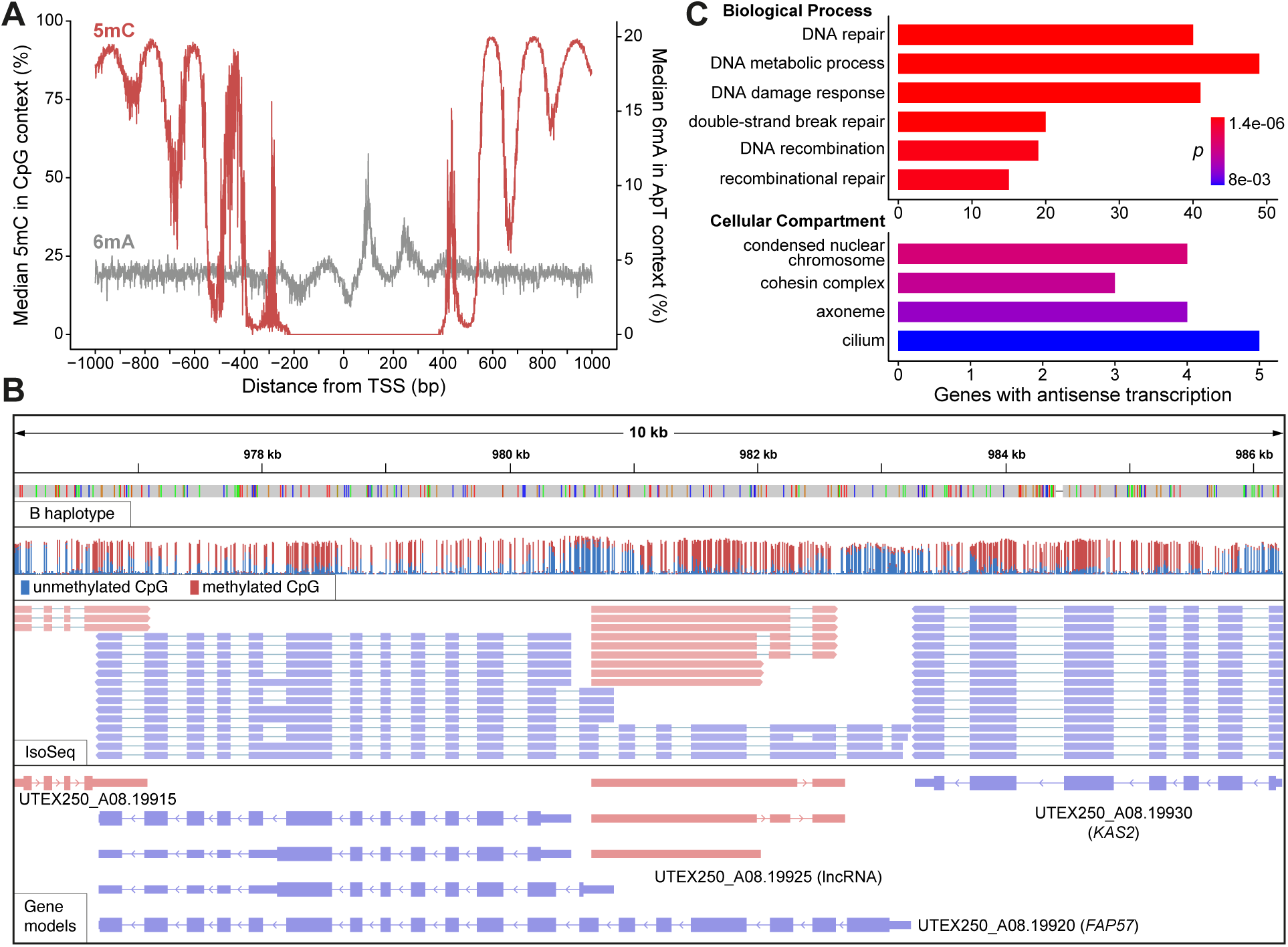
Promoter methylation and antisense lncRNAs in *Auxenochlorella* UTEX 250-A. A) Median per site 5mC (red) and 6mA (grey) around transcription start sites. B) IGV screenshot showing example of internal bidirectional promoter producing antisense lncRNA and truncated isoform for *FAP57* (UTEX_A08.19920). The top row shows the B haplotype of the UTEX 250-A genome mapped against the A haplotype, with colored bands corresponding to single nucleotide polymorphisms. The second row shows read coverage of ONT reads from CCAP 211/7C where only CpG sites are shown. The proportion of red to blue corresponds to the proportion of 5mC calls in the ONT reads. The third row shows IsoSeq data, where pink reads are on the forward strand, and blue reads the reverse. The final row shows the annotated gene models, where the thin lines correspond to introns, intermediate lines to UTRs, and thick lines to coding sequence. C) GO term enrichment analysis for 780 genes overlapped by antisense lncRNAs. See Dataset S16 for full results.

#### Antisense long non-coding RNAs

Aside from the biological importance of periodic methylation, these signatures also provide evidence for promoter regions, similar to H3K4me3 ChIP-seq datasets that only exist for a limited number of green algae (Strenkert et al. 2022; Petroll et al. 2025). Figure 6B shows a representative locus featuring four genes where CpG hypomethylation clearly corresponds to the regions downstream of the IsoSeq-based TSSs. Notably, one of these genes (UTEX250_A08.19925) is a lncRNA transcribed from a bidirectional hypomethylated promoter located within the CiliaCut gene *FAP57* (UTEX250_A08.19920), which is essential for the correct assembly of inner dynein arms (Lin et al. 2019). While manually curating genes in the UTEX 250-A genome, we noticed many similar cases, which motivated a thorough search for lncRNA genes that are supported by multiple IsoSeq reads (Table 1). After manual verification, 165 protein-coding genes were identified with internal bidirectional promoters that produced antisense lncRNAs on at least one of the two haplotypes (Dataset S15). For genes such as *FAP57*, it is unclear whether the truncated isoforms transcribed from the internal promoter could be functional, while for many others the truncated isoforms clearly have no coding capacity. We also identified 615 protein-coding genes that exhibit substantial overlap with an antisense lncRNA (>30% of the transcript length) transcribed from an internal, or immediately adjacent, unidirectional promoter. The annotated lncRNAs are expected to be polyadenylated based on the IsoSeq protocol and ∼59% are spliced.

To explore the potential functions of widespread antisense transcription in UTEX 250-A, we analyzed the predicted functions of the genes with associated lncRNAs. Manual curation of proteins encoded by genes with internal bidirectional promoters recovered potential functions associated with DNA, such as repair, chromatin and replication, followed by functions related to post-translational regulation, cilia, and the cell wall. A gene ontology (GO) enrichment analysis focusing on the full set of 780 genes with evidence for antisense transcription supported these observations. The most significant terms for biological processes were all related to DNA repair and recombination, including the repair of double-strand breaks by homologous recombination (Figure 6C, Dataset S16). Indeed, the list includes many fundamental DNA repair genes, including *RAD51*, *RAD21*, the RECQ helicases *RECQ1* and *RECQ5*, and the DNA polymerases *POLK1* and *POLQ2* (Dataset S15). For cellular compartments, the two most significant terms, “condensed nuclear chromosome” and “cohesin complex”, are both related to mitosis, meiosis and recombination, whereas the next two significant terms are related to cilia. The most significant molecular function term, “calcium ion binding” (Dataset S16), is also associated with cilia genes. Indeed 14 of the CiliaCut genes (including *FAP57*) exhibit antisense transcription, alongside five of the eight UTEX 250-A core meiotic and syngamy genes (Figure 5F), representing a significant enrichment (χ^2^= 32.05, *p* = 1.5×10^−8^). The nuclear fusion gene *GEX1* represents an extreme case where all of the IsoSeq reads at the locus are antisense (Figure S11).

Based on these results, we speculate that antisense lncRNAs may play a regulatory role in *Auxenochlorella*. In synchronized cultures of *C. reinhardtii*, some DNA repair genes are specifically upregulated at the transition from dark to light (e.g., the ribonucleotide reductase subunit *RIR2L*, which features an antisense lncRNA in UTEX 250-A), suggesting a possible role in photodamage response (Strenkert et al. 2019). Many human DNA repair genes are also upregulated under genotoxic stresses such as UV radiation and ROS (Christmann and Kaina 2013). In *C. reinhardtii*, several of the core meiotic genes are among a small set of genes that lack H3K4me3 at their TSSs during vegetative growth, supporting expression restricted to the sexual cycle (Strenkert et al. 2022). It therefore seems plausible that genes overlapped by antisense lncRNAs must be specifically translated at certain points of the cell and life cycles, or under specific conditions. While this gene set is enriched for putative functions in DNA repair and sex, we also observe this phenomenon for many genes that are predicted to have unrelated functions, suggesting that this may be a general regulatory mechanism. Antisense lncRNAs can regulate expression of associated protein-coding genes by several mechanisms (Werner et al. 2024). For example, in compact genomes such as *S. cerevisiae*, transcriptional interference can occur via the physical collision of polymerase complexes on opposite strands (Prescott and Proudfoot 2002; Hobson et al. 2012). Experimental characterization, including differential expression analyses of the protein-coding and lncRNA gene pairs at these loci, will be required to test this hypothesis. However, we are not aware of similar observations in other green algae, suggesting widespread antisense transcription may have emerged as an unusual gene regulatory mechanism during *Auxenochlorella* evolution. One notable observation is that the UTEX 250-A genome encodes only 15 predicted F-box domain proteins, relative to 43 in *C. vulgaris* and 38 in *C. reinhardtii* (based on the Pfam domains PF12937 and PF00646). F-box proteins function in post-translational regulation as a component of the SCF (Skp, Cullin, F-box containing) ubiquitin-ligase complex, with the F-box protein responsible for binding specific substrates destined for proteolysis (Kipreos and Pagano 2000). Although currently speculative, it is possible that transcriptional regulation of at least a subset of fundamental genes by lncRNAs emerged in the compact *Auxenochlorella* genome in contrast to a decrease in regulation via protein degradation.

### An *Auxenochlorella* genetic toolkit: reverse genetics, selectable markers, inducible promoters, and fluorescent reporters for cellular localization

In combination with the genomic resources presented thus far, nuclear gene targeting by homologous recombination in *Auxenochlorella* facilitates reverse genetic analysis of biochemical pathways and processes. To illustrate this capability, we targeted *CHL27*, encoding the chlorophyll (Chl) biosynthesis enzyme Mg-protoporphyrin IX monomethylester (MgPMME) aerobic cyclase, reasoning that loss of Chl would provide a clear visual phenotype, and that non-photosynthetic mutants could be maintained heterotrophically on glucose in the dark. The UTEX 250-A *CHL27* locus is heterozygous and exemplifies the hybrid origin of the strain. The *CHL27-1* allele on chromosome B12 is closely related to the corresponding heterozygous *CHL27* alleles in UTEX 25, and the homozygous *CHL27* locus in UTEX 2341; conversely, *CHL27-2* on A07 has more polymorphisms in common with the heterozygous CCAP 211/61 *CHL27* alleles (Figure 7A, S12A). *CHL27* polymorphisms cluster in the introns and in the flanking intergenic region between the *CHL27* 3’ UTR and the downstream gene, and all but one of the coding sequence polymorphisms are synonymous, so six of the seven alleles encode identical polypeptides (Figure S12A, B). A non-synonymous polymorphism in UTEX 2341 *CHL27* substitutes Arg for Gly at position 33, but this change is in the region predicted to encode the plastid transit peptide, so all of the alleles are expected to produce identical mature polypeptides after chloroplast import and transit peptide cleavage (Figure S12B).

**Figure 7.**
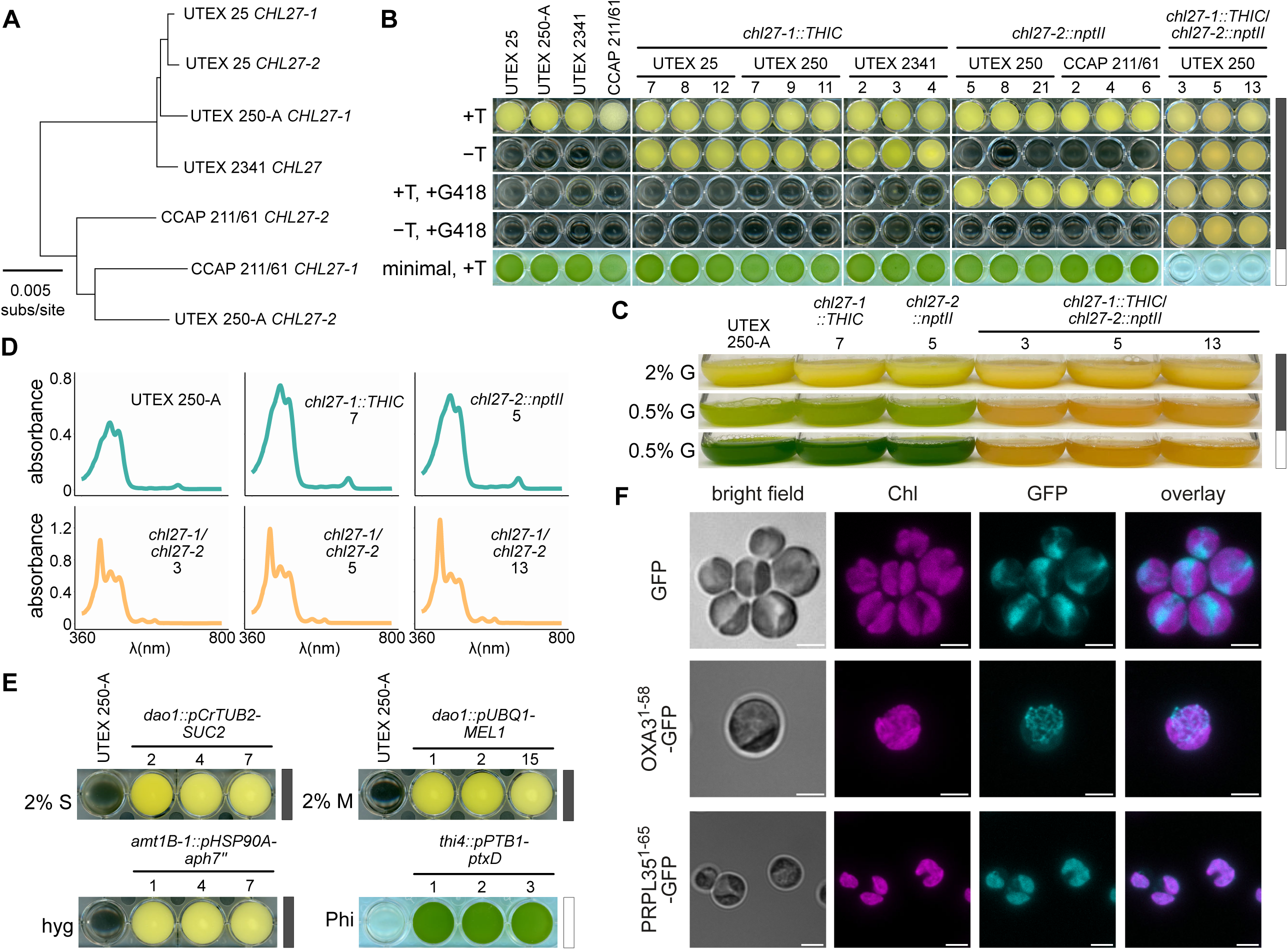
*Auxenochlorella* reverse genetics and genetic toolkit. A) Neighbor-joining phylogeny of *CHL27* alleles in UTEX 25, 250-A, 2341 and CCAP 211/61. Sequence comparisons encompassed the *CHL27* alleles and the 5’ and 3’ flanking regions used to make targeting constructs (see Figure S12A). All bootstrap values >70%. B) Selection and photosynthetic growth phenotypes of representative thiamine auxotrophic and G418-resistant transformants versus the corresponding wild-type controls. Cultures were grown for five days at 26°C with 140 rpm shaking in the dark, or for ten days at 24°C with approximately 40 μmol.m^− 2^.s^−1^ of white light provided by cool white fluorescent bulbs. All strains grew in medium supplemented with 2 μM thiamine (+T), but only transformants expressing *THIC* grew in medium without added thiamine (- T). Resistance to 100 μg/mL of G418 (+T, +G418) was conferred by *nptII*. *chl27* double knockout strains grew under both selection conditions (-T, +G418). Wild-type and *chl27* heterozygotes grew photoautotrophically, but *chl27* homozygous mutants were non-photosynthetic (minimal, +T). C) Pigment accumulation under different trophic conditions in 25 mL flask cultures of UTEX 250-derived *chl27* heterozygous and homozygous mutants versus UTEX 250-A. Cultures were grown with 2 μM thiamine and 20 g/L (2% G) or 5 g/L (5% G) of glucose for three days at 28°C, with 200 rpm shaking, in the dark or with approximately 5 μmol.m^−2^.s^−1^ of white light provided by warm white LEDs. D) Absorption spectra of extracts from UTEX 250-A, *chl27* heterozygous and homozygous mutant strains. Cultures were grown for four days in the dark with 20 g/L glucose at 24°C, 160 rpm in 24-well plates. Shoulders at 421 nm and peaks around 450 nm and 475 nm are consistent with absorption by lutein and zeaxanthin, and a minor peak at 663 nm reflects a small amount of chlorophyll in heterotrophic wild-type and *chl27* heterozygotes. Peaks at 418, 551 and 590 nm in extracts from chl27 double knockouts are indicative of absorption by Mg-protoporphyrin IX or Mg-PMME. E) Selectable marker growth phenotypes of transformants compared to UTEX 250-A. Strains were cultured for four days in the dark at 26°C with 140 rpm shaking, in 20 g/L sucrose (2% S), 20 g/L melibiose (2% M), 300 μg/mL hygromycin B (hyg), or for 5 days with approximately 40 μmol.m^−2^.s^−1^ of white light provided by cool white fluorescent bulbs, at 24°C with 140 rpm shaking, in tris-acetate medium containing 1.1 mM phosphite (Phi). *SUC2* was targeted to the *DAO1* locus, and driven by the *TUB2* promoter from *C. reinhardtii* (*pCrTUB2*); *MEL1* was integrated at *DAO1* and driven by the endogenous *UBQ1* promoter (*pUBQ1*); *aph7’’* was controlled by the promoter of *HSP90A* (*pHSP90A*) and targeted to *AMT1B-1*; and *ptxD*, under the control of the *PTB1* promoter (*pPTB1*), was integrated at *THI4*. F) Fluorescence imaging of UTEX 250-A transformants expressing sucrose invertase and GFP. Cells were from cultures grown with 5 g/L of sucrose for three to four days at 25°C, shaking at 160 rpm, and approximately 40 μmol.m^−2^.s^−1^ of white light provided by cool white fluorescent bulbs.

We designed UTEX 250 *CHL27-1* and *CHL27-2* targeting constructs to fully eliminate the coding sequence of each allele (Figure S12C, D). Since thiamine auxotrophy is a characteristic of all *Auxenochlorella* and *Prototheca* species (Pore 1972), we used *THIC* from *A. thaliana* (Franklin et al. 2013; Moseley et al. 2024) as the marker for transformation targeting *CHL27-1*. Neomycin phosphotransferase II (*nptII*), conferring resistance to G418 (Geneticin) (Franklin et al. 2011; Moseley et al. 2024; Dueñas et al. 2025a), was the selectable marker for targeting *CHL27-2*. We did not make allele-specific targeting constructs for UTEX 25, UTEX 2341 or CCAP 211/61, reasoning that the UTEX 250 *CHL27* homology arms should have sufficient sequence identity to enable recombination (Figure S12A). Representative *chl27-1::THIC* transformants from UTEX 25, UTEX 250 and UTEX 2341 are thiamine prototrophs, while *chl27-2::nptII* integration into UTEX 250 and CCAP 211/61 conferred resistance to G418 (Figure 7B). Integration of the constructs at the *CHL27* loci was confirmed by PCR amplification of genomic DNA using primers flanking the homology arms (Figure S12E), and subsequent sequencing of the PCR products. *CHL27-1* was disrupted in 8/11 thiamine prototrophic UTEX 250 transformants and 3/11 had the integrating construct mis-targeted to the *CHL27-2* allele, while integration at the *CHL27-2* locus was confirmed in 9/9 G418-resistant UTEX 250 transformants. Genetic tractability appears to be a feature of the *Auxenochlorella* genus, as evidenced by successful transformation and targeted gene replacement by homologous recombination in both *A. protothecoides* and *A. symbiontica* strains, and allele-specific targeting in the UTEX 250 hybrid. Integration into the nuclear genome occurs via homologous recombination in almost all *Auxenochlorella* transformants, in contrast to 2% of transformants reported for *Ostreococcus* (Lozano et al, 2014). We have documented a step by step guide to transformation of *Auxenochlorella* using a lithium acetate/polyethylene glycol method on protocols.io (Dueñas et al. 2025b).

Representative *chl27* loss-of-function mutants (*chl27-1::THIC*/*chl27-2::nptII*) were generated by sequential targeting of the *CHL27* alleles. Double knockout strains were both thiamine prototrophs and G418-resistant (Figure 7B). Heterotrophic growth was robust for all strains, but *chl27* mutants bleached and were incapable of photoautotrophic growth on minimal media (Figure 7B). Photoautotrophic growth of single allele knockouts was equivalent to the wild-type parents, demonstrating that one copy of *CHL27* was sufficient to maintain adequate Chl for photosynthesis under the growth conditions (Figure 7B). In agreement with the observations of Shihira-Ishikawa and Hase (1964), wild-type UTEX 250-A and the *chl27* heterozygous mutants grown under heterotrophic conditions with a high C/N ratio (2% glucose) contained minimal amounts of Chl, with the yellow color attributed to xanthophyll pigments (Figure 7C, D). Partial greening was observed in wild-type and single-allele knockout cultures grown with limiting glucose (0.5%) in the dark, and full greening was stimulated by just 5 μmol.m^−2^.s^−1^ of white light (Figure 7C). In contrast, representative *chl27* double knockout cultures accumulated a red pigment under the same growth conditions (Figure 7C). Absorbance spectra from cell extracts of wild-type and single allele knockout strains grown in the dark with 2% glucose had minor Chl peaks at 663 nm, and were consistent with lutein and zeaxanthin as the major xanthophyll pigments (Figure 7D). Chl was not detected in the *chl27* double knockouts, which instead had a major peak at 418 nm with minor peaks at 551 and 590 nm (Figure 7D), indicative of Mg-protoporphyrin IX or MgPMME accumulation (Tottey et al. 2003; Hollingshead et al. 2012). This is the expected result of blocking the aerobic cyclase that converts red MgPMME to green divinyl protochlorophyllide (Chen et al. 2021). The photosensitive phenotype of *chl27* mutants (Figure 7B) is likely a consequence of reactive oxygen species damage caused by the highly photoactive protoporphyrin pigments (Mock 2001).

To expand our capabilities for gene targeting and metabolic engineering, we developed additional selectable markers for transformation (Figure 7E). Two additional selectable markers have been described in *Prototheca moriformis*: *SUC2*, encoding secreted sucrose invertase from *S. cerevisiae*, enables heterotrophic growth on sucrose (Franklin et al. 2011), whereas *MEL1*, encoding secreted α-galactosidase from *Saccharomyces carlsbergensis*, enables heterotrophic growth on melibiose (Franklin et al. 2013). We demonstrated that *Auxenochlorella* transformants expressing *SUC2* or *MEL1* consume sucrose or melibiose, respectively, for heterotrophic growth (Figure 7E, S13A). The secreted SUC2 and MEL1 glycosyl hydrolases act in a non-cell-autonomous manner, making rigorous single-colony purification essential to avoid persistence of untransformed parental cells. Another aminoglycoside 3’-phosphotransferase antibiotic resistance gene, *aph7”*, from *Streptomyces hygroscopicus*, is commonly used as a *C. reinhardtii* transformation marker (Berthold et al. 2002), and confers resistance to 300 μg/mL hygromycin B to *Auxenochlorella* (Figure 7E, S13A). As was reported for *C. reinhardtii* (Loera-Quezada et al. 2016), phosphite oxidoreductase, encoded by *ptxD* from *Pseudomonas stutzeri* (Costas et al. 2001), enables mixotrophically grown *Auxenochlorella* to utilize phosphite as a P-source (Figure 7E, S13A). Together with *nptII* and *THIC*, described above, the four selectable markers illustrated in Figure 7E can be used in serial transformations to generate homozygous knockout mutants at up to three loci, or to make heterozygous mutations at six independent loci.

We next set out to establish fluorescent protein expression in *Auxenochlorella* for intracellular localization studies. *GFP* and *GFP* fusions were driven by the RuBisCo small subunit (*RBCS1*) promoter and targeted to the neutral *DAO1* locus (Dueñas et al. 2025a). Strains were cultured in the light under mixotrophic conditions such that transgene expression was activated in green cells. Chl and GFP fluorescence imaging by confocal microscopy showed that untargeted GFP was located in the cytoplasm (Figure 7F, S13B). Fusion of GFP to the mitochondrial targeting sequence of the OXA3 membrane insertase revealed a mitochondrial network encircling the cytoplasm (Figure 7F, S13B), while the chloroplast ribosomal protein L35 (PRPL35) plastid transit peptide targeted GFP to the chloroplast, as evidenced by co-localization of GFP fluorescence and Chl autofluorescence (Figure 7F, S13B). These strains may serve as references for the localization of cytoplasmic, mitochondrial and plastid-targeted proteins in future research.

Next, we asked whether one of the inferred trisomies in UTEX 250-A (Figure 4A) could be confirmed by allele-specific targeting. To do so, we took advantage of the characteristic metabolic switch of *Auxenochlorella*. The ammonium transporter gene *AMT1B*, which is highly upregulated under N-depletion in heterotrophic growth in UTEX 25 (Yan et al. 2013), is located on the right arms of chromosomes A03 and B03, with a putative third copy on the duplicate of B03 (termed “C03”, Figure 8A). Since A03 and B03/C03 are heterozygous at this region, we designed an allele-specific construct with homology arms that specifically corresponded to the flanking sequences of the *AMT1B* allele of B03/C03 (termed *AMT1B-1*, Figure 8A). The construct replaced the *AMT1B-1* ORF with an ORF encoding the Venus fluorescent protein, which is under the control of the endogenous *AMT1B-1* promoter. As above, *nptII* served as a transformation marker conferring resistance to G418. We designed allele-specific PCR primers that targeted the left and right flanks of the endogenous *AMT1B* alleles, in addition to primer pairs that targeted the construct, with one primer within the construct and the other in the sequence flanking the homology arms (Figure 8A). Wild-type sequences for both *AMT1B* alleles (*AMT1B-1* on B03 or C03, *AMT1B-2* on A03) could be amplified from wild-type UTEX 250-A and from three representative transformants, but the transformants also contained mutant *amt1B-1::Venus* integrations, consistent with a trisomic state featuring one copy of *AMT1B-2* and two copies of *AMT1B-1* in the wild-type nuclear genome (Figure 8B). We next compared Venus fluorescence between transformants and the UTEX 250-A wild-type negative control grown under photoautotrophic and heterotrophic conditions (Figure 8C, D). Since biomass concentration is higher in late log-phase heterotrophic cell cultures than in photoautotrophic cultures, the greater depletion of N from the medium leads to the activation of the N-deficiency-responsive *AMT1B* promoter. As a consequence, transformants displayed 19-fold higher Venus fluorescence in heterotrophy than in phototrophy, and on average 66-fold higher Venus fluorescence compared to the wild-type negative control (Figure 8C, D). We previously utilized the promoter of Photosystem I subunit D (*PSAD1*) to induce expression of a synthetic construct in phototrophy (Dueñas et al. 2025a). In addition to confirming trisomy, we here show that the *AMT1B* promoter represents a convenient counterpart for inducing expression in heterotrophy.

**Figure 8.**
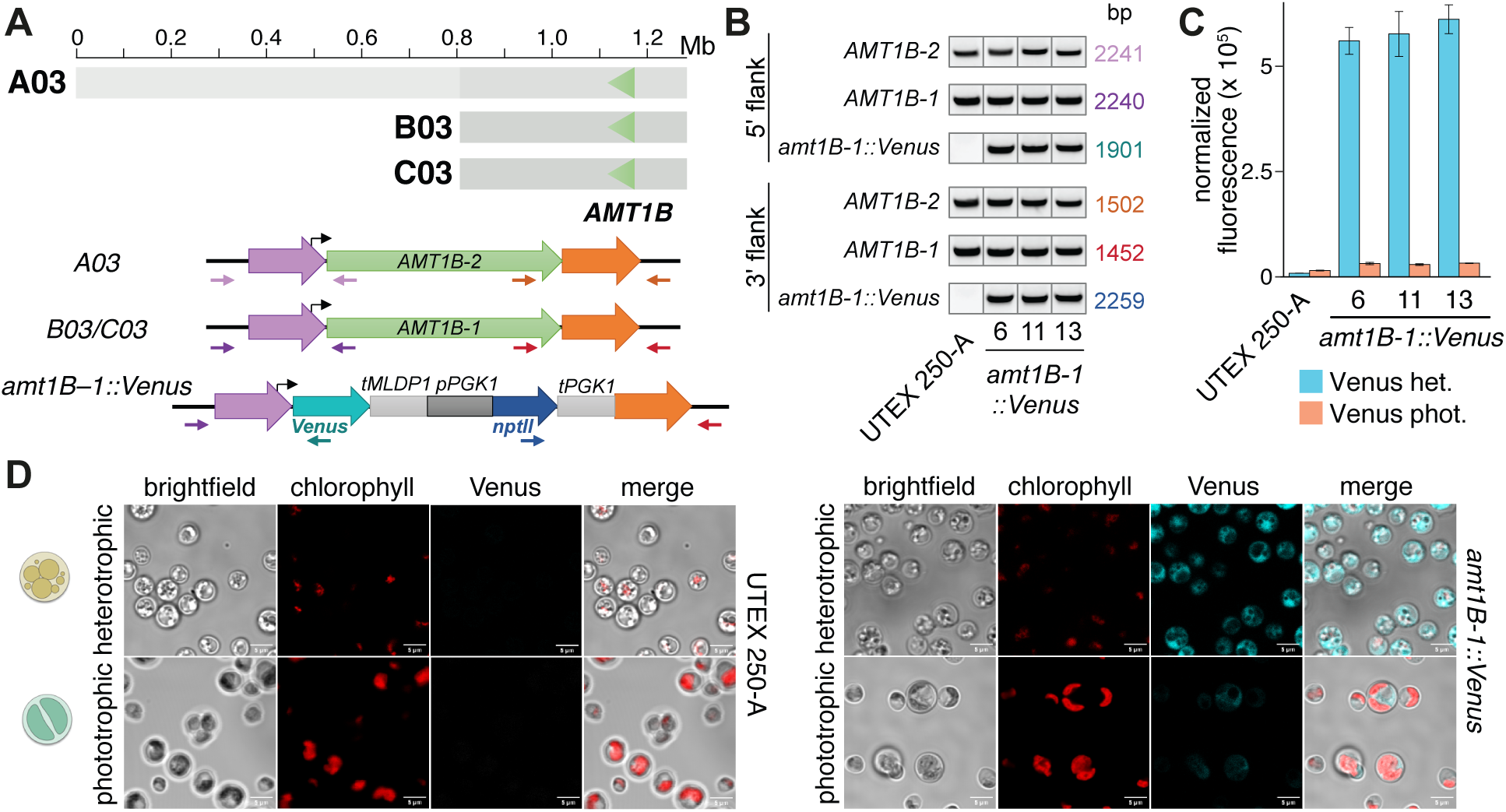
Allele-specific targeting of a trisomic chromosome. A) Schematic of chromosome 3 copies (A03 and B03/C03) with green triangles marking the location of *AMT1B*. The mutant locus features a gene cassette comprised of a *Venus* ORF and a 537 bp terminator sequence containing the endogenous 3’ UTR of *MLDP1* (*MAJOR LIPID DROPLET PROTEIN 1*), and a gene cassette comprised of a *nptII* ORF flanked by the endogenous promoter/5’ UTR and terminator/3’ UTR of *PGK1B* (*PHOSPHOGLYCERATE KINASE 1B*), which is targeted to the *AMT1B-1* allele of B03/C03. *Venus* expression is under the control of the endogenous *AMT1B-1* promoter. B) PCR amplification of *AMT1B-1* (B03/C03) and *AMT1B-2* (A03) 5’ and 3’ flanking regions using allele-specific primers in wild-type UTEX 250-A and three representative *amt1B-1::Venus* transformants, with additional amplification of *amt1B-1::Venus* integration in the transformants. C) Venus fluorescence detection in late-log heterotrophic transformant cultures compared to wildtype UTEX 250-A negative control. D) Comparison of chlorophyll autofluorescence (red) and Venus fluorescence (cyan) in wild-type UTEX 250-A versus a representative transformant under heterotrophic or photoautotrophic conditions using confocal fluorescence microscopy.

Finally, given the effectiveness of homologous recombination in *Auxenochlorella* species, we assessed the UTEX 250-A genome for genes involved in non-homologous end joining. All of the core non-homologous end joining genes (Chang et al. 2017) are present in UTEX 250-A and *C. reinhardtii*, namely Ku70 and Ku80, and DNA ligase IV (*LIG4*) and *XRCC4* (Dataset S17). A gene encoding the DNA dependent protein kinase catalytic subunit (DNA-PKcs) is present in *C. reinhardtii* but is absent in UTEX 250-A, as it also is absent in *A. thaliana* (Khan & Ochi 2023). DNA-PKcs appears to be absent in all Chlorellales genomes analyzed, and its absence is therefore unlikely to explain the effectiveness of homologous recombination in *Auxenochlorella*. Overall, we conclude that *Auxenochlorella* has the core complement of genes functioning in non-homologous end joining.

## DISCUSSION

Species that reproduce clonally as diploids may experience specific phenomena including allodiploid hybridization, mitotic recombination, loss-of-heterozygosity and aneuploidy. Although these processes can act as strong evolutionary forces, for example in speciation or adaptation to new environments, most of our knowledge is presently based upon yeasts and filamentous fungi. In green algae, observations consistent with these phenomena have been reported sporadically, including trisomy in *C. primus* (Lemieux et al. 2019), and chromosome-scale LOH and possible hybridization in *T. obliquus* (Biondi et al. 2024). Here, we find that each of these phenomena occurs in the trebouxiophyte genus *Auxenochlorella*, demonstrating the generality of these important “yeast-like” molecular and evolutionary processes in vegetatively diploid eukaryotes.

*Auxenochlorella* UTEX 250 is an allodiploid hybrid of two closely related species, *A. protothecoides* and *A. symbiontica*, that are differentiated by extensive karyotypic variation and possibly ecological niches. Following hybridization, the parental haplotypes have presumably been shuffled by mitotic crossovers, and homogenized via LOH events mediated by recombination, DNA repair, and potentially transient aneuploidy. Post hybridization, the outcomes of these events can be driven by selection to resolve genomic incompatibilities between the divergent parental genomes in a process termed genome stabilization (Sipiczki 2008; Gabaldón 2020). It is presently unclear to what extent the crossovers, LOH events and aneuploidies of UTEX 250 are involved in stabilization, although at least some of the LOH events that occurred prior to laboratory culture were presumably beneficial (e.g., at rDNA arrays). However, our observations from authentic strains of *A. protothecoides* and *A. symbiontica* suggest that LOH and aneuploidy are not just associated with hybridization: they are generally expected to be prominent evolutionary forces in *Auxenochlorella* that can occur on the timescale of laboratory culture. It is also unclear whether the sampling of UTEX 250 was a rare chance event, or whether it may represent a successful hybrid lineage that is persistent and potentially abundant (albeit likely sterile, assuming a sexual life cycle). Given the number of *Auxenochlorella* strains in culture, and the potential to isolate new strains (Asker and Awad 2019; Chen et al. 2024), it would be interesting to apply population genomics approaches to investigate the prevalence of LOH, aneuploidy, and possibly other hybridization events, in the genus.

The resources and genetic toolkit presented here will serve as a foundation for functional genomics and systems biology analyses, as well as the design of transformation constructs for molecular manipulation of the genome. *Auxenochlorella* has great potential for development as an algal model for fundamental discovery research and bioengineering, and we hope that it will complement existing reference systems. As introduced, *C. reinhardtii* is the premier green alga for forward and reverse genetics, with well-developed classical haploid genetics (Harris 2001), established methods for transforming the nuclear, plastid and mitochondrial genomes (Boynton et al. 1988; Kindle et al. 1989; Kindle 1990; Randolph-Anderson et al. 1993; Shimogawara et al. 1998), well-annotated genome assemblies (Merchant et al. 2007; Craig et al. 2023), a plethora of genetic markers and molecular components (promoters, UTRs, etc.) (Crozet et al. 2018), ribonucleoprotein (RNP)-mediated gene editing (Nievergelt 2025), and extensive community resources including wild-type and mutant collections (Li et al. 2016; Li et al. 2019). Nonetheless, two major hurdles have inhibited routine transformation of the *C. reinhardtii* nuclear genome: the essentially random integration of transgenes by non-homologous end joining, and the transcriptional silencing of transgenes (Schroda 2019; Nievergelt 2025). Recently, RNP-mediated gene-editing has been combined with scar-less homology-directed repair for transgene insertion at specific loci (Ferenczi et al. 2017; Akella et al. 2021; Nievergelt et al. 2023), substantially reducing position effects on transgene expression compared to un-targeted integration, and improving the proportion of transformants with seamless insertions at the recombination site up to 27% (Jacobebbinghaus et al. 2025). Mutant strains that robustly express transgenes have also been characterized (Neupert et al. 2020; Schroda and Remacle 2022).

Despite these advances, targeting of the *Auxenochlorella* nuclear genome by homologous recombination provides several advantages. Transformation only requires linear double-stranded DNA, in contrast to endonuclease-based methods that also depend on guide RNAs and DNA repair templates. Homologous recombination is not constrained by the requirement for double-strand breaks adjacent to Protospacer Adjacent Motif (PAM) sequences and thus provides greater flexibility in integration sites. Efficient integration by homologous recombination appears to be a general property of wild-type *Auxenochlorella* strains, and extends to species from the related genus *Prototheca* (Franklin et al. 2013; Moseley et al. 2024). Transgene integration occurs at high frequency via homologous recombination in the red algae *C. merolae* and *Galdieria partita* (Minoda et al. 2004; Fujiwara et al. 2013; Hirooka et al. 2022), but among green algae appears to be unique to *Auxenochlorella* and *Prototheca*. We have observed that the majority of *Auxenochlorella* transformants, typically 70-100%, exhibit precise, allele-specific integration at the targeted locus (Figure 7, 8). The upper limit for deletions is not yet defined, but in this work homology arms that are separated by more than 2 kb were used to delete the entire CDS of *CHL27* and *AMT1B* (Figure 7, 8, S12). Moreover, large cassettes containing up to four transgene expression modules, each with its own promoter, CDS and UTRs, have been integrated successfully (Bhat et al. 2025). However, we presently lack the ability in *Auxenochlorella* to simultaneously target multiple loci or both alleles of the same locus in a single transformation – an advantage offered by RNP-mediated approaches. Ideally, both methods would serve complementary roles depending on the application.

Beyond the versatility of genetic manipulation, several other features make *Auxenochlorella* an attractive reference system. Gene families containing multiple members in green algae, and especially in *C. reinhardtii*, more frequently feature just a single gene in *Auxenochlorella* UTEX 250-A. The reduced genetic redundancy is expected to improve the likelihood of achieving informative phenotypes following gene knockout or knockdown, analogous to the lower redundancy of regulatory gene families in the liverwort *Marchantia polymorpha* relative to *A. thaliana* and other angiosperms (Bowman et al. 2017). Although hundreds of genes and gene families have been entirely lost, including many genes that are expected to play roles in photosynthesis, these genes could be re-introduced to the *Auxenochlorella* genome to probe their functions in the species where they do occur. Mutations in genes involved in photosynthesis, such as *CHL27* demonstrated herein, are also possible due to the facultative autotrophy of the species. The metabolic switch of the organism enables chloroplast biogenesis to be studied, and lends itself to the synthetic use of promoters induced in phototrophy or heterotrophy. Genomic manipulation also presents opportunities to address fundamental questions related to genetic mechanisms, as demonstrated by the dissection of the mechanism underlying the translation of bicistronic genes (Dueñas et al. 2025a). *Auxenochlorella* species are highly oleaginous and have GRAS status, and the genomic features reported here could also find utility in bioengineering.

The introduction of artificial chromosomes is a powerful technique that enables multiple traits to be stacked at a single locus (Birchler et al. 2024). If the short AT-rich regions present on *Auxenochlorella* chromosomes can be confirmed to function as centromeres, it may be possible to maintain minichromosomes by introducing an AT-rich region, similar to the synthetic episomes that can be maintained in diatoms (Diner et al. 2017). Alternatively, the aneuploidy of UTEX 250-A and UTEX 25 could be exploited. We demonstrated allele-specific knock-in to a trisomic chromosome, and presumably it would be possible to sequentially manipulate this chromosome with minimal impact on fitness. Finally, since *Auxenochlorella* species are generally transformable, including the hydra symbiont *A. symbiontica* CCAP 211/61, the genetics of host/symbiont establishment and maintenance could be studied (Huss et al. 1994).

Collectively, the yeast-like evolutionary biology and genetic manipulation of *Auxenochlorella*, thus far unique among green algae, presents many exciting prospects for discovery in plant biology and utilization in bioengineering.

## METHODS

### Strains, media and growth conditions

*Auxenochlorella* strains UTEX 250, UTEX 25 and UTEX 2341 were originally obtained from the University of Texas Culture Collection of Algae. A single colony of UTEX 250 was isolated and used in all experimental work (i.e., UTEX 250-A) except for *CHL27* gene targeting, which was carried out in the original UTEX 250 strain. *A. symbiontica* CCAP 211/61 and *Auxenochlorella* CCAP 211/7C were obtained from the Culture Collection of Algae and Protozoa. Strains were typically grown in TAP (for DNA extraction) or HPv1 (for RNA) culture media using the trace element recipe of Kropat et al. (2011). HPv1 is a modified version of TP that replaces 20 mM Tris with 20 mM HEPES. Standard liquid cultures were generally grown under constant light (100 μmol photons m^−2^ sec^−1^) and shaken at 200 rpm. Specific growth conditions were used for preparing RNA from UTEX 250-A for the generation of IsoSeq and RNAseq datasets. These included a range of photoautotrophic, mixotrophic or heterotrophic conditions, different light conditions, CO_2_ levels, and elemental adjustments to media (Dataset S18).

### Extraction and sequencing of nucleic acids

UTEX 250-A DNA was extracted at UC Davis DNA Technologies Core from a frozen cell pellet and used to prepare and sequence PacBio HiFi and OmniC linked-read libraries. Individual IsoSeq libraries were produced from RNA extracted from cultures grown in autotrophic replete and heterotrophic with 2% (w/v) glucose conditions (HPv1 media), and each library was sequenced on a Sequel II SMRT cell at UC Davis. For all remaining conditions (Dataset S18), Illumina RNAseq with poly(A) selection was performed on a Novoseq 6000 platform at UC Davis, generating stranded 150 bp paired-end reads.

UTEX 25, UTEX 2341, CCAP 211/61 and CCAP 211/7C were cultured mixotrophically in TAP medium. High molecular weight DNA was extracted using a modified version of a CTAB and Phenol:Chloroform protocol (Camacho et al. 2025). Cell pellets were ground in liquid nitrogen in a mortar and pestle prior to the CTAB incubation. Size selection was performed using the PacBio short read eliminator (SRE) kit following user instructions.

DNA from UTEX 25, UTEX 2341 and CCAP 211/61 was prepared in a multiplexed library along with five other unrelated samples and sequenced on a single PacBio Sequel II SMRT cell at QB3 Genomics, UC Berkeley. ONT sequencing for CCAP 211/61 and CCAP 211/7C was performed using the Ligation Sequencing Kit V14, two independent R10.1.4 flow cells, and a MinION Mk1B device following user instructions.

### Genome assembly of *Auxenochlorella* UTEX 250-A

The UTEX 250-A PacBio HiFi reads were first subsampled to remove any reads shorter than 18 kb, resulting in ∼0.98 Gb of total sequence. An initial phased diploid assembly was produced by passing the filtered HiFi reads and Omni-C reads to Hifiasm v0.16.1-r375 (Cheng et al. 2022), which was run with default parameters. The initial assembly consisted of two sets of phased contigs, haplotype 1 and 2, featuring 17 and 21 nuclear contigs, respectively.

Several steps of manual assessment and assembly were then performed to achieve a final telomere-to-telomere phased diploid assembly. First, haplotype 1 and haplotype 2 contigs were aligned to each other using minimap2 v2.23-r1117 (Li 2021) with the parameter “-x asm20”. The raw PacBio HiFi reads were also mapped against each haplotype assembly with minimap2 (“-x map-hifi”). Assembly and read-based alignments were manually inspected using IGV v2.16.2 (Robinson et al. 2011). Due to the rearranged nature of the parental haplotypes, we found that the two automated haplotype sets did not individually represent a complete copy of the haploid genome (i.e. some genomic regions were present twice in one haplotype assembly, and absent from the other). We therefore transferred contigs between the two haplotype assemblies to arrive at two sets of contigs that each represented a complete haploid genome, which were arbitrarily labelled haplotype A and B.

Via comparison of the haplotype A and B assemblies it was possible to identify contigs that could potentially be fused i.e. at locations where the ends of two contigs from one haplotype mapped to a single assembled region from the other haplotype. Based on manual inspection of the raw reads, four pairs of contigs were successfully fused. This process involved either removing redundant sequence from one of the contig ends, or adding additional sequence that filled the gap between the two contigs. Where gap filling was required, haplotype-specific raw reads spanning the gap were extracted, aligned with MAFFT v7.490 (Katoh and Standley 2013), and reduced to a single consensus sequence that was subsequently trimmed and inserted between the two contigs. Following a similar approach, the termini of several chromosomes were manually extended to reach the telomeric repeats. Uniquely mapped reads that exhibited soft-clipped bases extending beyond the assembled chromosome were extracted, aligned, and reduced to a consensus sequence that was trimmed and appended to the chromosome. These manual steps resulted in the A and B haplotypes each being represented by 12 gapless chromosomes that either terminated in telomeric repeats or rDNA arrays.

Two additional contigs represented the third copies of the trisomic chromosomes, C03 and C05. The contig corresponding to chromosome C05 was manually truncated at the location of the internal telomere (see Figure S7), since the original contig was misassembled as an exact duplicate of chromosome B05. Finally, eight short contigs were removed from the assembly; six contigs entirely featured rDNA that is already represented as truncated arrays on the assembled chromosomes, and two contigs were identified as duplicate redundant sequences that are similarly represented in the chromosomal assembly. The assembly and phasing of the entire final assembly was manually verified by examining read alignments to each haplotype.

The plastome and mitogenome were each identified as single linear contigs in the Hifiasm assembly. Circularization was performed manually by identifying and removing redundant sequence from one end of each of the organellar contigs.

### Oxford Nanopore basecalling and methylation analyses

ONT raw reads derived from CCAP 211/61 were basecalled using Dorado v0.3.2 (https://github.com/nanoporetech/dorado), which was run in super accuracy basecalling mode using the command “basecaller” and the model “dna_r10.4.1_e8.2_400bps_sup@v4.2.0”. ONT reads for CCAP 211/7C were basecalled using Dorado v0.6.0 using the options “sup,5mCG_5hmCG” or “sup,6mA” to perform basecalling in super accuracy mode with the detection of 5mC and 5-hydroxymethylcytosine (5hmC) calls at CpG dinucleotides, or 6mA calls, respectively. The resulting BAM files were converted to FASTQ using samtools 1.18-1 (Danecek et al. 2021) using the “-T ‘*’” option to retain base modifications. The resulting FASTQ files were mapped against the UTEX 250-A assembly using minimap2 with recommended parameters for ONT 10.4.1 chemistry (“-x map-ont -k19 -w19 -U50,500 -g10k -a -y”). The 5mC status of each CpG site was then called using modkit v0.2.7 (https://github.com/nanoporetech/modkit) with the command “pileup --preset traditional”, which aggregates methylation calls on both strands and returns only 5mC information. The 6mA status of each ApT site was similarly called using the command “pileup --motif AT 0”. Sites with less than 5 mapped reads were removed. For each site, the fraction of modified bases at each site was then extracted, and the median for each nucleotide position relative to the TSS was calculated across all genes that were predicted using IsoSeq data (see below).

### Genome assembly of Auxenochlorella protothecoides UTEX 25 and UTEX 2341, and Auxenochlorella symbiontica CCAP 211/61

The UTEX 25, UTEX 2341 and CCAP 211/61 genome assemblies were produced using Hifiasm v0.19.8-r603. For UTEX 25 and UTEX 2341, all PacBio HiFi reads were passed to Hifiasm, which was run with default parameters. For CCAP 211/61, PacBio HiFi reads were supplemented with ONT reads, which were passed to Hifiasm with the flag “--ul”.

When run without linked-read data (e.g. Omni-C), Hifiasm produces a primary haploid assembly, as well as a diploid assembly consisting of two partially phased haplotypes. For each strain, we first mapped the two (pseudo)haplotypes of the diploid assembly to each other using minimap2 (“-x asm20”) and manually inspected the alignments to confirm that all contigs exhibited a one-to-one syntenic relationship between the two haplotypes (i.e. they are homologous without interchromosomal rearrangements as observed in UTEX 250-A). We then produced a representative haploid assembly by mapping the primary contigs to the UTEX 250-A genome assembly with minimap2 (“-x asm20”) and manually inspecting alignments. For CCAP 211/61, 14 primary contigs corresponded to 12 nuclear chromosomes and two circular organellar genomes. For both UTEX 25 and UTEX 2341, one pair of nuclear contigs and one pair of plastome contigs were each fused to produce the final assemblies, with all other major contigs corresponding to near-complete chromosomes. Contig fusion and circularization of the organellar contigs was performed using the manual assembly approaches described above. As with the UTEX 250-A assembly, additional short contigs featuring rDNA or redundant sequence were removed.

### Quantification of heterozygosity and divergence among strains

To quantify heterozygosity (i.e. individual-level genetic diversity) between the A and B haplotypes of the UTEX 250-A assembly, the haplotype A and B chromosomes were aligned against each other using minimap2 (“-x asm20”). Variants segregating between the A and B haplotypes were then identified using the minimap2 script paftools.js and the parameters “call -l 2000 -L 10000” (i.e., a minimum alignment length to compute coverage of 2 kb, and a minimum alignment length to call variants of 10 kb). Chromosomes were divided in to 10 kb windows, and heterozygosity was calculated based on the number of single nucleotide variants (mapping quality of 60) relative to the number of alignable sites per window. Windows with <2000 aligned sites were excluded. Zero-fold, two-fold and four-fold degenerate sites were extracted from the coding sequence of the primary transcript of each A haplotype gene model using the degenotate tool (https://github.com/harvardinformatics/degenotate). All overall heterozygosity values were reported after removing homozygous genomic tracts that are putatively the product of LOH (see below).

Since the UTEX 25, UTEX 2341, and CCAP 211/61 genomes were assembled to the haploid-level, a different approach was used to quantify heterozygosity. Raw PacBio HiFi reads for each strain were mapped against the A haplotype of the UTEX 250-A assembly using minimap2 (“-x map-hifi”). DeepVariant v1.6.0 (Poplin et al. 2018) was then used to call variants directly from the PacBio HiFi read alignments for each strain independently, using the parameter “model_type PACBIO”. Heterozygosity was calculated in 10 kb windows by identifying heterozygous single nucleotide variants segregating among the reads of the strain in question. Callable sites for each strain were determined by mapping the haploid genome assembly of the relevant strain against the A haplotype of UTEX 250 with minimap2 (“-x asm20”) and identifying alignment blocks with paftools.js (“call -l 2000 -L 10000”). Genetic divergence for a given strain was calculated in 10 kb windows as the proportion of variant sites relative to the UTEX 250-A haplotype A. Divergence was averaged over both haplotypes for each strain (i.e., a site where only one of the two haplotypes varied relative to the UTEX 250-A reference contributed half as much to divergence as a site where both haplotypes varied). Divergence was also calculated relative to the UTEX 250 B haplotype using the same approach.

Neighbor joining trees for nuclear genome, plastome and mitogenome sites were produced using MEGA v11.0.13 (Tamura et al. 2021) with the Tamura Nei substitution model and default parameters. For the organelles, either the plastome or mitogenome assembly of each strain was aligned to the UTEX 250-A plastome or mitogenome assembly with minimap2 (“-x asm20”), and variant and invariant sites were called with paftools.js (“call -l 2000 -L 10000”). To be included in the nuclear genome analysis, a site had to meet two criteria, i) to be on a UTEX 250-A chromosome that had not undergone any reciprocal crossovers, and ii) to be in a region that was not affected by a LOH event in any of the strains. These criteria were enforced to ensure that ancestral evolutionary relationships among haplotypes were recovered, and resulted in the analysis of 651,399 sites on chromosome A11. Variants were extracted from either the paftools.js analysis (for variants segregating between the UTEX 250 A and B haplotypes) or the DeepVariant analyses (for variants segregating within and between the other strains). Variants in UTEX 25, UTEX 2341 and CCAP 211/61 were phased relative to their respective genome assemblies (note that these may represent pseudohaplotypes with a small number of phase changes per chromosome).

### Loss-of-heterozygosity

LOH events in the UTEX 250-A genome were called from the alignment of the A and B haplotype assemblies and paftools.js analysis (see above). LOH events were arbitrarily assigned to regions that featured no heterozygous sites over a stretch of 1 kb or more. Events were designated as terminal if they extended to a chromosome end, or interstitial if they fell within a chromosome.

LOH events were similarly called as invariant regions of at least 1 kb for UTEX 25, UTEX 2341 and CCAP 211/61 based on heterozygous variants identified from the DeepVariant analysis.

CCAP 211/7C heterozygous sites and LOH events were also identified using DeepVariant, ran with the parameter “model_type ONT_R104” and supplied with an alignment of the raw ONT reads against the UTEX 250-A haplotype A assembly (minimap2 “-x map-ont”). These alignments were manually inspected to confirm that the karyotype of UTEX 250-A and CCAP 211/7C are identical.

### Aneuploidy

The raw reads for all available strains were first mapped against the relevant genome assembly to calculate coverage. For PacBio datasets, the UTEX 250-A reads were mapped against the phased UTEX 250-A assembly, whereas UTEX 25, UTEX 2341 and CCAP 211/61 reads were mapped against the representative haplotype from their respective haploid assemblies, all using minimap2 (“-x map-hifi”). CCAP 211/7C ONT reads were mapped against the UTEX 250-A assembly using minimap2 (“-x map-ont”), and the “0710” reads were mapped against the UTEX 25 assembly using bwa mem (Li 2013). The coverage at each nucleotide was calculated using the samtools “depth” command, and the average coverage per 20-kb sliding window was calculated.

### *Auxenochlorella* UTEX 250-A gene model annotation

Gene models were annotated using a hybrid approach that utilized both the IsoSeq and RNAseq datasets, followed by extensive manual curation. The A and B haplotypes of the UTEX 250 assembly were annotated independently, except for the entirely homozygous chromosomes (04, 09 and 10).

IsoSeq reads were pre-processed using the IsoSeq3 pipeline v3.8.1 (https://github.com/PacificBiosciences/IsoSeq). Primers and poly(A) tails were first removed from circular consensus sequencing (CCS) reads. The resulting reads from the two libraries were then combined and mapped against the UTEX 250-A genome assembly using minimap2 (“-x splice:hq”). These read mappings were used to partition the reads into sets that either primarily mapped to the A haplotype (≥20 mapping quality), primarily mapped to the B haplotype, or mapped to neither with high quality (<20 mapping quality; this set was enriched for reads derived from homozygous genomic regions). Read clustering was then performed individually on each read set (“isoseq3 cluster –use-qvs”), to avoid clustering reads that derived from different alleles of the same gene. The three clustered read sets were then re-combined and aligned collectively to the UTEX 250-A assembly using pbmm2 v1.9.0 (“--preset ISOSEQ”).

Unique isoforms were called using the isoseq3 collapse command with the options “--do-not-collapse-extra-5exons --max-5p-diff 20 --max-3p-diff 20”. These options were selected following manual assessment of the quality of the IsoSeq data and effectively enabled alternative TSSs (within a different exon of a longer isoform, or >20 bp away if on the same exon) and termination sites (>20 bp if on the same exon) to be preserved as independent isoforms.

Gene models were predicted from the isoseq3 output using SQANTI3 v5.1 in QC mode with default parameters (Pardo-Palacios et al. 2024), which utilizes GeneMarkS-T (Tang et al. 2015) to predict ORFs from isoforms. The isoseq3 pipeline groups neighboring genes under a single gene model if any of their isoforms overlap on the same strand, which is a frequent occurrence in compact genomes such as that of *Auxenochlorella*. We thus had to “decouple” the isoforms of incorrectly merged genes based on comparison of the protein-coding coordinates of each isoform (i.e. isoforms which did not share coding sequence were separated to independent gene models). Next, we removed any isoforms that were supported by <5 reads or that were derived from <5% of the total reads associated with a given gene. This step minimized the number of false isoforms derived from 5’ truncated reads. Finally, the primary isoform for each gene was selected based on a combination of read abundance and length criteria. For the majority of genes with multiple isoforms, the isoform with the longest coding sequence was also the most abundant and was deemed the primary isoform (where multiple isoforms had the same coding length but differed in their UTRs, the most abundant was selected). For genes where the longest coding sequence was not the most abundant, it was deemed the primary isoform if it was supported by ≥50% of the reads associated with the most abundant isoform. If this was not the case, the most abundant isoform was selected if its coding length was ≥80% of the isoform with the longest coding length. In all other cases, the longest isoform was selected to avoid overly short isoforms being selected. since they may be overrepresented in IsoSeq data.

We noticed that many ORFs were truncated due to predicted initiation at an internal ATG codon. To perform ORF extension, we first built a model of the Kozak sequence surrounding annotated initiation sites from primary transcripts that had no in-frame upstream ATG codons (i.e. ORFs that could not be extended), based on the assumption that most of these annotations represented true initiation sites. The strength of the Kozak sequence surrounding in-frame upstream ATG codons for the remaining transcripts was then calculated by comparison to this Kozak model (see Cross (2015) and Dueñas et al. (2025a) for details). ORF extension was performed if an upstream in-frame start codon existed with a Kozak score greater than the 25^th^ percentile of the scores from which the Kozak model was built, or if the upstream Kozak score was simply higher than that of the annotated start codon.

In parallel to the above annotation, we used high confidence gene models derived from the photoautotrophic IsoSeq dataset to train the AUGUSTUS v3.3.2 gene predictor tool (Stanke et al. 2008), following the protocols described by Hoff and Stanke (2019). Pre-processing and gene model annotation using isoseq3 and SQANTI3 was performed as above. The training set of gene models was produced by extracting the most abundant isoform supported by a minimum of 10 reads from gene models derived from IsoSeq reads that mapped uniquely to the A haplotype. Accurate AUGUSTUS training requires information on the noncoding DNA flanking the training set genes. To avoid including unannotated genes in the flanking regions, we performed a preliminary de novo gene annotation using BRAKER v2.1.2 (Hoff et al. 2016). Each RNAseq dataset was mapped against the UTEX 250-A assembly using STAR v2.7.10a (“--twopassMode Basic --alignIntronMax 5000”) (Dobin et al. 2013), and subsequently passed to the braker.pl script using default parameters. We then extracted BRAKER gene models that had no overlap with the IsoSeq-derived training gene set using bedtools v2.30.0 (Quinlan and Hall 2010). The combined dataset of IsoSeq training genes and nonoverlapping BRAKER genes was then used to compute the flanking region length (1,366 bp), and the final training set was produced by extracting the training set genes with flanking DNA. This dataset was then split randomly into a gene set for training (N=3,444 genes) and for testing the trained parameters (N=1,000). Training was performed using both the standard hidden-Markov model and the conditional random field (CRF) approach, which can result in more accurate gene prediction but is less robust to errors in the training dataset. We selected the CRF parameters based on increased sensitivity when applied to the test dataset. A final RNAseq-based gene model prediction was then performed by passing the RNAseq BAM files (see above) to BRAKER using the trained CRF parameters (“— useexisting”).

The IsoSeq and AUGUSTUS-based gene models were then semi-manually integrated. IsoSeq-based models were given precedence, and an AUGUSTUS model was automatically retained if it did not share any same-strand coding exons with an IsoSeq model. AUGUSTUS models that exactly matched an IsoSeq model (with the exception of the first and/or last exon, which was often incorrectly predicted by AUGUSTUS) were immediately filtered. All other AUGUSTUS models, which could have partial overlap with an IsoSeq model, were manually curated using all IsoSeq and RNAseq evidence. Alternative transcripts were analyzed by first removing isoforms that differ only in their UTR sequences, before passing the resulting annotation to SUPPA v2.4 (Trincado et al. 2018) run with the command “-f ioe -e SE RI SS” to classify alternative isoforms into the categories “skipping exon”, “retained intron” and “alternative splice site”.

Annotation of the plastome and mitogenome was performed using the GeSeq webserver (Tillich et al. 2017), which was provided with existing annotations from *A. protothecoides* “0710” and *Prototheca wickerhamii* (Yan et al. 2015). Additional genes were manually annotated by searching for long ORFs in the unannotated sequence.

### Repeat annotation

Preliminary annotation of interspersed repeats was performed by running RepeatModeler 2.0.5 (Flynn et al. 2020) on the UTEX 250-A genome with the parameter “–LTRStruct”. Consensus sequences for specific transposon families were then manually curated following Goubert et al. (2022) (see Dataset S3, Figure S2). The manually curated models were then combined with the automated models and passed to RepeatMasker v4.1.6 (https://github.com/Dfam-consortium/RepeatMasker) (“–gccalc”). Tandem Repeats Finder v4.10.0 was run on the UTEX 250-A genome to identify tandem repeats using the parameters “2 7 7 80 10 50 2000 -f -d -m - ngs”. Total repeat density was calculated by combining the output of RepeatMasker and TRF. Transposon genes (see Table 1, Figure S2) were identified and marked in the GFF3 annotation by intersecting the coordinates of manually curated transposons and gene model coding sequence, followed by manual inspection.

Repeat densities for the genome assemblies in Table 1 were similarly performed by combining the output of RepeatModeler/RepeatMasker and TRF. RepeatModeler was not run for *A. protothecoides* 0710, which was repeat masked using the UTEX 250-A repeat library (see above), nor *C. reinhardtii*, for which an extensive repeat library exists (Craig et al. 2021).

### Gene family and phylogenetic analyses

Gene family analyses were performed using the 19 species shown in Figure 1A. Protein sequences for each species were reduced to their primary isoforms, and where possible genes on the organelle genomes were removed. Annotated transposon proteins were included if known.

For UTEX 250-A, all A haplotype genes were arbitrarily selected and supplemented by genes that are putatively unique to the B haplotype (see below). For *Picochlorum* sp. BPE23, the genes from only haplotype A were arbitrarily used. Metadata for protein datasets are presented in Dataset S19.

OrthoFinder v2.5.5 was run on the 19 protein sets using the Diamond search method (“-S diamond_ultra_sens”) (Emms and Kelly 2019). Orthogroups featuring a known transposon protein were filtered out (the results of which are reflected in Table 1). GreenCut2 and CiliaCut orthologs were extracted for UTEX 250-A and *C. vulgaris* by comparison to *C. reinhardtii* (i.e., from the “orthologues” output directory of OrthoFinder). For the specific genes shown in Figure 5E and F (LHCA, NPQ, meiosis and syngamy), the gene trees for the underlying orthogroups were manually validated.

The species tree in Figure 1A was produced from a concatenated alignment of the 669 single-copy orthologs identified by OrthoFinder. For each ortholog, proteins were aligned using MAFFT v7.525 (“–maxiterate 1000 –localpair”), and the resulting alignments were trimmed using trimAl v1.4.rev22 (“–automated1”) (Capella-Gutiérrez et al. 2009). The trimmed alignments were concatenated and a maximum likelihood phylogeny was produced using IQ-TREE v2.3.0 (“-m MFP -bb 1000”) (Minh et al. 2020). For Figure 1B, rDNA sequences were accessed from NCBI (Dataset S1), aligned with MAFFT (“L-INS-i”), manually trimmed, and passed to IQ-TREE as above.

### Gene IDs and functional annotation

Allelic gene pairs were determined by running SynChro (Drillon et al. 2014), a tool for detecting syntenic orthologs between genomes, on the A and B haplotype genomes and gene models.

Genes were assigned a unique 5-digit number starting with “00005” for the first gene on chromosome A01 (i.e., UTEX250_A01.00005), with each subsequent gene along the A chromosomes receiving a number that increased in increments of five (i.e., UTEX250_A01.00010, UTEX250_A01.00015, etc.). Genes on the B haplotype that had been identified as part of an allelic pair were assigned the gene number of their corresponding A gene, regardless of their order on the B chromosomes. Genes that were determined to be unique to haplotype B were given new gene numbers that extended beyond the last number that was used on haplotype A. Genes on the plastome were given unique numbers starting at “80000” (e.g., UTEX250_xCp.80000), and genes on the mitogenome starting at “90000” (e.g., UTEX250_xMt.90000). Isoforms were distinguished by adding a number to the gene ID, with the primary isoform labelled as “1”, e.g., UTEX250_A01.00030.1.

In addition to IDs, genes were also assigned a unique name or “gene symbol” where available following the nomenclature of *C. reinhardtii* (Craig et al. 2023). Gene symbols were transferred based on the gene orthology relationships between *C. reinhardtii* and *Auxenochlorella* UTEX 250-A determined by OrthoFinder (“Orthologues” output directory). For genes with 1:1 orthology the *C. reinhardtii* symbol was directly transferred to UTEX 250-A. For multicopy gene families in *C. reinhardtii* that are represented by a single gene in UTEX 250-A, either the symbol with the lowest number was used (e.g., *FOX1*/*FOX2* gene pair orthologous to *FOX1* in UTEX 250-A), or if letters are used, they were removed (e.g., *CMT1A*/*CMT1B* gene pair orthologous to *CMT1* in UTEX 250-A). For the reverse situation, letters were appended to the UTEX 250-A gene symbol where possible (e.g., *LPAAT1* orthologous to *LPAAT1A*/*LPAAT1B* gene pair in UTEX 250-A). Genes on the nuclear genome in *C. reinhardtii* but on an organelle genome in UTEX 250-A were changed accordingly (e.g., *CHLI1* orthologous to *chlI* in UTEX 250-A plastome). Several gene symbols were manually curated, including a small number of genes that are absent from *C. reinhardtii* (e.g., the photosystem I subunit gene *psaM*), and all of the histone genes (for which orthology relationships are difficult to determine). All gene symbol relationships between *C. reinhardtii* and UTEX 250-A are presented in Dataset S7.

Protein domains were identified by running InterProScan v5.67-99.0 (“-dp -goterms”).

### Identification of antisense long noncoding RNA genes

LncRNA genes were annotated based on IsoSeq data using two approaches. First, IsoSeq-based gene models that had initially been annotated as protein-coding but had no orthology to proteins from other species, nor a recognized Pfam or CDD domain, were reassigned as lncRNAs if they overlapped coding sequence of another gene that did meet one of these criteria. Second, IsoSeq gene models supported by at least 5 reads for which no ORF had been annotated by SQANTI3 were added. The first category of genes had been assigned gene IDs as described above, and retained these IDs as lncRNAs instead of protein-coding genes. New gene IDs starting from “50000” and increasing in increments of 5 were introduced for the second category of lncRNA genes.

The coordinates of protein-coding genes with either a homolog in at least one other species or a recognized domain were then intersected with the lncRNA gene coordinates, and those with antisense overlap spanning at least 30% of the protein-coding gene length were selected for manual curation. Validated gene models with antisense lncRNAs were then divided into those with internal birdirectional promoters, or those with internal or immediately adjacent promoters without evidence for birdirectional activity. Gene ontology enrichment analysis was performed using topGO (Alexa and Rahnenfuhrer 2023) using the GO term output from InterProScan and the classic Fisher Test option.

### Genetic toolkit

Transformation, fluorescence measurements and microscopy were performed following Dueñas et al. (2025a) and Dueñas et al. (2025b). Full details are provided in Supplementary File S1, the sequence of all primers used in this study can be found in Dataset S20, and the sequences of plasmids as a GenBank file in Supplementary File S2.

## DATA AVAILABILITY

All sequencing data associated with *Auxenochlorella* UTEX 250-A is available from NCBI under the BioSample SAMN45466464. The UTEX 250-A genome and gene annotations are specifically available under the BioProjects PRJNA1195245 (Haplotype A) and PRJNA1195244 (Haplotype B). Haploid genome assemblies and sequencing data for UTEX 25, UTEX 2341, CCAP 211/61 and CCAP 211/7A (sequencing data only) are available from NCBI under the BioProject PRJNA1328465.

## ACCESSION NUMBERS

NCBI accession numbers for all *Auxenochlorella* UTEX 250-A genes named in the main text are available in Dataset S21.

## ACKNOWLEDGMENTS

Confocal fluorescence microscopy was performed at the CRL Molecular Imaging Center, RRID:SCR_017852.

## AUTHOR CONTRIBUTIONS

RJC, JLM and SSM designed the research. RJC, MAD, DLC, SDG, MCA-M, Y-TL and JLM performed the research. RJC, MAD, SDG, MCA-M, CLB-H and JLM analyzed the data. RJC, MAD, JLM and SSM wrote the paper.

