## Supplementary File S1 for "Targeted genetic manipulation and yeast-like evolutionary genomics in the green alga *Auxenochlorella*"

### Plasmid construct assembly, transformation and DNA preparation

Cloning workflow was adapted from that of Dueñas et al. (2025a) and Dueñas et al. (2025b). A detailed description of primers and plasmids used in this study can be found in Dataset S20 and Supplementary File S2. Primers for plasmid construct assembly were designed using the NEBuilder® Assembly Tool (https://nebuilder.neb.com/) and synthesized by Integrated DNA Technologies (IDT, San Diego, CA). Genotyping primer design and all *in silico* cloning was conducted using Geneious Prime by Dotmatics (Boston, MA).

PCR was performed using Herculase II Fusion DNA Polymerase (Agilent #600679). Thermocycler conditions were set according to the manufacturer's recommendations based on template and primer concentration, target sequence length, and primer Tm (Dataset S20). PCR products were purified with the Omega Bio-Tek E.Z.N.A.® Gel Extraction Kit. Plasmid contructs were assembled using the NEBuilder® HiFi DNA Assembly mix, transformed into E. cloni® 10G Chemically Competent Cells (LGC Biosearch Technologies, Middlesex, UK) and selected overnight on Luria Broth plates with 200 µg/mL ampicillin (LB-amp) at 37°C. Colonies were cultured overnight in 2 mL of LB-amp media at 37°C with 250 rpm shaking. Plasmids were purified using an E.Z.N.A.® DNA Plasmid Mini Kit I (Omega Bio-Tek Inc. Norcross, GA), and DNA concentrations were measured with a NanoDrop One spectrophotometer (Thermo-Fisher Scientific, Waltham, MA). Correct plasmid construct assembly was validated by Sanger or ONT sequencing (UC Berkeley DNA Sequencing Facility). For scale-up, 1.5 µL of sequenced plasmid miniprep was transformed into 10G competent cells and 200 mL LB-amp cultures were grown overnight at 37°C with 200-250 rpm shaking. Plasmid DNA maxipreps were performed using an E.Z.N.A.® DNA FastFilter Maxi Kit (Omega Bio-Tek Inc. Norcross, GA), and DNA concentrations were measured with a NanoDrop One spectrophotometer.

### Media and growth conditions

ApM1 culture media recipe was adapted from a composition described in US Patents US8927522B2, US12037630B2, and Dueñas et al. (2025a). Salts and compounds were added to 1 L MilliQ-purified water to achieve the following concentrations: 991 mg (7.5 mM) ammonium sulfate (MP Biochemicals 808211) provided by 50 mL of a 150 mM stock solution, 4.2 g (24.1 mM) potassium phosphate dibasic anhydrous (Fisher P288-500), 3.57 g (25.9 mM) sodium phosphate monobasic monohydrate (Fisher S369-1), 240 mg (974 µM) magnesium sulfate heptahydrate (Fisher M63-500), 250 mg (1.3 mM) citric acid (Fisher BP339-500), 30 mg (170 µM) of calcium chloride dihydrate (Fisher C79-500) provided from 1.7 mL of a 100 mM stock solution, 675 µg (2 µM) thiamine-HCl (Sigma T1270-25G) provided from 1 mL of a 2 mM stock solution, and 10 mL of 100X Ap trace element solution. A 1 L stock of 100X Ap trace element solution consisted of 2.743 g (14.3 mM) citric acid (Fisher BP339-500), 11 mg (14.3 μM) copper sulfate pentahydrate (Fisher BP346-500), 330 mg (340.5 μM) boric acid (Sigma B9645-500G), 1.4 g (4.89 mM) zinc sulfate heptahydrate (Sigma Z0251-500G), 948.5 mg (4.79 mM) manganese chloride tetrahydrate (Sigma M3634-100G), 3.9 mg (161 μM) sodium molybdate dihydrate (Sigma M1651-100G), and 110 mg (396 μM) iron sulfate heptahydrate (MP Biochemicals 194663). 5 g/L (0.5% w/v) or 20 g/L (2% w/v) glucose (Fisher D19-212), 20 g/L (2% w/v) sucrose (Fisher S2-500), or 20 g/L (2% w/v) melibiose (Chem-Impex International, Ltd. 00808) were added to media for mixotrophic or heterotrophic growth. 15 g/L (1.5% w/v) agar (Fisher BP1423-500) or 8 g/L (0.8% w/v) agarose (Fisher BP1356-500) was added to make solid media. Ammonium sulfate, thiamine, sugars and antibiotics were added from filter-sterilized stocks after autoclaving and cooling to 50°C to avoid precipitation of salts and degradation of organic compounds.

Tris-acetate-phosphite (TA-Phi) medium for *ptxD* selection was modified to contain 245 mg/L (1.1 mM) of sodium phosphate pentahydrate dibasic (Thermo Scientific Chemicals AC428500010), and 136 mg/L (1.8 mM) of potassium chloride (Fisher P217-500), substituting for potassium phosphate in the standard tris-acetate-phosphate (TAP) medium recipe (Sueoka 1960; Gorman and Levine 1965). Sodium phosphite was replaced with 161 mg/L (1.1 mM) sodium phosphate dibasic (Fisher S374-500) to make TA-Pho medium for positive growth controls. Trace metals were provided according to the revised mineral nutrient supplement described by Kropat et al. (2011).

Strains were grown in temperature-controlled standing (Yamato Scientific America Inc. IC103C, Percival Intellus), or shaking incubators (Innova, New Brunswick Scientific, or HT Multitron Pro, Infors AG). Heterotrophic cultures were grown in the dark, while mixotrophic or photoautotrophic cultures were grown with 40-100 $\mu mol.m^{-2}.s^{-1}$ of white light (cool white fluorescent bulbs or warm white LEDs). Temperatures ranged from 24-28°C, and shaking was between 140-200 rpm.

### ***Auxenochlorella* transformation, selection and genotyping**

Lithium acetate/polyethylene glycol (LiAc/PEG) transformations, heterotrophic selections for thiamine prototrophy, G418 resistance and sucrose hydrolysis, and subsequent genotyping of transformants were performed as described in US Patent US12037630B2 and Dueñas et al. (2025a). A detailed step-by-step protocol for preparing linear DNA, LiAc/PEG transformation, plating and growth of colonies on selectable media, and single colony purification is described by Dueñas et al. (2025b). For antibiotic selections (100 μg/mL G418 or 300 μg/mL hygromycin B), after overnight incubation with linear DNA, LiAC and PEG, transformed cells were recovered for 8 h without selection in 10 mL of ApM1, 2% glucose, 2 μM thiamine media to allow accumulation of the enzymes conferring antibiotic resistance before plating. Selection for thiamine prototrophy was carried out on plates made with 0.8% agarose and no thiamine supplementation to minimize thiamine or thiamine precursor contamination. Cells for phosphite selection were grown mixotrophically in TA-Pho medium with 2 μM thiamine at 26°C, 150-200 rpm shaking, 50-100 μmol.$m^{-2}$.$s^{-1}$ white light (cool white fluorescent bulbs) to $OD_{750}$ between 1.5 and 3. LiAc/PEG transformations were carried out as for heterotrophic cells, transformants were plated on TA-Phi, and phosphite-utilizing colonies were selected at 28°C, with 100 µumol.$m^{-2}$.$s^{-1}$ white light (cool white fluorescent bulbs). Phosphite selection was not successful when starter cultures for transformation were grown in ApM1, presumably due to luxury Pi uptake and increased internal Pi stores for cells grown with 50 mM Pi in ApM1 compared to 1.1 mM in TA-Pho.

Single colonies were purified by plate streaking for cell autonomous selections (G418 resistance, hygromycin B resistance, phosphite utilization), and by serial dilutions for non-cell autonomous selections (thiamine prototrophy, sucrose hydrolysis, melibiose hydrolysis) as described by Dueñas et al. (2025b).

For genotyping, DNA from wild type and transformant strains was extracted using a CTAB method adapted from European Patent EP2785835A2 (Somanchi & Bhat, 2014). 2 mL of stationary phase cultures in screw-capped tubes were centrifuged at maximum speed for 5 minutes, the supernatant was discarded, and cell pellets were frozen at -20°C with a 4 mm glass bead (Fisher 11-312B) for at least 24 h. Next, 300 µL grinding buffer, 1.5 µL RNase A (VWR E866-1ML), and 250 mg of 0.5 mm glass beads (Fisher 35-535) were added to each cell pellet, and the tubes were shaken for 5 minutes in a Mini-Beadbeater-24 (BioSpec Products, Inc. Bartlesville, OK) to lyse the cells. 40 µL of 5M NaCl (VWR E529-500ML) and 66 µL 5% CTAB (VWR 0833-500G) were added to the lysates, mixed by inversion, and incubated at 65°C for 1 hour. After incubation, samples were centrifuged at maximum speed for 10 minutes, and supernatants were transferred to 1.5 mL Eppendorf tubes. Lysates were extracted with 300 µL of 25:24:1 phenol-chloroform-isoamyl alcohol pH 8 (Fisher BP17521-400) and centrifuged at maximum speed for 10 minutes. The aqueous phases were transferred to new 1.5 mL tubes, mixed with 210 µL of isopropanol (0.7 volumes), and incubated at room temperature for 1 hour or at 4°C overnight to precipitate the DNA. Genomic DNA was pelleted by centrifugation at maximum speed for 30 minutes, then the pellets were washed twice with 500 µL 70% ethanol and dissolved in elution buffer (10 mM Tris-HCl. pH 7.5). DNA concentrations were measured with a NanoDrop One spectrophotometer and samples were diluted to 100-200 ng/µL for use as PCR amplification templates. PCR amplifications were performed using Herculase II Fusion DNA Polymerase (Agilent 600679) with reaction buffer, dNTPs, enzyme, and primer concentrations according to the manufacturer's recommendations. 5% DMSO was included in reactions to improve amplification of GC-rich *Auxenochlorella* sequences. Primers used for genotyping are listed in Dataset S20.

### Fluorescence quantification and confocal microscopy

1 mL photoautotrophic and heterotrophic cultures of *Auxenochlorella* were grown in 24 well plates for 3 days to achieve log phase, at which point $OD_{750}$ was measured for normalization with a Spectramax iD3 (Molecular Devices, LLC., San Jose, CA) well plate reader, which was also used for subsequent fluorescence measurements. Before taking a measurement, cell plates were set to

shake at a speed of “high” for 5 seconds. Venus fluorescence was measured using excitation-emission wavelengths of 515/555 nm. Three biological replicates each of the UTEX 250-A wild-type negative control, and three *amt1B-1::Venus* (MAD15) transformant strains were utilized for fluorescence quantification and confocal microscopy. Triplicate technical measurements were taken of each biological replicate and averaged. Error is reported as the standard deviation of the fluorescence intensity of biological replicates for each strain. 50 µL aliquots of each culture were sampled for confocal microscopy and centrifuged at 10 000 x *g* for 1 minute. The cells were then resuspended in 50 µL of ApM1 media, and 10 µL samples were pipetted onto Fluorescent Antibody Slides (Thermo Scientific Cat. No. 3032), covered with a Microscope Cover Glass (Fisherbrand 12-542A) and sealed with nail polish. Samples were observed under a Zeiss LSM880 Laser Scanning Confocal Microscope with a 63x oil objective lens (numerical aperture 1.15). Venus fluorescence was observed with a laser excitation line at 514 nm (intensity, 20%); emission was collected between 519-599 (gain 650; offset 0). Chl fluorescence was observed with a laser excitation line at 635 nm (intensity, 4%); emission was collected between 647-721 (gain 650; offset 0). Corresponding brightfield images were acquired with a transmitted detection module with a Photon Counting PMT (T-PMT, gain 371; offset 0). Images were processed and color corrected using ImageJ software. The brightness and contrast range for Chl was set between 10 and 200. The brightness and contrast range for Venus was set between 10 and 150. The Brightness and Contrast range for Venus was set between 10 and 220.

Strains expressing GFP were grown in ApM1 medium containing 5 g/L (0.5% w/v) sucrose as a carbon source, with 40-50 $\mu mol.m^{-2}.s^{-1}$ of white light (cool white fluorescent bulbs) at 25 °C for 3-4 days. Cells contained in 50 µl of the cell suspension were recovered by centrifugation at 8 600 x *g* for 3 min at room temperature and resuspended in 50 µl of 10 mM sodium phosphate buffer. About 1 µl of the cell suspension was spotted on an agarose cushion (2% low melting point agarose in 150 mM Sorbitol) previously mounted on a microscope slide and gently covered with a coverslip. Samples were then analyzed with a confocal ZEISS LSM 710 microscope (Molecular Foundry Facilities, Lawrence Berkeley National Laboratory) or confocal ZEISS LSM 880 microscope (Molecular Imaging Center, University of California Berkeley).

**Acknowledgments**

Confocal fluorescence microscopy was performed at the CRL Molecular Imaging Center, RRID:SCR_017852. We thank Holly Aaron, Feather Ives, and Luis Alvarez for their microscopy advice and support.
