## Supplementary Figures for "Targeted genetic manipulation and yeast-like evolutionary genomics in the green alga *Auxenochlorella*"

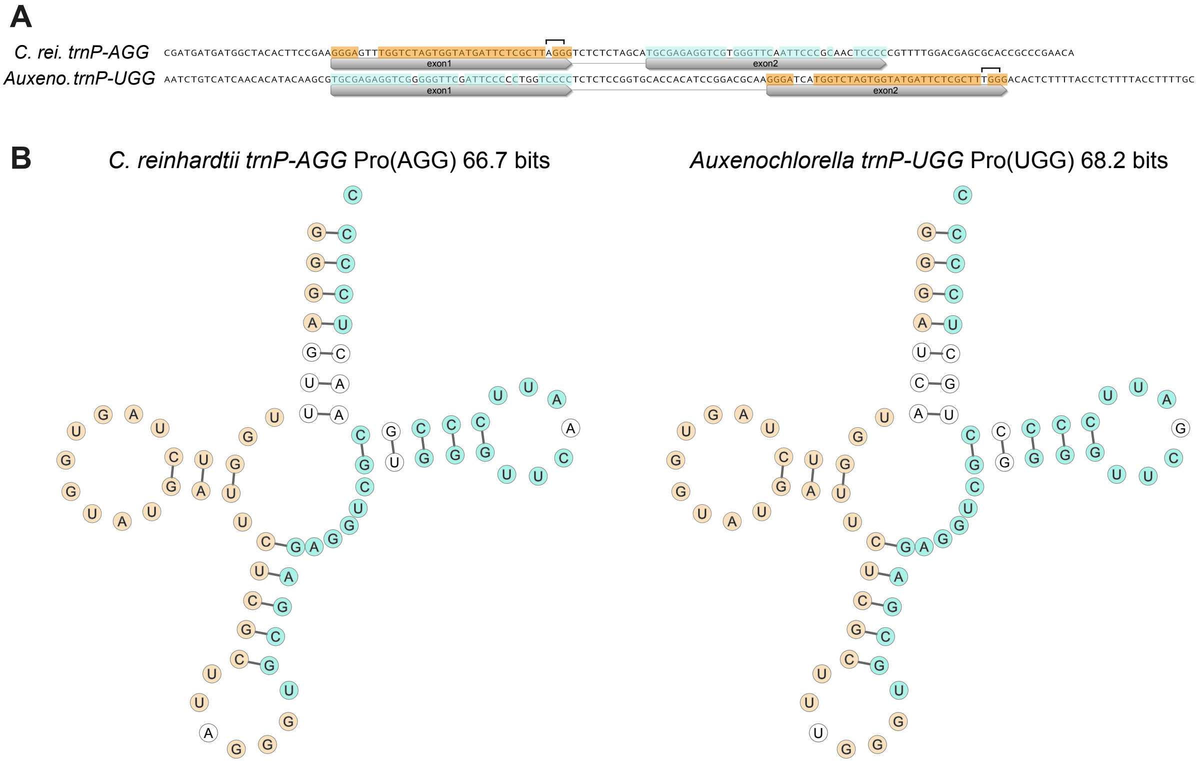


**Figure S1.** Permuted tRNA genes in *Auxenochlorella* UTEX 250-A. A) Gene structure comparison between a *Chlamydomonas reinhardtii* proline tRNA with an AGG anticodon, and an *Auxenochlorella* UTEX 250-A proline tRNA with a UGG anticodon. Orange shading illustrates sequence identity between the first exon of the *C. reinhardtii* tRNA and the second part of the permuted *Auxenochlorella* tRNA. Blue shading illustrates sequence identity between the second exon of the *C. reinhardtii* tRNA and the first part of the *Auxenochlorella* tRNA. The positions of the anticodons are indicated with brackets. B) tRNAScan-SE 2.0 RNA secondary structure predictions of *C. reinhardtii trnP-AGG* and *Auxenochorella trnP-UGG*. **Supports Table 1.**

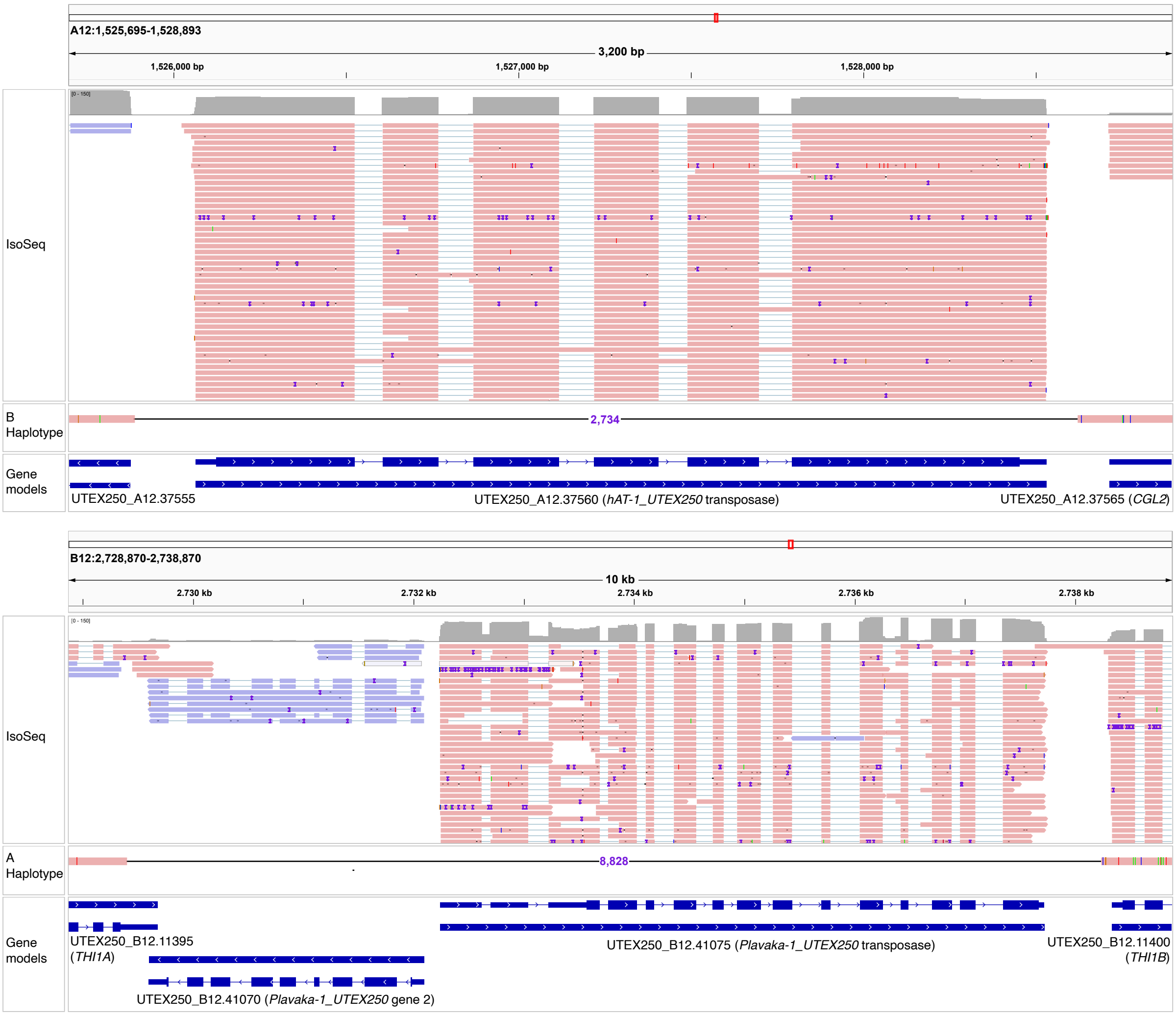


**Figure S2.** IGV screenshots of transposable element insertions that are polymorphic between the A and B haplotypes. Top panel shows a copy of *hAT-1_UTEX250* that is unique to haplotype A, and bottom panel shows a copy of *Plavaka-1_UTEX250* that is unique to haplotype B. Both transposable elements carry expressed genes encoding transposases, as shown by IsoSeq reads. **Supports Table 1.**

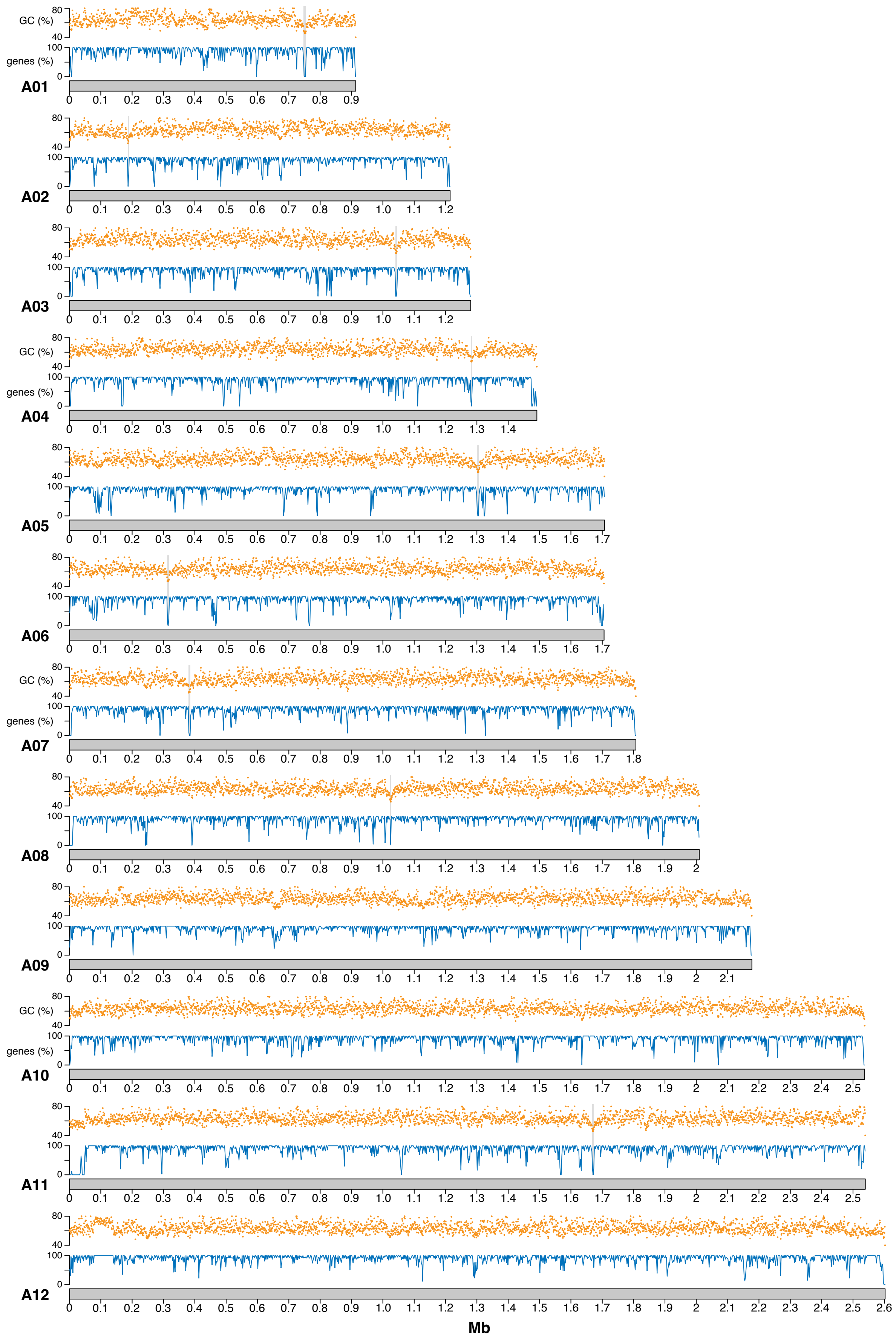


**Figure S3.** Putative AT-rich short regional centromeres in *Auxenochlorella* UTEX 250-A Haplotype A. GC content (1 kb windows) and gene density (2 kb windows) across A haplotype chromosomes. Putative centromeres are marked by grey bars. **Supports Figure 2.**

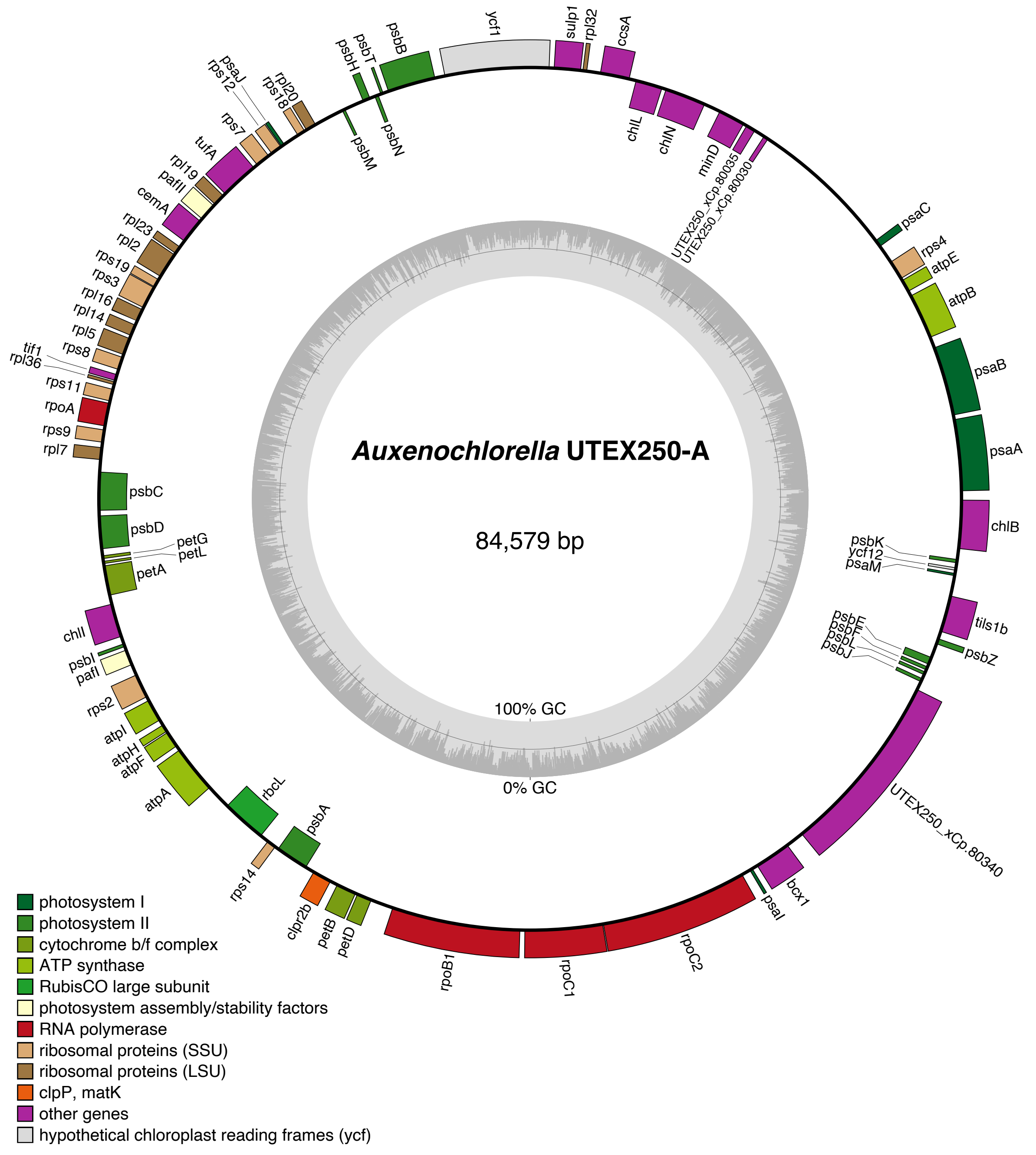


**Figure S4.** Map of *Auxenochlorella* UTEX250-A plastome. Protein-coding genes on the forward strand are shown on the exterior of the map, and genes on the reverse strand on the interior. Inner ring shows GC content. **Supports Table 1.**

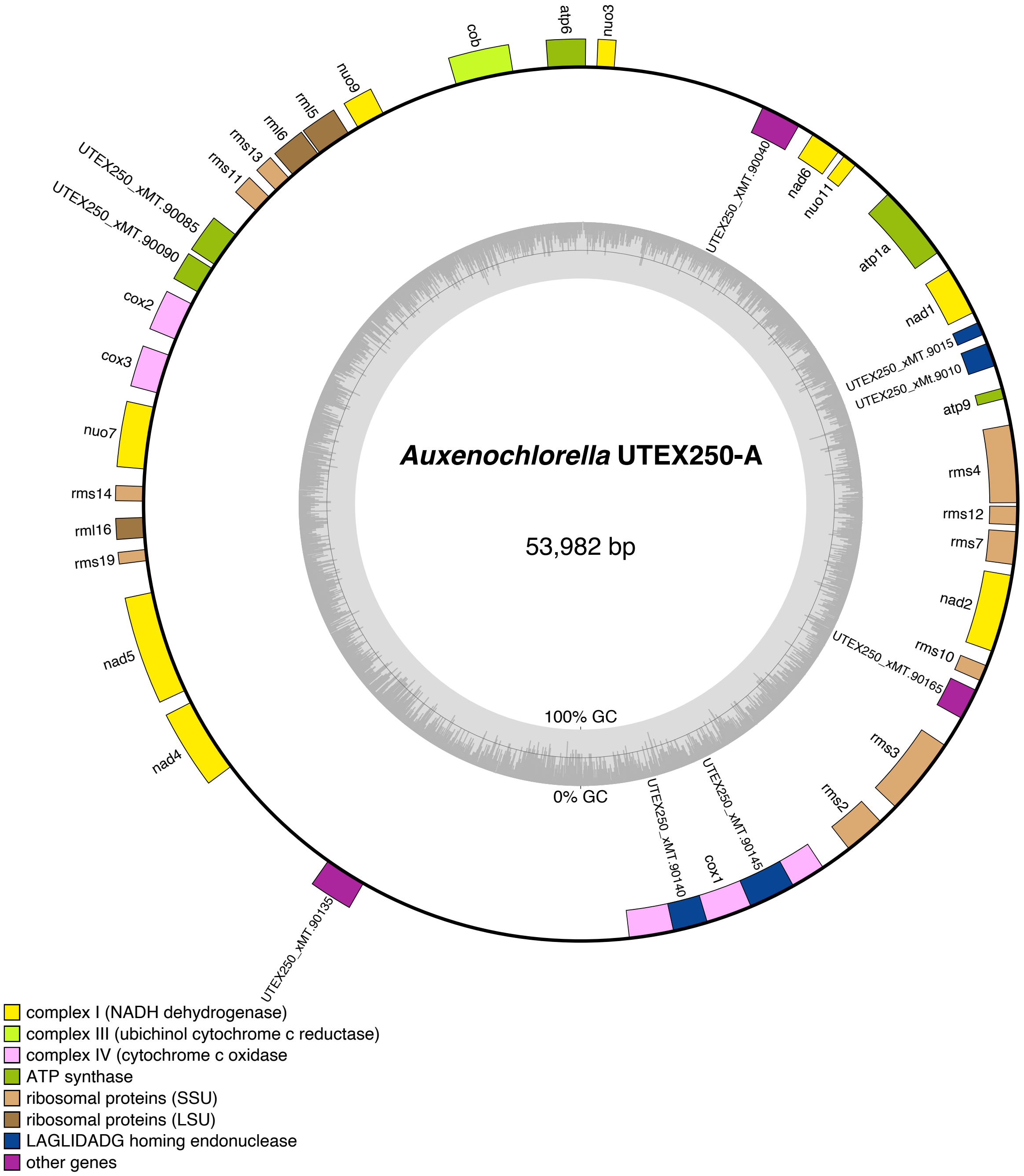


**Figure S5.** Map of *Auxenochlorella* UTEX250-A mitogenome. Protein-coding genes on the forward strand are shown on the exterior of the map, and genes on the reverse strand on the interior. Inner ring shows GC content. **Supports Table 1.**

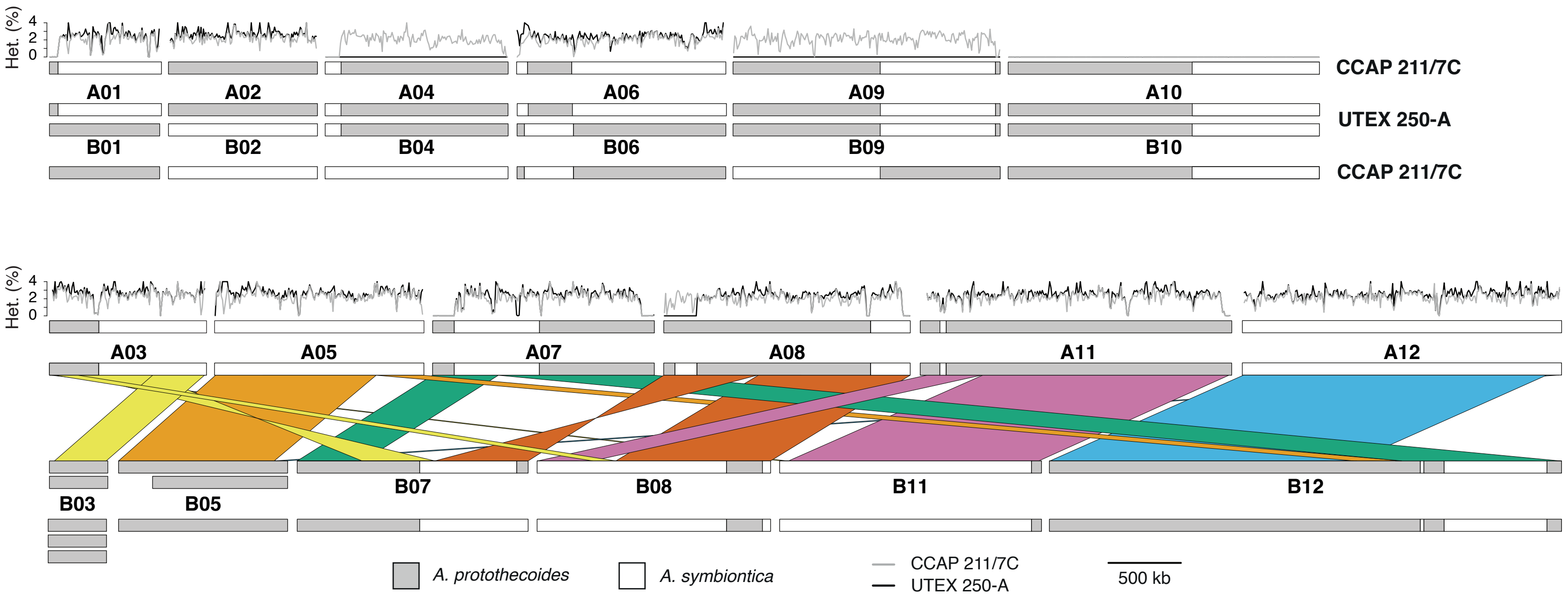


**Figure S6.** UTEX 250-A and CCAP 211/7C chromosome schematics and heterozygosity. The top row shows chromosomes that are homologous, the second row shows rearranged chromosomes with synteny relationships that correspond to the Circos plot in Figure 2B. The central pair of chromosomes correspond to UTEX 250-A, and the top and bottom chromosomes to CCAP 211/7C. Heterozygosity is plotted along each "A" chromosome in 10 kb windows, corresponding to Figure 3B. Aneuploid chromosomes in UTEX 250-A (B03 and B05) and CCAP 211/7C (B03) are represented by extra copies, corresponding to Figure 4A, B. **Supports Figures 3 and 4.**

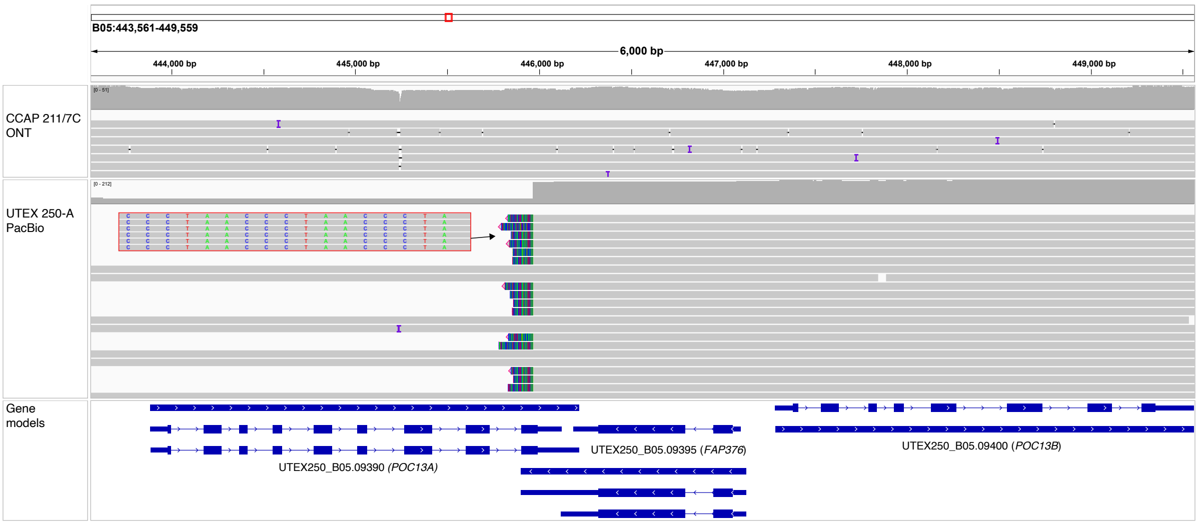


**Figure S7.** IGV screenshot of putative de novo telomere on duplicate fragment of chromosome B05. ONT reads (top panel) from CCAP 211/7C and PacBio reads from UTEX 250-A (middle panel) are mapped against the UTEX 250-A genome assembly. No change in read coverage or soft-clipped reads are observed in the ONT reads, demonstrating that B05 is present in a single copy in CCAP 211/7C. The PacBio reads from UTEX 250-A contain a population of reads that map across the locus and a second population that exhibit soft-clipped bases featuring the telomeric repeat (see zoomed in box showing transition to telomeric repeats), suggesting a second fragmented copy of B05 featuring a de novo telomere. **Supports Figure 4.**

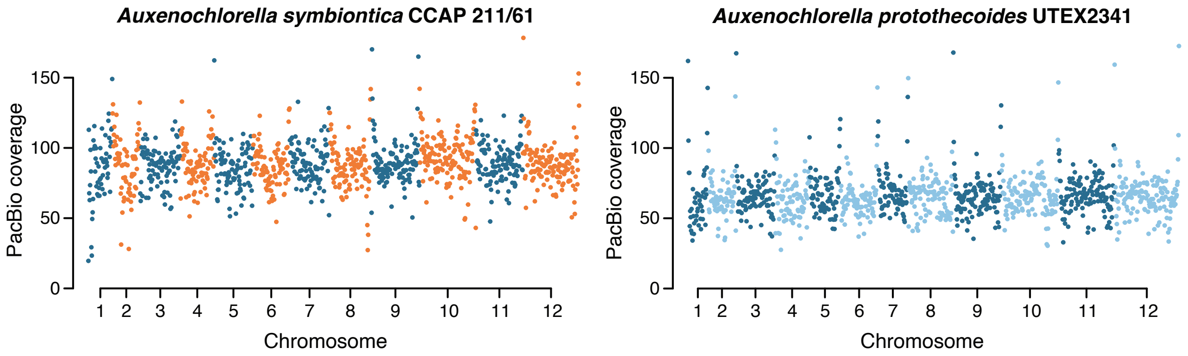


**Figure S8.** PacBio read coverage in 20-kb windows across a representative haplotype of the *Auxenochlorella symbiontica* CCAP 211/61 (left panel) and *Auxenochlorella protothecoides* UTEX 2341 (right panel) genome assemblies. **Supports Figure 4.**

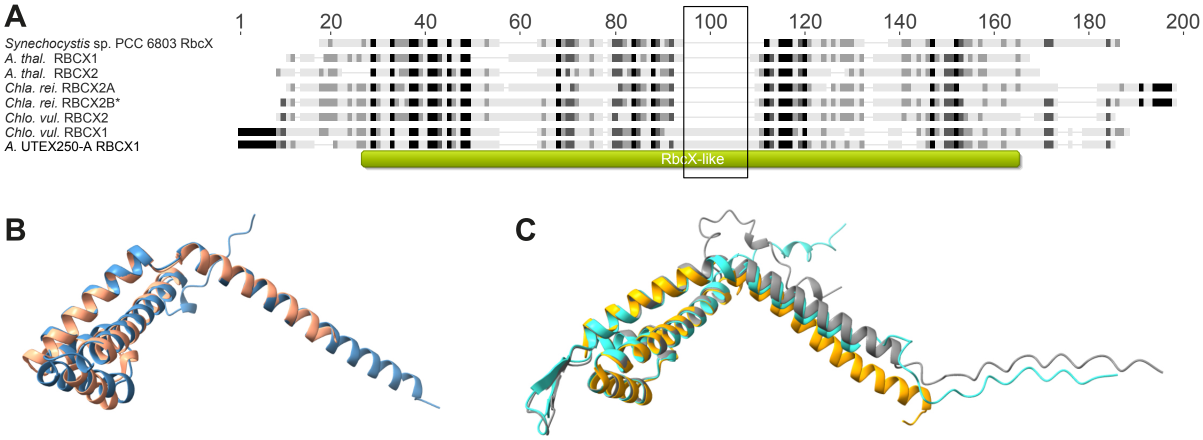


**Figure S9**. Putative RBCX orthologs in *Auxenochlorella* UTEX 250-A and *Chlorella vulgaris*. A) Alignment of RBCX orthologs from *Arabidopsis thaliana*, *C. reinhardtii*, *Synechocystis* sp. PCC 6803, *Chlorella vulgaris* and *Auxenochlorella* UTEX 250-A. Positions with 100% similar amino acids are shaded black, 80-100% similar are shaded dark grey, 60-80% similar are medium grey, and less than 60% similar are not shaded. The green box shows the position of the predicted RbcX-like domain (InterPro ID IPR038052) in UTEX 250-A RBCX1. (*) The Chlamydomonas RBCX2B sequence was truncated to remove a 133 amino acid C-terminus extension. B) Superimposition of the *C. vulgaris* RBCX2 AlphaFold predicted structure (blue) on the *A. thaliana* RBCX1 chain A monomer from PDB accession 4gr2 (salmon) (Kolesinski, et al. 2013). Note that *A. thaliana* RBCX1 is the RBCX-II isoform, and the naming convention in green algae is reversed. C) UTEX 250-A (turquoise) and *C. vulgaris* (grey) RBCX1 AlphaFold predicted structures superimposed on the *Synechocystis* sp. PCC 6803 (orange) RbcX chain A monomer from PDB accession 2py8 (Tanaka, et al. 2007). The boxed region in panel A highlights an insertion unique to the UTEX 250-A and *C. vulgaris RBCX1* orthologs that forms a predicted anti-parallel beta-sheet. **Supports Figure 5.**

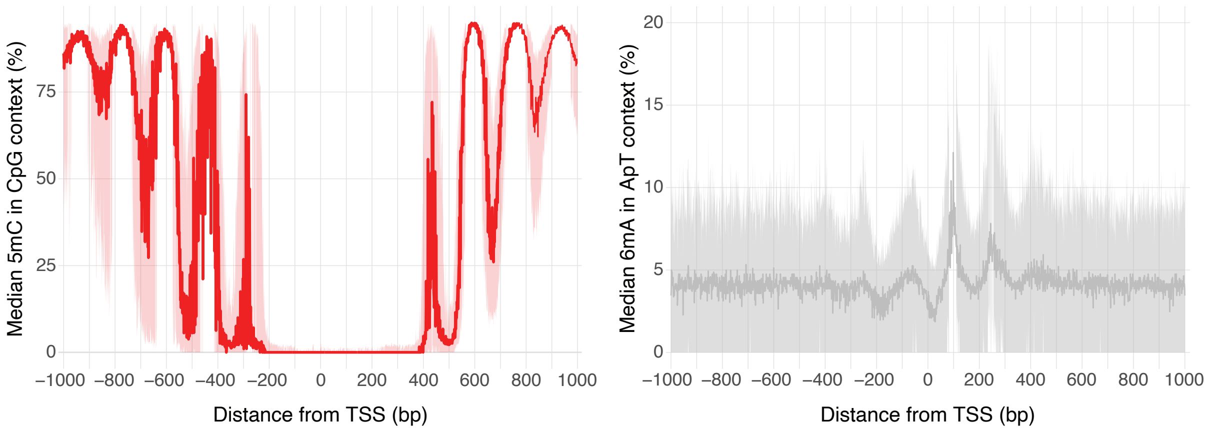


**Figure S10.** 5-methylcytosine (5mC, left panel) and $N^6$-methyladenine (6mA, right panel) around transcription start sites (TSS) of UTEX 250-A genes. Solid line represents the median methylation level across all genes, and shaded area represents second and third quartiles. Methylation was calculated using the ONT reads of CCAP 211/7C. **Supports Figure 6.**

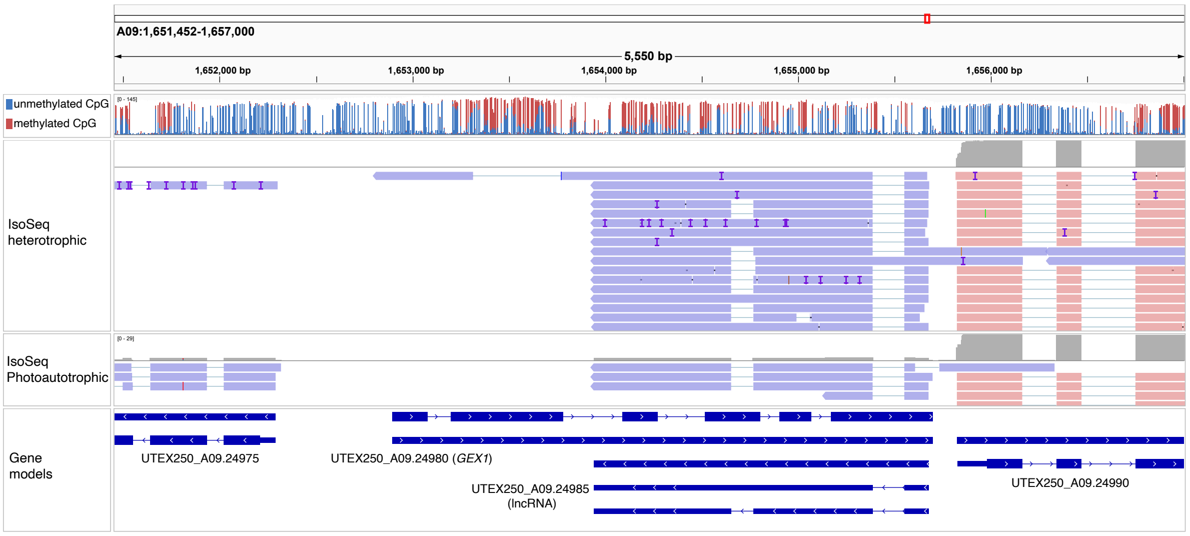


**Figure S11.** IGV screenshot of *GEX1* and antisense lncRNA (UTEX250_A09.24985). Top row shows read coverage of CCAP 211/7C ONT reads where only CpG sites are shown, with the proportion of red to blue corresponding to the proportion of 5mC calls. Second and third rows show IsoSeq reads from heterotrophic or photoautotrophic conditions. Final row shows gene models. **Supports Figure 6.**

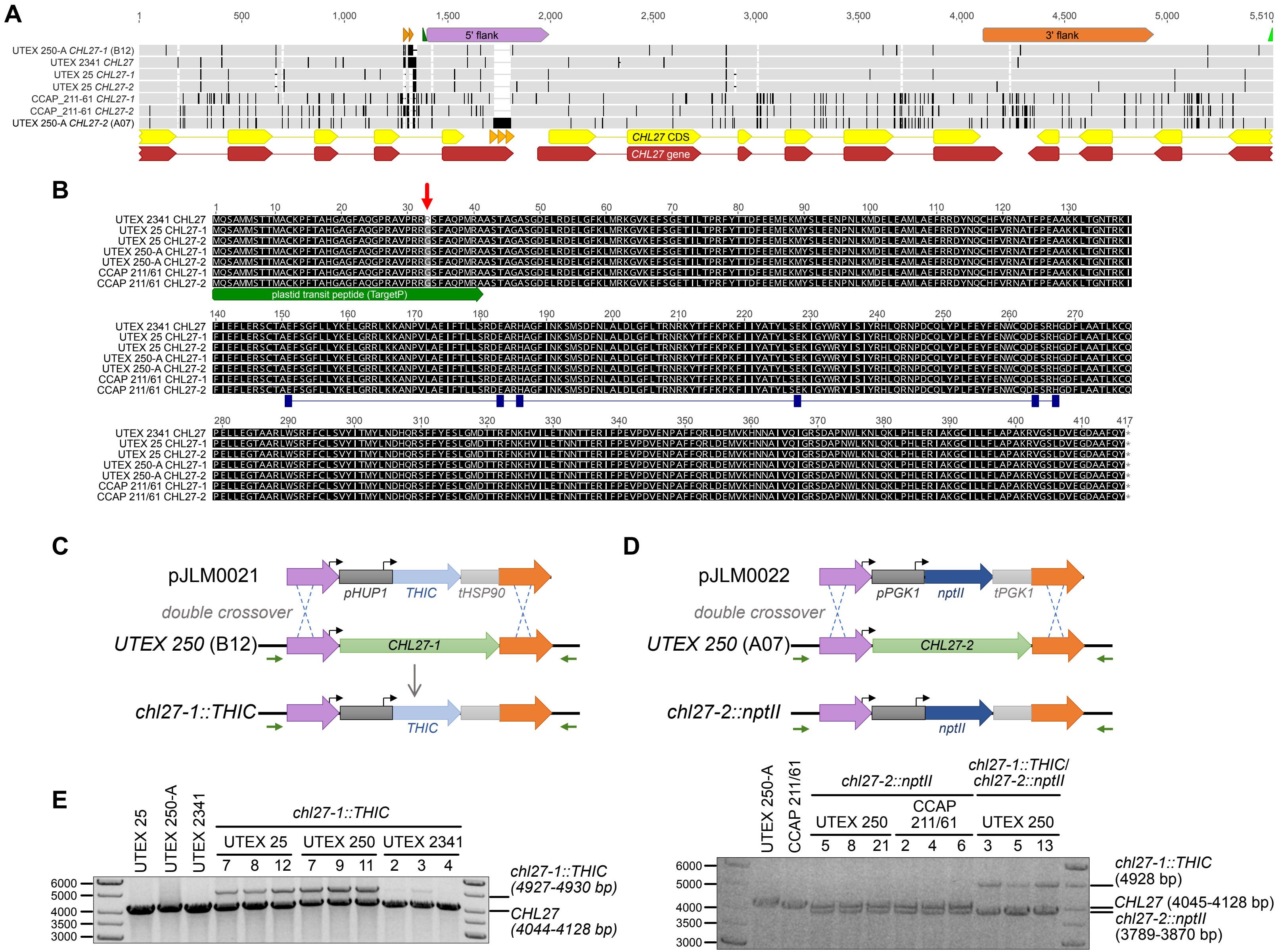
A
1
500
1,000
1,500
2,000
2,500
3,000
3,500
4,000
4,500
5,000
5,510
5' flank
3' flank
UTEX 250-A CHL27-1 (B12)
UTEX 2341 CHL27
UTEX 25 CHL27-1
UTEX 25 CHL27-2
CCAP_211-61 CHL27-1
CCAP_211-61 CHL27-2
UTEX 250-A CHL27-2 (A07)
CHL27 CDS
CHL27 gene
B
UTEX 2341 CHL27
UTEX 25 CHL27-1
UTEX 25 CHL27-2
UTEX 250-A CHL27-1
UTEX 250-A CHL27-2
CCAP 211/61 CHL27-1
CCAP 211/61 CHL27-2
plastid transit peptide (TargetP)
C
pJLM0021
pHUP1
THIC
tHSP90
double crossover
UTEX 250 (B12)
CHL27-1
chl27-1::THIC
THIC
D
pJLM0022
pPGK1
nptII
tPGK1
double crossover
UTEX 250 (A07)
CHL27-2
chl27-2::nptII
nptII
E
UTEX 25
UTEX 250-A
UTEX 2341
chl27-1::THIC
UTEX 25
UTEX 250
UTEX 2341
7
8
12
7
9
11
2
3
4
6000
5000
4000
3500
3000
chl27-1::THIC
(4927-4930 bp)
CHL27
(4044-4128 bp)
UTEX 250-A
CCAP 211/61
chl27-2::nptII
UTEX 250
CCAP 211/61
5
8
21
2
4
6
chl27-1::THIC/
chl27-2::nptII
UTEX 250
3
5
13
6000
5000
4000
3500
3000
chl27-1::THIC
(4928 bp)
CHL27 (4045-4128 bp)
chl27-2::nptII
(3789-3870 bp)

**Figure S12.** *Auxenochlorella* reverse genetics. A) MUSCLE alignment (Edgar 2004) of the *CHL27* loci and flanking regions from UTEX 25, 250-A, 2341 and CCAP 211/61. Grey shading denotes sequence identity and polymorphic nt or regions are shaded black. The 5' and 3' flanking regions targeted by integrating constructs are indicated with orange and magenta boxes. Partial gene models for the upstream and downstream gene and the complete *CHL27* gene model are illustrated below the alignment in red, and the coding sequence models are illustrated in yellow. Orange triangles denote three copies of a 40 bp direct repeat found in the 3' UTR of the gene upstream of UTEX 250-A *CHL27-2* on chromosome A07; this sequence is only present in a single copy upstream of *CHL27-1* and in the other *Auxenochlorella* strains. A GC-rich, variable length direct repeat sequence is found in the last intron of the gene upstream of the *CHL27* locus, illustrated by brown triangles. This region was not resolved in any of the genome assemblies that were generated by short-read sequencing (Gao et al. 2014; Vogler et al. 2018). Green triangles illustrate the positions of primers SM001211 (forward) and SM000018 (reverse) that were used for genotyping by PCR. B) Geneious sequence alignment (Geneious Prime 2025.1.3 (https://www.geneious.com)) of the CHL27 polypeptides translated from the UTEX 25, 250-A, 2341 and CCAP 211/61 alleles. The green arrow indicates the N-terminal plastid transit peptide predicted by TargetP-2.0 (Almagro Armenteros et al. 2019). The position of a G33R amino acid replacement in the UTEX 2341 CHL27 predicted plastid transit peptide is indicated with a red arrow. Blue rectangles illustrate the predicted active site diiron motif ligands. C & D) Schematic representations of targeting constructs for *CHL27* deletion by homologous recombination. pJLM0021, a gene cassette comprised of UTEX 250 *CHL27-1* 5' and 3' flanking sequences, a 745 bp *HUP1* promoter, a codon-optimized, synthetic *Arabidopsis THIC* selectable marker, and a 301 bp terminator sequence containing the 3' UTR of *HSP90*, targets *CHL27-1*. pJLM0022, containing UTEX 250 *CHL27-2* 5' and 3' flanking sequences, a 795 bp *PGK1* promoter, a *nptII* selectable marker, and a 250 bp terminator sequence containing the 3' UTR of *PGK1*, targets *CHL27-2*. Green arrows show the positions of primers SM01211 and SM000018 used to test for transgene integration. E) Genotyping by PCR amplification of *CHL27* loci in wild-type and transformed strains. The expected wild-type PCR amplification product sizes for *CHL27-1* and *CHL27-2* were 4043 and 4044 bp, respectively in UTEX 25, 4044 bp in UTEX 2341, 4045 and 4128 bp, respectively, in UTEX 250-A, and 4048 and 4049 bp, respectively in CCAP 211/61. PCR amplification across the *chl27::THIC* mutant allele in UTEX 25 yielded products of 4927 or 4928 bp, and 4928 bp in UTEX 250 and UTEX 2341. The *chl27::nptII* mutant allele PCR products were 3870 in UTEX 250 and 3789 in CCAP 211/61. **Supports Figure 7.**

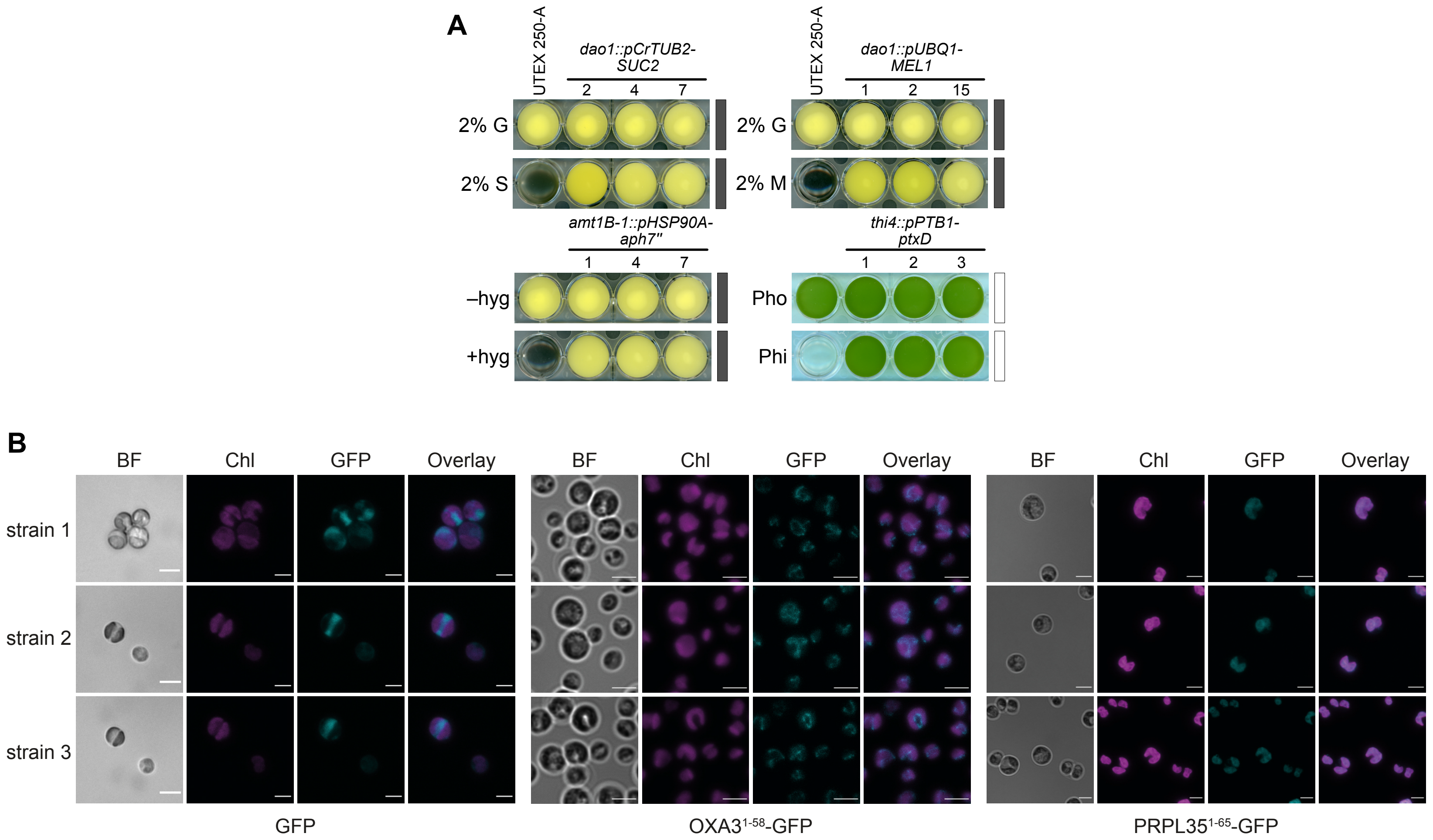

A
UTEX 250-A
dao1::pCrTUB2-SUC2
2
4
7
2% G
2% S
UTEX 250-A
dao1::pUBQ1-MEL1
1
2
15
2% G
2% M
amt1B-1::pHSP90A-aph7''
1
4
7
–hyg
+hyg
thi4::pPTB1-ptxD
1
2
3
Pho
Phi
B
BF
Chl
GFP
Overlay
strain 1
strain 2
strain 3
GFP
OXA3¹⁻⁵⁸-GFP
PRPL35¹⁻⁶⁵-GFP

**Figure S13.** *Auxenochlorella* selection markers and intracellular localization. A) Selectable marker growth phenotypes of transformants compared to UTEX 250-A. Positive growth controls were cultured for four days in the dark (illustrated with black bars) at 26°C with 140 rpm shaking, in 20 g/L glucose without antibiotic selection, or for five days with approximately 40 µmol.m$^{-2}$.s$^{-1}$ of white light (illustrated with white bars) provided by cool white fluorescent bulbs, at 24°C with 140 rpm shaking, in tris-acetate medium containing 1.1 mM phosphate (Pho). Transformants expressing *SUC2* targeted to *DAO1* and controlled by the *C. reinhardtii TUB2* promoter (*dao1::pCrTUB2-SUC2*) grew heterotrophically on sucrose (2% S); *MEL1* integrated at *DAO1* and driven by UTEX 250-A *UBQ1-1* promoter (*dao1::pUBQ1-MEL1*) enabled growth on 20 g/L melibiose (2% M); *aph7"* transformants, targeted to *AMT1B-1* and controlled by the endogenous *HSP90A-1* promoter (*pHSP90A-1*) were resistant to 300 µg/mL hygromycin B (hyg); and *ptxD*, under the control of the *PTB1-1* promoter (*pPTB1*) and integrated at *THI4* allowed mixotrophic growth with 1.1 mM phosphite (Phi) as the sole P-source. All images for transformants (bottom rows) are duplicates of the images represented in main text Figure 7E. B) Fluorescence imaging of UTEX 250-A transformants expressing sucrose invertase and GFP. Cells were from cultures grown with 5 g/L of sucrose for three to four days at 25°C, shaking at 160 rpm, and approximately 40 µmol.m$^{-2}$.s$^{-1}$ of white light provided by cool white fluorescent bulbs. Representative cells from three independent transformants expressing each construct are shown. *GFP* and *targeting peptide-GFP* fusion coding sequences were controlled by the *RBCS1* promoter. BF = bright field. Images for strain 3 for the PRPL35$^{1-65}$-GFP transformant (bottom row) are duplicates of images in main text Figure 7F. **Supports Figure 7.**
